# A syntelog-based pan-genome provides insights into rice domestication and de-domestication

**DOI:** 10.1101/2023.03.17.533115

**Authors:** Wu Dongya, Lingjuan Xie, Yanqing Sun, Yujie Huang, Lei Jia, Chenfeng Dong, Enhui Shen, Chu-Yu Ye, Qian Qian, Longjiang Fan

## Abstract

Asian rice is one of the world’s most widely cultivated crops. Large-scale resequencing analyses have been undertaken to explore the domestication and de-domestication genomic history of Asian rice, but the evolution of rice is still under debate. Here, we construct a syntelog-based rice pan-genome by integrating and merging 74 high-accuracy genomes based on long-read sequencing, encompassing all ecotypes and taxa of *Oryza sativa* and *Oryza rufipogon*. Analyses of syntelog groups illustrate subspecies divergence in gene presence-and-absence and haplotype composition and identify massive genomic regions putatively introgressed from ancient Geng/*japonica* to ancient Xian/*indica* or its wild ancestor, including almost all well-known domestication genes and a 4.5-Mb centromere-spanning block, supporting a single domestication event in rice. Genomic comparisons between weedy and cultivated rice highlight the contribution from wild introgression to the emergence of de-domestication syndromes in weedy rice. This work highlights the significance of inter-taxa introgression in shaping diversification and divergence in rice evolution and provides an exploratory attempt by utilizing the advantages of pan-genomes in evolutionary studies.

## Introduction

As one of the most important calorie sources, Asian rice (*Oryza sativa*), which is considered to be domesticated from its wild progenitor (*Oryza rufipogon*), is widely grown worldwide. Despite its indispensable roles in food supply and fundamental studies about plant biology, the origination, domestication, and subsequent diversification of rice have been under debate for decades, although a considerable amount of archaeological and genetic evidence has been proposed to infer the evolutionary trajectory of rice (Molina et al., 2011; Huang et al., 2012; Civáň et al., 2015; Gross and Zhao, 2014; Choi et al., 2017; Carpentier et al., 2019; Zhang et al., 2021). Multiple taxa or groups in classification, recurrent artificial hybridization during breeding, long-distance dispersal by global trade and other factors have hindered our understanding of rice evolution. The most disputed issue is whether domestication events have happened only once or independently multiple times. Regardless, many domestication- and improvement-related genes have been identified to underlie domestication syndromes, such as plant architecture (*Prog1*), grain shattering (*sh4*), awn length (*An-1* and *LABA1*), pericarp color (*Rc*) and dormancy (*Sdr4*) (Chen et al., 2019). Recently, the issue of rice feralization or de-domestication has attracted great attention in both agricultural production and basic biology because the de-domesticated ecotype of rice (weedy rice, *Oryza sativa* ssp. *spontanea*) has severely threatened rice yield and quality as a commonly seen weed in paddy fields. The genetic resources of weedy rice also show potential for use in enhancing abiotic stress adaptation in rice breeding (Sun et al., 2022). How atavism occurs in rice is an intriguing biological question that has further extended and complicated the evolutionary history of rice (Wu et al., 2021). Previous studies have suggested that the genomes of weedy rice were mostly derived from local cultivated rice by recurrent and independent de-domestication events and that genetic introgression from wild rice may have contributed to weediness (Song et al., 2014; Li et al., 2017; Qiu et al., 2017; Sun et al., 2019; Qiu et al., 2020).

Pan-genomic studies have been conducted in a wide range of crops, including rice (Zhao et al., 2018; Wang et al., 2018b; Qin et al., 2021; Zhou et al., 2020; Zhang et al., 2022; Shang et al., 2022). By comparing *de novo* assemblies, large SVs have been discovered, underlying important traits that could not be explained by small-scale variations. How to utilize pan-genomes in evolutionary studies has been relatively little explored. Here, we integrate high-accuracy rice genomes covering all ecotypes and taxa and high-depth resequenced genomes of wild rice to revisit the origin, domestication, diversification and de-domestication processes of rice based on a syntelog-based pangenome. Whole-genome mosaic haplotype maps intuitively reveal massive introgression footprints of domesticated genomic blocks from the proto-GJ (initial domesticates of subspecies Geng/*japonica*) to XI (subspecies Xian/*indica*) ancestor, strongly supporting the hypothesis of single domestication in rice. Structural variations between weedy and cultivated rice indicate that the introgression events from wild progenitors to different cultivated rice groups probably underlie the parallel convergence in weedy traits (e.g. pericarp and hull color). Briefly, our study comprehensively investigates the complex relationship among different rice taxa and ecotypes from a pan-genome view and highlights the significance of genomic introgression in both rice domestication and de-domestication.

## Results

### High-quality rice genome assemblies

To fully capture the genomic diversity and dynamics in rice domestication, improvement and further feralization, we created a panel of high-quality rice genome assemblies, including newly generated assemblies of 11 weedy and one cultivated accession, using PacBio HiFi mode with an average sequencing depth of 32.4×. Contigs were anchored on chromosomes using a reference-guided approach, and Hi-C interaction confirmed the order and orientation accuracy for four accessions (Supplementary Fig. 1). The average contig N50 of the newly assembled genomes was 19.16 Mb (from 10.95 Mb to 30.43 Mb), and the LAI score was on average 21.71 (from 20.2 to 23.9), equivalent to those of previous assemblies using PacBio CLR mode or Nanopore sequencing (Supplementary Fig. 2a). Averagely, 97.32% of the 4896 core conserved Poales genes (BUSCO) were assembled (Supplementary Table 1). The whole-genome synteny against the reference assembly Nipponbare and gapless assembly MH63RS3 (Song et al., 2021) suggested high completeness (Supplementary Fig. 3).

Before adopting more assemblies into the construction of the rice pan-genome, we systematically evaluated the base-level accuracy and assembly completeness on rice genomes, including recently released assemblies (Qin et al., 2021; Zhou et al., 2020; Zhang et al., 2022). The *k*-mer-based assembly validation results revealed higher assembly consensus quality values (QVs) for HiFi assemblies generated in this study (average QV = 44.16) than those for previous assemblies (Fig. 1a; Supplementary Fig. 4a). We quantified the assembly accuracy by calling homozygous single nucleotide polymorphisms (SNPs) and short insertions and deletions (InDels) by mapping available NGS reads for each accession against its own assembly. At the single-base level, averagely the HiFi assemblies showed fewer errors than the PacBio CLR mode and Nanopore sequencing (Fig. 1a; Supplementary Fig. 4b). In terms of InDels, HiFi assemblies showed no obvious differences from CLR mode assemblies, but there were fewer InDels in HiFi than in Nanopore assemblies. The annotation of the potential assembly errors for each accession suggested high (stop loss and gain, start loss, and frame-shift variants) and moderate effects (inframe insertion/deletion and missense variants) in the predicted gene models (Supplementary Fig. 4b). The low assembly quality at the base level directly interferes with the accuracy of haplotype inference, especially for Nanopore-based assemblies despite further polishing using short reads; thus, the recently released Nanopore-based assemblies of cultivated rice were excluded. Given that the available assemblies for wild rice using PacBio sequencing were only for W2014 (Ma et al., 2020) and IRGC106162 (Xie et al., 2020), nine Nanopore-based wild assemblies (Shang et al., 2022) were adopted. Finally, 11 wild, 51 cultivated and 12 weedy rice assemblies were used in the following pan-genomic analysis (Supplementary Table 1). Phylogeny based on 11.6 million whole-genome SNPs of the 74 genomes revealed four *aromatic* (aro), four tropical (trp), two subtropical (subtrp) and 13 temperate (tmp) accessions in the GJ subspecies (*n* = 23 in total) and four *aus*, three XI2, seven XI3, ten XI1A and 16 XI1B accessions in the XI subspecies (*n* = 40) (Fig. 1b). Whole-genomic features, e.g., genome size, number of annotated genes, and transposon element size and proportion, were significantly differentiated between XI and GJ (Supplementary Fig. 2b). The wild population includes two accessions from Or-3 (generally considered to be the ancestral group of GJ), four from Or-2, three from Or-1 (ancestral group of XI), and two from Or-4 (Fig. 1b). The representativeness of the 74 genomes regarding diversity was validated by a kinship analysis using a global panel of approximately seven thousand rice accessions (Supplementary Fig. 5), suggesting that the rice genomes used have covered all major taxa and all ecotypes. This provides a good opportunity to revisit the evolutionary trajectory of rice from a pan-genome view.

**Figure 1.**
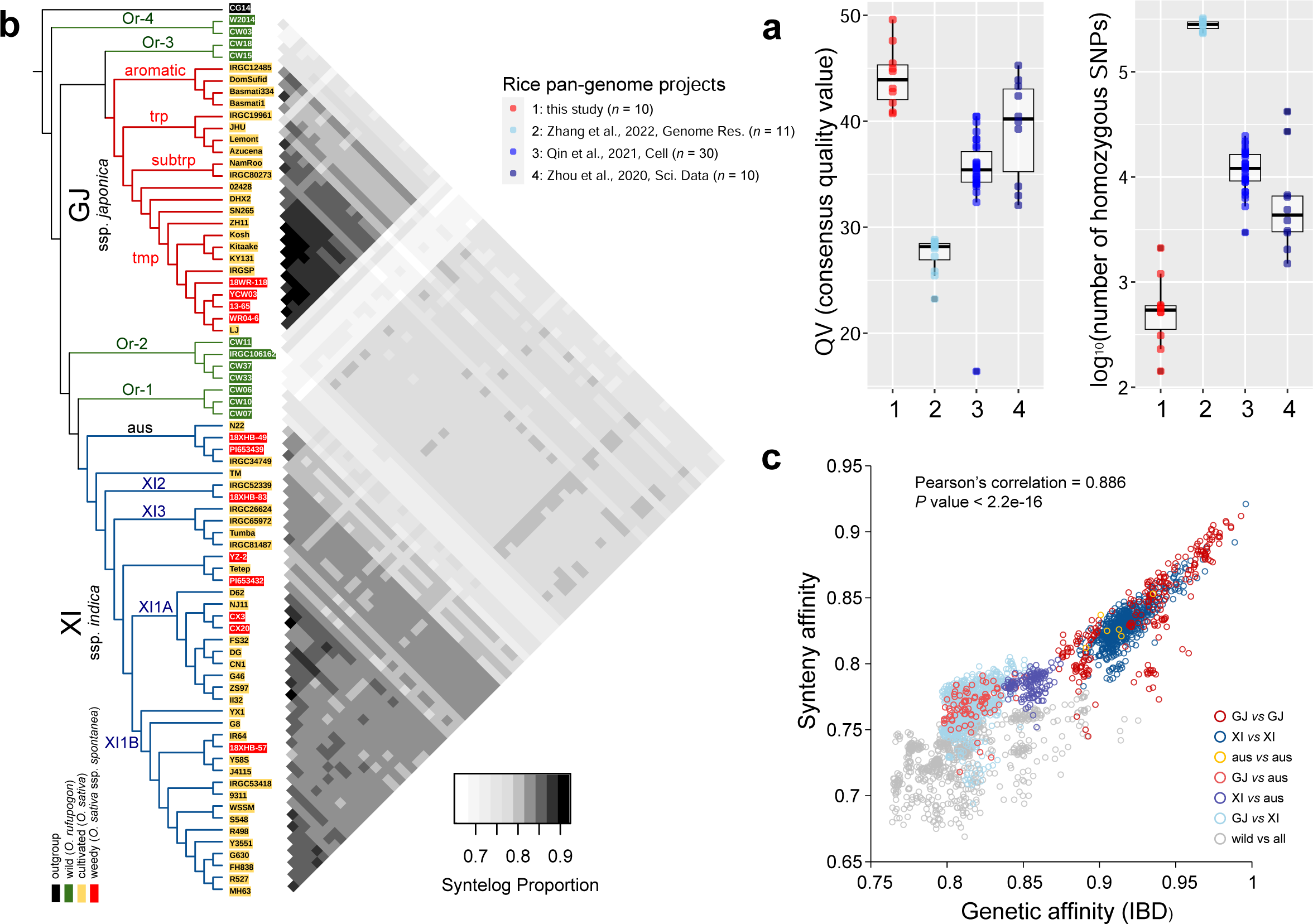
Quality assessment of rice genome assemblies and their relationship based on phylogeny and synteny. (**a**) Consensus quality values and number of homozygous SNPs based on self-mapping for genomes in four pan-genome projects. (**b**) The relationship among rice genomes measured by genetic distance (IBD, identity-by-descent) based on SNPs and synteny based on gene orders.

### Syntelog-based pan-genome of rice

Compared to variation maps obtained by mapping whole-genome resequencing short reads against one reference genome, *de novo* assemblies provide accurate and complete haplotype-resolved genetic and predicted protein sequences as well as genomic coordinates along chromosomes. To incorporate positional information, we constructed a synteny-based pan-genome by clustering approximately 3.10 million genes from the 74 rice genomes with SynPan (see Methods). Pairwise alignments for each pair of genomes were performed first to identify inter-individual syntelogs (syntenic orthologs) and merged together. Compared to the fast aligner Diamond (Buchfink et al., 2021), which is designed for high-performance analysis of big sequence data, pairwise alignment using BLASTP identified more syntelogs between two genomes, with an average addition of 669.1 pairs per genome alignment (Supplementary Fig. 6). Therefore, the syntelog datasets based on the BLASTP approach were used in the pan-genome construction.

First, a coalescence-free kinship was built based on pairwise whole-genome synteny (Fig. 1b). Generally, the affinity measured by whole-genome synteny among accessions is linearly correlated with that using identity-by-descent (Pearson’s correlation = 0.886, *P* value < 2.2e-16) (Fig. 1c). For GJ groups (aro, trp, subtrp and tmp), each group showed a closer relationship to the other GJ groups, and Or-3 was the closest wild-type group. For XI groups, *aus* exhibited apparent differences from the other XI groups (XI2, XI3, XI1A and XI1B) in genomic arrangements (Fig. 1b). Although *aus* is sister to other XI groups and nested within wild group Or-1 in the phylogenetic tree (Fig. 1b), the kinship between *aus* and Or-1 measured by synteny was distant and even larger than that between *aus* and Or-4 (the basal wild outgroup), which implied that the wild ancestor of *aus* is distinct from that of the other XI groups (Supplementary Fig. 7a). Notably, some XI accessions suggested closer affinity with GJ, which reflected the inter-subspecies hybridization in modern breeding (e.g., Y58S and its offspring J4115) (Fig. 1b; Supplementary Fig. 7b). Y58S is an XI-type photothermosensitive genic-male-sterile (PTGMS) line with the characteristics of high-light-efficiency use and disease and stress resistance, and it is widely used for the breeding of two-line hybrid rice varieties, especially super hybrids. The GJ accession Lemont is one of the parental lines used in the breeding of Y58S (China Rice Data Center, https://ricedata.cn/).

Based on whole-genome synteny, 175,528 syntelog groups (SGs) were clustered, with 13,908 core (present in the genomes of all accessions), 14,423 soft-core (present in the genomes of >90% accessions), 62,425 dispensable (present in the genomes of less than 90% of all but at least two accessions) and 84,772 private SGs (only present in a single genome) (Fig. 2a; Supplementary Fig. 8a). The SG size is 1.62 times larger than the number of orthogroups or ortholog groups (OGs) clustered by the Markov clustering (MCL) algorithm (*n* = 67,080 when the inflation parameter is 1.5, including 15,749 core, 11,725 soft-core, 18,901 dispensable and 20,705 private OGs). Even though the inflation parameter was set as 2.5, the OG size increased to *n* = 72,226, which included only 41.1% of the SG size (Supplementary Fig. 9). In a perfect ortholog group, one gene from one genome is expected, despite individual-specific duplication. The MCL method does not perform well in distinguishing paralogs against orthologs, especially in the core and soft-core groups (Supplementary Fig. 8b). By taking advantage of the genomic coordinates of genes in assemblies, the SGs provide a more precise and accurate ortholog classification.

**Figure 2.**
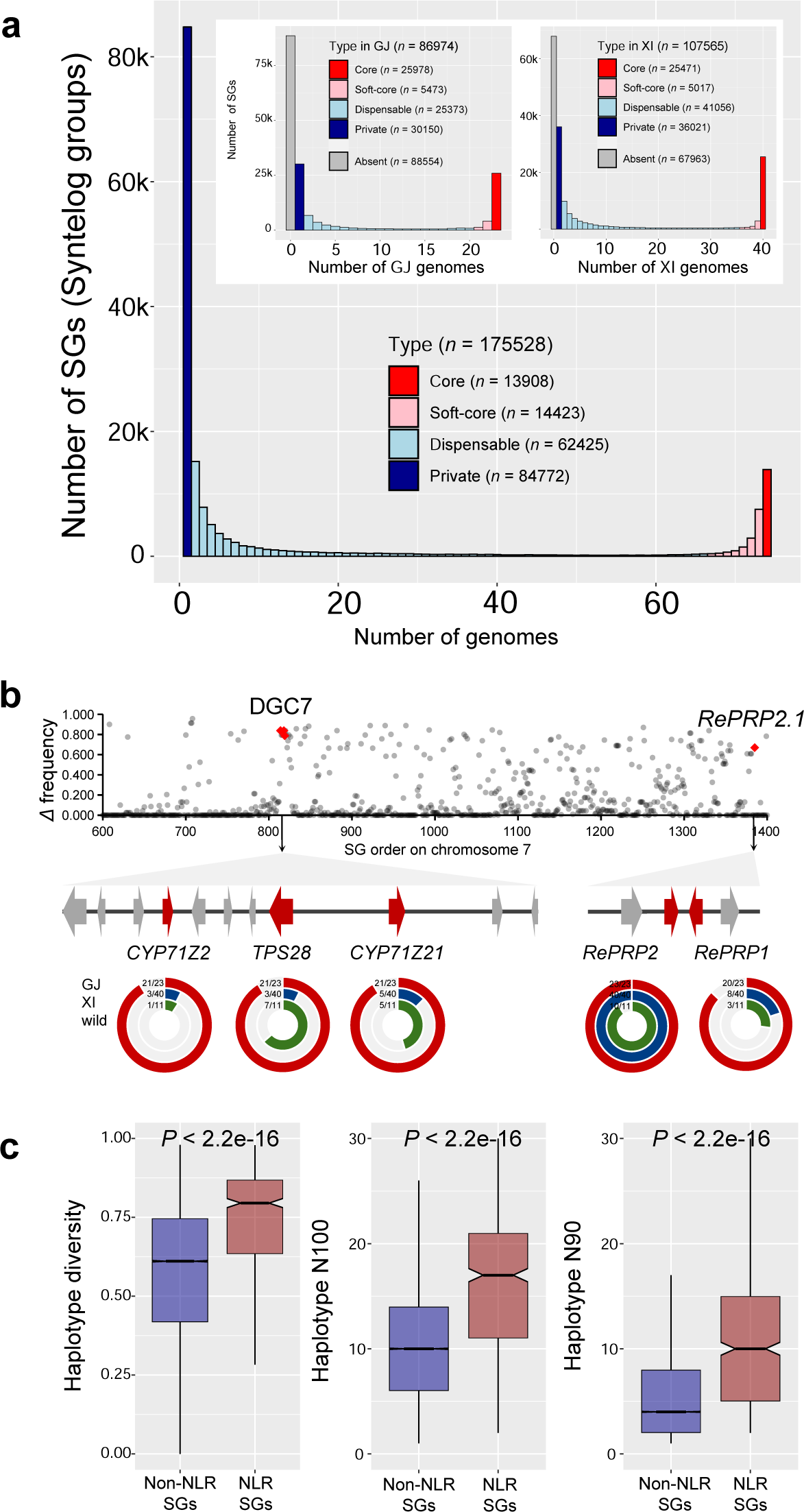
Syntelog-based pan-genome of rice and differential gene PAVs in subspecies. (**a**) Number of syntelog groups (SGs) represented in all 74 rice genomes versus the number of genomes. The subspecies pan-genome compositions for GJ and XI were extracted from the whole rice pan-genome. The numbers of core, soft-core, dispensable, private and absent SGs were counted. (**b**) Frequency differences in gene presence between GJ and XI along chromosome 7. Each dot represents one SG. The gene PAV profiling of the antimicrobial diterpenoid biosynthetic gene cluster DGC7 and tandem duplicates *RePRP2.1* and *RePRP2.2*, suppressors of root cell expansion, is shown.

Approximately 34.5%, 34.9%, 27.7% and 2.8% of genes were assigned to core, soft-core, dispensable and private SGs, respectively (Supplementary Fig. 10a). Protein domains could be identified using InterProScan (Jones et al., 2014) in a total of 81.5% and 68.9% of core and soft-core genes, which is nearly twice as high as 36.9% and 28.7% for dispensable and private genes, respectively (Supplementary Fig. 10b). Protein domain gain-and-loss variation was found within a single SG, indicating functional diversification in rice evolution. In total, 1.10% of core genes from 14.1% of core SGs suggested domain gain-and-loss, and these percentages were lower than those for soft-core and dispensable types (2.0% of soft-core genes in 19.5% of soft-core SGs and 5.0% of dispensable genes in 18.9% of dispensable SGs) (Supplementary Fig. 10c). For example, two adjacent genes, *SaM* and *SaF,* encode a ubiquitin-like modifier E3 ligase-like protein and an F-box protein, respectively, and their interactions are responsible for XI-GJ hybrid male sterility (Long et al., 2008). The SGs of *SaM* and *SaF* were both soft-core genes present in 68 and 72 genomes, respectively. The domains of all *SaF* syntelogs are completely annotated, while the domains of 29 syntelogs from *SaM* SG were not found, which suggested the dynamics domain gain-and-loss in conserved SGs.

Only 49.5% (*n* = 86,974) and 61.3% (*n* = 107,565) of SGs were present in the GJ and XI subspecies, respectively, indicating the large genomic diversity of wild accessions and genetic bottlenecks due to artificial selection or genetic drift (Fig. 2a). In total, 4,662 SGs showed presence-and-absence (PAV) frequency biases between GJ and XI, with a frequency difference greater than 0.6, including 1,918 SGs absent from the reference genome Nipponbare. For example, the key genes in a casbene-derived diterpenoid biosynthetic gene cluster DGC7 on chromosome 7 suggested differential PAVs between GJ and XI, where *CYP71Z21*, *TPS28* and *CYP71Z2* were almost absent in XI but fixed in GJ (Fig. 2b), which may be related to the differential responses of subspecies to biotic stress (Zhan et al., 2020). *RePRP1* and *RePRP2* are functionally redundant suppressors of root cell expansion (Tseng et al., 2013). Both *RePRP1* and *RePRP2* were present in GJ accessions, while only *RePRP2* was found in most XI and wild accessions, which implied that the copies of RePRP genes may underlie the differential root development in different subspecies (Fig. 2b).

### Rice pan-NLRome

Pan-genomes provide an opportunity to uncover the diversity of highly variable gene families, such as those encoding nucleotide-binding leucine-rich repeat (NLR) proteins related to disease resistance, across species (so-called pan-NLRome). A total of 37,079 NLR genes in 74 rice genomes were identified, ranging from 452 (W2014 from wild group Or-4) to 532 (FH838 from group XI1B) NLRs per genome. Distinct from the NLR composition in the *Arabidopsis thaliana* pan-NLRome, no TIR-NLR (TNL) genes were found in the rice genomes. Rice NLRs were categorized into three types: CNL (including CC-NB-LRR or CC-NB), NL (including NB-LRR or NBS) and null (NLR genes identified by syntelogs whose encoding proteins contain no canonical NBS domain). The sizes of NLRs in cultivated and weedy genomes were both significantly larger than those in the wild group (*P* = 2.8e-5 and 4.3e-5, Student’s *t* test) (Supplementary Fig. 11a), which could be the consequence of disease-resistance gene aggregation during domestication and improvement. However, the relatively low assembly quality of the wild genomes may also be related to this difference, considering that the completeness of assemblies was significantly correlated with the NLR size (Pearson’s correlation = 0.53, *P* value = 9.7e-7) (Supplementary Fig. 11d). At the subspecies level, although NLR gene numbers were similar between XI and GJ, the XI genomes contained more NLs than those in GJ (*P* = 4.3e-10, Student’s *t* test), and GJ had more CNLs (*P* = 3.6e-6, Student’s *t* test) (Supplementary Fig. 11b). Null NLRs without canonical NBS domains were more abundant in GJ than in XI (*P* = 1.6e-10, Student’s *t* test), implying that more NLRs in GJ may degenerate functionally by losing domains (Supplementary Fig. 11b). Adopting the definition by Wang et al. (2019) of an NLR cluster containing more than two NLR genes distributed within a 300-kb genomic region, 43.2%-59.9% of NLR genes in rice were located in such clusters, where GJ showed more clusters in both numbers and proportions across all NLRs than XI (Supplementary Fig. 11c). Head-to-head pairing of NLR genes is highly associated with disease resistance in plants, with one NLR acting in effector recognition (known as a sensor) and the other acting in signaling activation (known as a helper). We found 28 to 54 such paired NLRs per genome. A total of 6,008 NLRs encoded at least one non-canonical NLR domain (NBS, LRR and CC), also known as the integrated domain (ID), representing 116 distinct Pfam domains and 16.2% of the total NLRs, which was much higher than that in *Arabidopsis thaliana* (5.0%). We identified 480 distinct architectures in the pan-NLRome, of which only 67 were found in the Nipponbare reference genome (IRGSP v1.0). Fewer than 3% of architectures, 12, correspond only to different configurations of the canonical CC, NBS and LRR domains, even though they accounted for the majority (83.8%) of NLRs.

As expected, the haplotype diversity of NLR SGs was significantly higher than that of non-NLR SGs (Fig. 2c). In total, 0.64% (*n* = 238) of all NLRs were present in only one accession, representing 238 private SGs, while the remaining 10,878 (29.3%), 14,760 (39.8%) and 11,203 (30.2%) NLRs grouped into 147 core, 208 soft-core and 407 dispensable SGs, respectively. Although the NLR syntelogs among individuals were well defined by genome synteny, within a single SG, the functional types (CNL, NL or null) and structural types (e.g., in clusters or pairs) were diversified, particularly for the core NLR SGs (Supplementary Fig. 12a). Nucleotide diversity for NLRs in core and soft-core SGs was lower than that in dispensable SGs but was not significant. Tajima’s *D* values, which indicate balancing and purifying selection, showed no significant differences across different NLR classes, with all classes containing extremes in both directions (Supplementary Fig. 12b).

### Rice origin based on the mosaic genomic map of syntelog haplotypes

Although whole-genome single nucleotide variants provide a comprehensive variation landscape in genome evolution, millions of markers could be somewhat redundant because of synonymous mutations. Here, we used the predicted amino acid sequences of genes to dissect the genomic ancestry of each gene in all rice genomes, given that the protein sequences function directly and are degenerated with high tolerance to synonymous mutations. Haplotypes were first assigned for each of the 49,438 SGs whose syntelog members were present in at least ten genomes. The haplotype complexity for each SG based on protein sequences was highly reduced compared with those using full-length gene sequences and coding sequences, as indicated by haplotype diversity (the average haplotype differences between any two members from a single SG), haplotype N100 (total number of unique haplotype sequences) and N90 (the least haplotype number that needs to be included for covering 90% of sequences in an SG) (Supplementary Fig. 13a; Supplementary Fig. 14a). The haplotype number and diversity were higher for core and soft-core SGs than dispensable SGs. On average, 11.2 haplotypes were found for each SG, where haplotype numbers for core (*n* = 12.0) and soft-core SGs (*n* = 12.8) were higher than that for dispensable SGs (*n* = 9.5). Specifically, 147 core SGs were extremely conserved with the fully identical protein sequences of housekeeping genes involved in fundamental biological processes, such as *LEA5* in late embryogenesis (He et al., 2012), *eIF-4A* and *Os-eIF6;1* in translation initiation (Kato et al., 2010), *OsFd1* in photosynthesis (He et al., 2020) and *OsAtg8* in autophagy (Izumi et al., 2015). As expected, the haplotype diversity of XI was significantly higher than that of GJ, with average haplotype diversity values of 0.578, 0.362 and 0.441 for all rice accessions, GJ and XI, respectively (Supplementary Fig. 13b).

**Figure 3.**
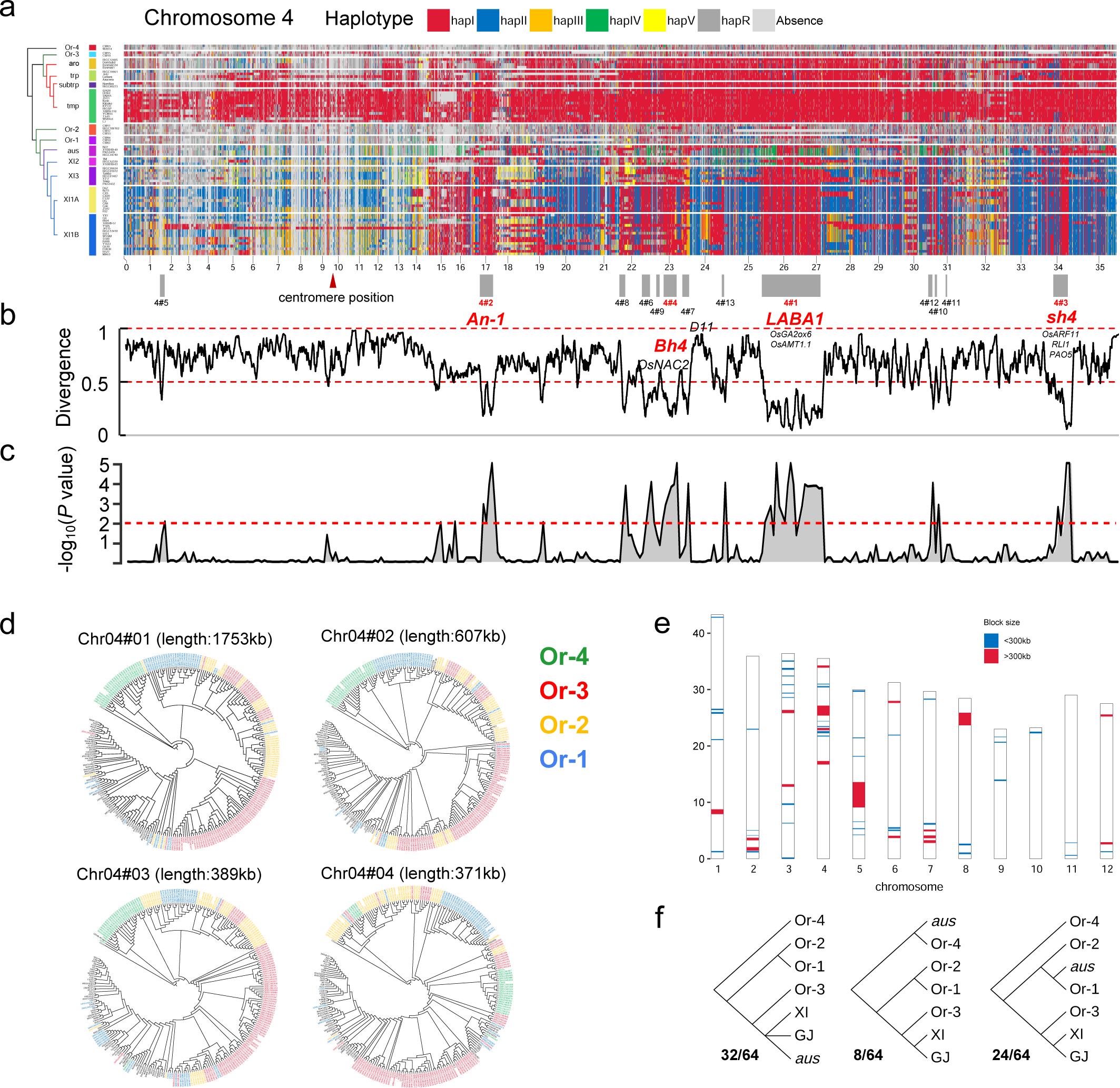
Syntelog-based ancestral haplotypes suggest widespread genomic introgression in rice evolution. (**a**) Ancestral haplotype landscape of SGs on chromosome 4. An SNP-based phylogeny of rice groups is illustrated on the left. For each SG, seven blocks with different colors represent different haplotypes of predicted protein sequences. Candidate introgression regions are numbered by genomic length and marked by gray blocks. The centromere position information of chromosome 4 is obtained from the Rice Genome Annotation Project and indicated by a red triangle. Functionally important genes are annotated within each block, and domestication genes are highlighted in red. (**b**) Haplotype divergence between XI and GJ for all SGs on chromosome 4. (**c**) Significance test on the non-random distribution of clustered SGs in introgression blocks by sampling 100,000 replicates. The horizontal red dashed line represents a *P* value of 0.01. (**d**) Phylogeny analysis of four large introgression blocks (length greater than 300 kb) indicates a single origin of domestication alleles from the Or-3 wild group. Four wild groups are highlighted in different colors. (e) Final putative introgression blocks from proto-GJ to XI, combing the evidence from haplotype divergence, phylogeny and ABBA-BABA tests. (**f**) Complex genomic contributions from wild groups to the emergence of *aus* group revealed by the phylogenetic trees in 64 introgression blocks.

**Figure 4.**
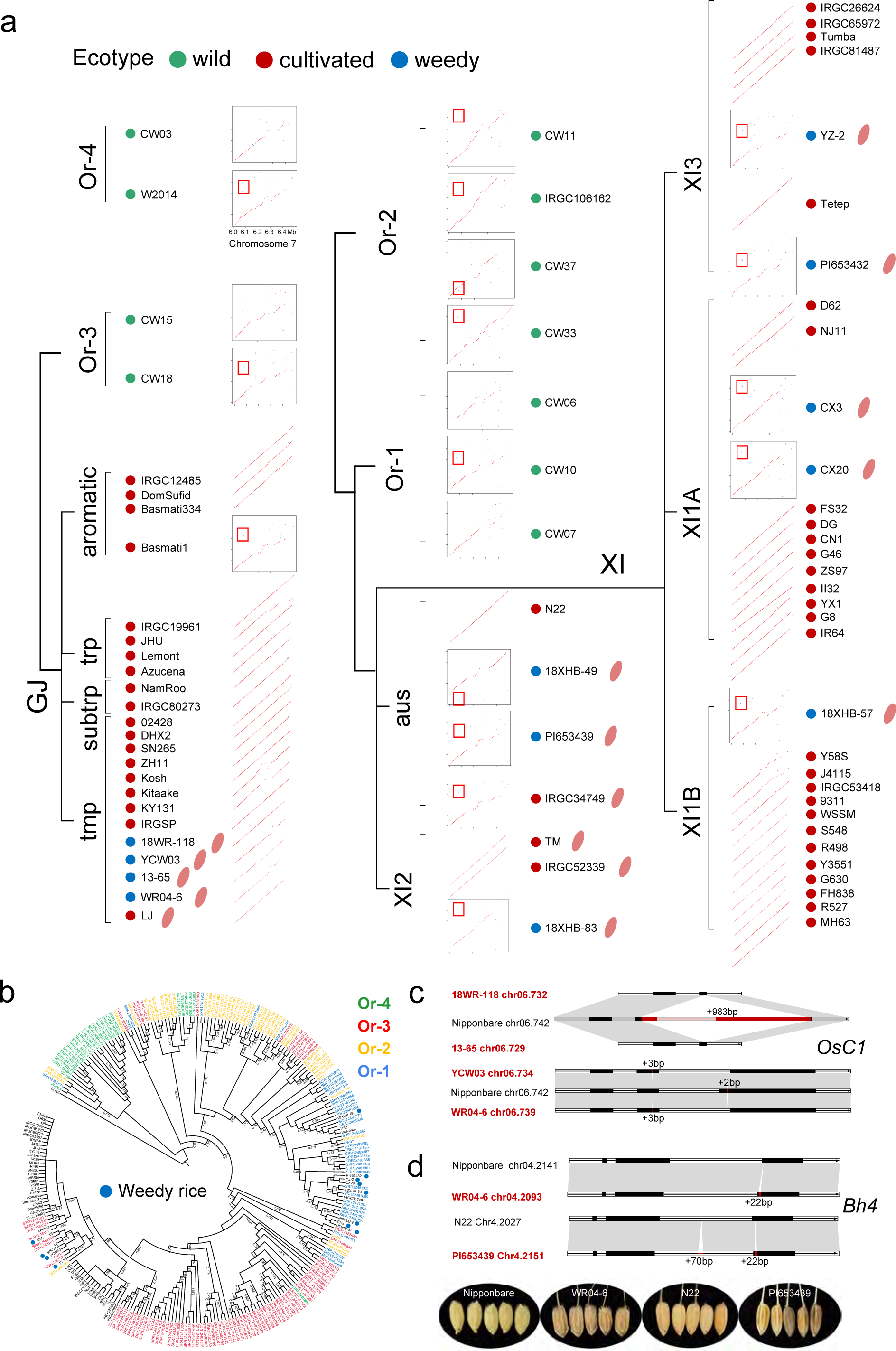
Structural variations in rice de-domestication. (**a**) Dot plots comparing all 74 assemblies against the de-domestication genomic island (from 6.0 to 6.5 Mb) on chromosome 7 of Nipponbare. The predominant translocations in wild and weedy accessions are highlighted by red boxes. The pink seed icons behind the accession IDs represent red or brown pericarp. (**b**) Phylogeny revealed by SNPs in the *Rc* region indicates introgression from Or-1 and Or-3 to XI and GJ, respectively. Labels with green, red, yellow and blue represent accessions from Or-4, Or-3, Or-2 and Or-1, respectively. Blue dots represent weedy accessions. The numbers on each branch indicate bootstrap values of less than 90%, based on 1000 replicates. (**c**) Structural variations in *OsC1* between the weedy and cultivated GJ accessions. Black rectangles represent exons. (**d**) Structural variations in *Bh4* between weedy and cultivated accessions and their seed appearances.

Using a semi-supervised approach, haplotypes were reassigned by comparing their abundance in different groups for each SG, labeled as hapI to hapV, hapR (all other rare haplotypes) and absence (represented by red, blue, orange, yellow, green, dark gray and light gray blocks in Fig. 3a and Supplementary Fig. 15) to represent the ancestral sources of rice haplotypes (Supplementary Fig. 13b). Apparently, the whole-genome haplotype maps visually reflected the shared haploblocks at the individual and group levels. For example, mosaic genomes of SN265, DHX2 and 02428 from the GJ tmp group and Y58S, J4115 and FH838 from the XI1B group suggested large introgressed regions from the other subspecies (Fig. 3a; Supplementary Fig. 15). Haplotypes in two Mb-level genomic regions at the head of chromosomes 6 and 7 showed that only XI1B in XI groups was shared with GJ, which was consistent with the fact that group XI1B was mainly composed of modern cultivars that were frequently bred by utilizing genetic resources from other subspecies (Supplementary Fig. 15).

**Figure 5.**
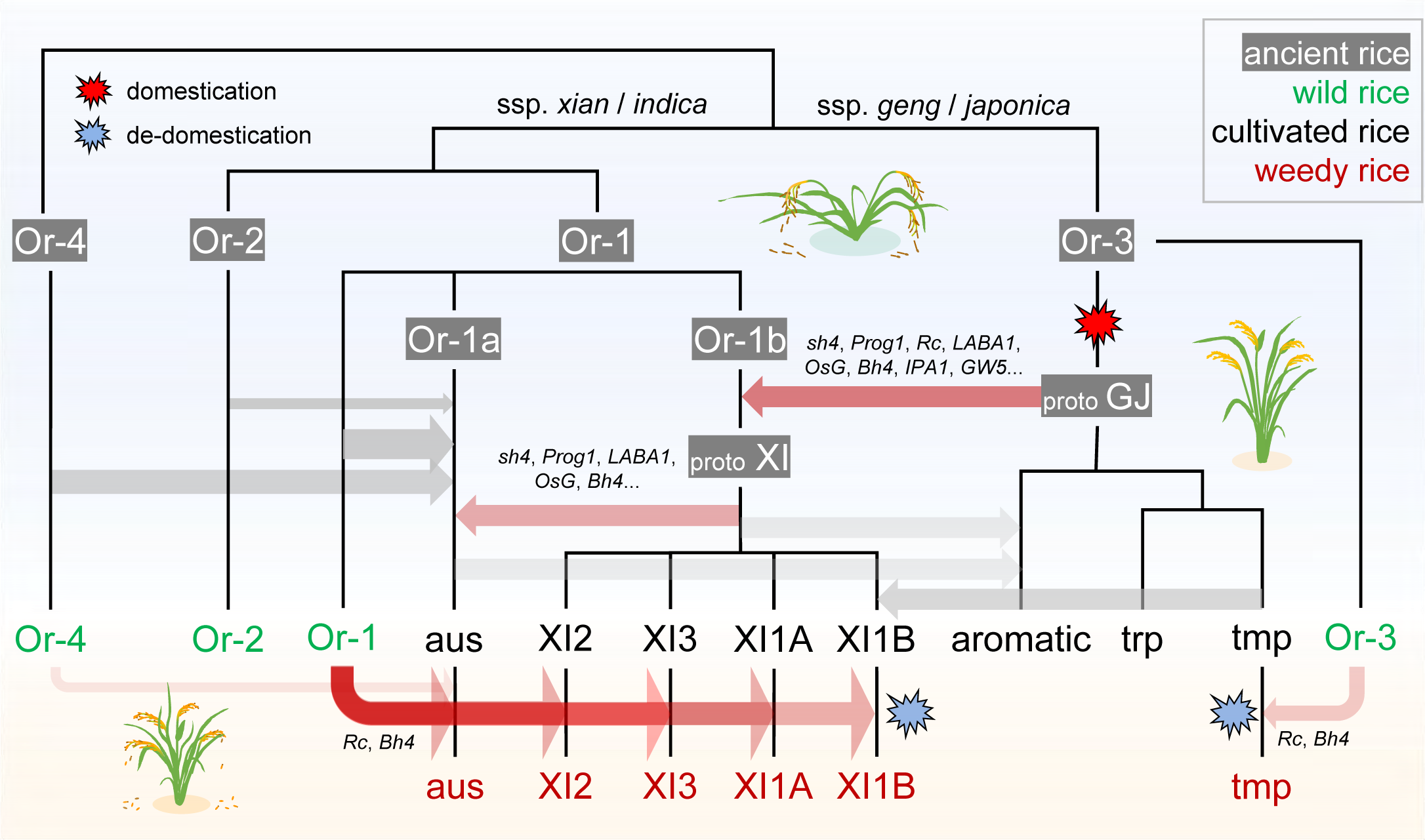
A brief schematic illustration of Asian rice evolution. The evolutionary scenario highlights that complex introgression events have contributed indispensably to rice domestication and de-domestication.

As observed from the mosaic genomic map, *aus* showed obvious differences from the other XI groups; thus, the *aus* group was excluded from XI in the following analyses (Fig. 3; Supplementary Fig. 15). We used inter-subspecies diversity to quantify haplotype divergence (HDG) between GJ and XI (Supplementary Fig. 13a). The average haplotype divergence between GJ and XI was 0.667, and 46.0% of SGs (*n* = 22,745) suggested high haplotype divergence with HDG > 0.8, indicating great divergence between subspecies at the translation level. Most (14/21) well-known improvement genes (Chen et al., 2019) showed high divergence, such as *DEP1* (HDG = 1.000), *Sd1* (0.940), *TAC1* (0.930), *GS5* (0.913), *GW6a* (0.925), *TGW6* (0.962), *GW7* (0.954), *GW8* (0.976), *Ghd7* (0.963), *Ghd8* (0.984), *Hd1* (0.979), *NRT1.1B* (0.977), *DRO1* (0.850), and *Chalk5* (0.933), implying independent selection for yield- and flowering-related genes in the improvement of XI and GJ. Notably, 720 SGs showed weak divergence with HDG < 0.2, including essential domestication genes *Prog1* (HDG = 0.025), *GAD1* (0.110) and *sh4* (0.155). The SGs with low divergence (HDG < 0.5) were clustered into local genomic blocks, with a total length of 23.38 Mb (6.24% of Nipponbare assembly) in 73 blocks, including 2,786 SGs (6.90% of all SGs in Nipponbare), implying putative genomic introgression between XI and GJ (Fig. 3a and 3b; Supplementary Fig. 15; Supplementary Tables 2 and 3). Typically, a total of 18 blocks were beyond 300 kb, and the largest three blocks were on chromosomes 5, 8 and 4, spanning over 4.48, 2.22 and 1.75 Mb, respectively. Introgression and incomplete lineage sorting (ILS) would both result in haplotype similarity and low divergence between sequences from two lineages. To distinguish introgression from ILS, which is more randomly distributed along chromosomes (Wu et al., 2022b; Edelman et al., 2019), we tested the significance of lowly divergent SG clustering by 100000-times random sampling. The significant nonrandom distribution of the lowly divergent SGs in these blocks implied that inter-subspecies introgression caused low GJ-XI divergence, rather than ILS (Fig. 3c; Supplementary Fig. 15). Additionally, as expected, the relative divergence indicated by the synonymous substitution rate (*K*s) between XI and GJ genes in putative introgression blocks was significantly lower than that of genes in adjacent genomic regions (Supplementary Fig. 16).

Most domestication genes (9/10) underlying key domestication syndromes (Chen et al., 2019) were found in the introgression blocks (Supplementary Fig. 15; Supplementary Table 2). On chromosome 4, four introgressed blocks larger than 300 kb were found, including domestication genes *LABA1* and *An-1* responsible for awn presence-and-absence and length in blocks 4#1 and 4#2 (Luo et al., 2013; Hua et al., 2015), *sh4* for grain shattering in block 4#3 (Li et al., 2006), and *Bh4* for hull color in block 4#4 (Zhu et al., 2011)(Fig. 3a). On chromosome 5, *GW5* from block 5#3 is a QTL for grain width and weight (Liu et al., 2017). On chromosome 7, *Prog1* was located in block 7#1, underlying the transition from prostrate plant architecture to erectness during domestication (Jin et al., 2008). *GAD1*, encoding a secreted awn development-related peptide, and *IPA1*, which is considered a typical improvement gene controlling ideal plant architecture and immunity, were both located in block 8#1 on chromosome 8 (Jiao et al., 2010; Wang et al., 2018a). Interestingly, except for *IPA1*, no other improvement and diversification genes were found in the introgression blocks, which suggested that introgression events occurred in the initial period of domestication (Supplementary Table 2).The largest introgression block, 5#1, with a length of 4.48 Mb, spanned the centromere region on chromosome 5 (Supplementary Fig. 15). Although more than three hundred genes were annotated in this region, no known domestication-related genes were found. The presence of block 5#1 with low divergence between XI and GJ was probably due to suppressed recombination over the centromere region rather than linkage by key domestication genes, which provides an ideal clue to subspecies introgression.

We further utilized 184 wild genomes with high whole-genome sequencing depth (averagely >8×), encompassing four groups Or-1 to Or-4, to confirm the introgression and trace the spread routes of domestication haplotypes (Supplementary Fig. 17). Totally the introgression inference in 64 blocks (96.2% in genomic length) were supported by phylogeny (Supplementary Table 3). Phylogenetic trees of these blocks (including all 18 blocks longer than 300 kb) indicated that GJ and XI accessions were nested within the Or-3 group of wild rice, indicating that Or-3 was the shared wild progenitor group of GJ and XI in most introgression blocks, and that the domesticated alleles in XI were likely to be derived from proto-GJ by introgression with local wild rice from Or-1 (Fig. 3d; Supplementary Table 3). Notably, between the major clade of domestication haplotypes and the Or-3 clade, wild accessions from the Or-1 or Or-2 group were commonly observed (Fig. 3d), which implied that these haplotypes contained relict ancient domesticated alleles, although some introgression events from cultivated to wild rice were observed. We speculate that a gene pool under early domestication was introduced into South Asia and partially maintained in the genomes of present Or-1 or Or-2 wild rice.

Statistical ABBA-BABA tests were also performed to confirm the introgression inference. In a total of 57 blocks (21.2Mb in length), high *f*d values were observed in the models with introgression direction from tmp(GJ) to XI groups (Supplementary Figs. 18 and 19). Thus combing haplotype inference, phylogeny, and ABBA-BABA test, 65 blocks (22.6 Mb in length) were finally determined as introgression regions from GJ to XI (Supplementary Table 3). Auxin related pathways were significantly enriched (Supplementary Table 4). Besides well-known domestication genes related awn presence, shattering, and tiller angle, many seed dormancy or germination related genes were observed in introgression blocks (Supplementary Table 5). Block 3#5 from chromosome 3 included a seed dormancy-related gene *OsG*, which has been parallelly selected in multiple crop families (Wang et al., 2018a), and *qLTG3-1*, a major quantitative trait locus controlling low-temperature germinability (Fujino et al., 2008). *OsC1*, which regulates hull pigmentation and pre-harvest sprouting, was located in block 6#3 on chromosome 6. Yield related genes include at least *HOX3* (1#2), *SPL6* (3#6), *GPA3* (3#6), *OsNAC2* (4#4), *D11* (4#7)and *FZP* (7#5).

We noticed that the *aus* group exhibited differences from the GJ and XI groups in most blocks (e.g., 1#1, 2#2 and 5#1). Phylogenetic analysis revealed that XI and *aus* belonged to two separate subclades, although both were nested within Or-1 group (Supplementary Fig. 17a). There were 32 blocks where the closest wild group to *aus* was Or-3, similar to XI and GJ, while in the other 24 and 8 blocks, the Or-1 and Or-4 groups were the wild progenitors of *aus*, respectively (Fig. 3e). In ABBA-BABA analysis, only 23 blocks showed gene flow signals from GJ to *aus* (Supplementary Fig. 18; Supplementary Table 3). Therefore, the *aus* group shared some domesticated alleles, suggesting that the introgression from proto-GJ or Or-3 to XI probably occurred after the divergence between XI and *aus*, and some domesticated alleles or blocks were subsequently transferred from XI to *aus* (e.g., 4#1) (Supplementary Fig. 15). Combining the evidence from whole-genome pairwise synteny (Fig. 1b), the *aus* group differs from other XI groups and has a novel evolutionary process.

### Structural variations during rice de-domestication

Previous studies have found a 0.5-Mb de-domestication genomic island on chromosome 7 from 6.0 Mb to 6.5 Mb that plays essential roles in rice feralization (Qiu et al., 2020). Within this region, the key gene *Rc* regulating the red pericarp and a cluster of seed storage-related genes (six *RAL* and three *LtpL* genes) contribute to the fitness of weedy rice. The genomic synteny in this region was investigated between weedy and cultivated rice. Although weedy rice originated independently and repeatedly from cultivated rice, the genomic landscape of XI weedy accessions in this region showed distinct patterns against their corresponding closest cultivated genomes (Fig. 4a). This weedy pattern was prevalent in the genomes of wild accessions.

Notably, a remarkable 10-kb translocation (including one gene encoding an RNA polymerase II transcription subunit) was found in all XI weedy rice when aligned to the Nipponbare assembly but was absent from all XI cultivated rice (except cultivar IRGC34749 with a red pericarp) and all GJ accessions (except Basmati1) (Fig. 4a). This translocation was found in 7 of 11 wild accessions. Thus, it was speculated that genomic introgression of the de-domestication genomic island from wild rice contributed to the feralization of XI cultivated rice and the emergence of XI weedy rice. A phylogenetic tree based on SNPs around the *Rc* region further supported the introgression and suggested that the Or-1 group from Southeast Asia was the ancestral progenitor of the de-domestication genomic island in XI weedy rice (Fig. 4b). For GJ weedy rice, no obvious signals of wild introgression in this de-domestication island were found (Fig. 4a). However, from the phylogeny of *Rc*, weedy accessions were clustered together and nested within some Or-3 wild accessions, which implied that Or-3 may be the donor of the *Rc* haplotype underlying the red pericarp (Fig. 4b). However, it should be noted that some cultivated rice also showed a red pericarp (e.g., LJ from GJ), suggesting another potential origin from local landraces of red rice.

To gain insight into detailed structural variations during de-domestication, we compared the genomic sequences between weedy accessions and their closest corresponding cultivated accessions based on phylogeny (Fig. 1b). Given that de-domestication occurred recently (Qiu et al., 2020; Sun et al., 2019), no large chromosome rearrangements were observed from genome synteny, except for the comparison between Tetep and PI653432, owing to the low assembly quality of Tetep (Supplementary Fig. 20). On average, 73,111 small insertions and deletions (InDels, ≤50 bp) and 2,810 large structural variants (SVs, >50 bp) were identified in ten weed-cultivar pairs, spanning genomic regions from 2.2 to 13.8 Mb in total (Supplementary Fig. 21). The number of structural variants was lower for the GJ weedy-cultivated pairs than for the XI and *aus* pairs. This could be a result of the recent origin of GJ weedy rice but more ancient origin of XI and *aus* (Qiu et al., 2020) and more introgression from other taxa or (sub)-species into XI and *aus* cultivated rice. The XI cultivar NJ11 and its weedy descendant CX20 were reported as a typical case of recent de-domestication events (Qiu et al., 2020). When the HiFi-based genome assemblies were compared, a minimum number of SVs (*n* = 1,299) were detected, where six SVs larger than 100 kb were close to peri-centromeric regions, and the largest SV spanned 823 kb and harbored 66 genes on chromosome 7 (Supplementary Fig. 22). *OsGWD1*, which is involved in transitory starch degradation in source tissues and is also a positive regulator of rice seed germination, was lost in the weedy CX20 genome, which may be related to the difference in rice quality between cultivated and weedy rice (Wang et al., 2021). Additionally, we noticed the absence of the rice blast resistance gene *Pid2* in weedy CX20. Compared with NJ11, equivalent insertions (*n* = 19,340 for small insertions and *n* = 666 for large insertions) and deletions (*n* = 19,410 for small deletions and *n* = 633 for large deletions) were found in CX20, spanning a total gain-and-loss length of 2.91 and 2.97 Mb, respectively. Gene Ontology analysis suggested that SV-associated genes were enriched in the biological process of reproductive system development (adjusted *P* = 0.00065).

Despite the independent and recurrent de-domestication events observed from the phylogeny, we found 3,614 SGs in which at least one SV was detected in at least four de-domestication lineages from six groups (tmp, *aus*, XI2, XI3, XI1A and XI1B), implying potential convergent genetic mechanisms underlying feralization. Within them, a 14-bp PAV in *Rc* was identified in all XI and GJ weedy lineages (Wu et al., 2021). SVs were found in other domestication-related genes regulating seed shattering, hull color and seed dormancy or germination, of which most were located in regulatory and intron regions (Supplementary Table 6). For shattering, a 2-bp insertion in exon 1 of *sh4* and a 12-bp deletion in exon 1 and a 146-bp deletion in exon 6 of *SHAT1* were found in the *aus* and XI weedy genomes. Despite *Rc*’s role in regulating seed dormancy and germination, SVs or InDels in *OsC1* and *Sdr4* were also found in the GJ and XI weedy genomes, respectively (Fig. 4c; Supplementary Table 6). *OsC1*, a rice R2R3-MYB transcriptional regulator that interacts with *Rc* and *OsVP1*, plays an important role in regulating preharvest sprouting tolerance in red pericarp rice (Wang et al., 2020). Two different variants in *OsC1* were found in GJ weedy accessions, which were a 983-bp deletion resulting in incompleteness of exon2 and loss of exon 3 in accession 18WR-118 from Korea and 13-65 from Italy and a 3-bp insertion combined with a 2-bp deletion in accessions YCW03 and WR04-6 from China, which led to MYB domain loss (Fig. 4c; Supplementary Fig. 23). Brown hull, which is mainly regulated by *Bh4,* is also a characteristic of feralization for some weedy rice (Zhu et al., 2011). A 22-bp insertion in *Bh4* was found in both the GJ weedy genome (WR04-6) and the *aus* weedy genome (PI653439), which corresponded to their seed hull phenotypes (Fig. 4d; Supplementary Fig. 23). The 22-bp PAV was under selection during rice domestication (Zhu et al., 2011). The phylogeny of *Bh4* confirmed that the 22-bp PAV in weedy rice was likely derived from different groups of wild rice (Or-1 for *aus* PI653439 and Or-3 for GJ WR04-6). The results also revealed that the brown or black hull color in weedy and cultivated rice was due to wild introgression (Supplementary Fig. 24). Briefly, although structural differences could be found between weedy and cultivated rice, almost no single causative mutation could explain the convergent phenotypic change for all different weedy rice groups, with the only exceptional being *Rc* for red pericarp and seed dormancy; in other words, weedy rice from different groups may have experienced independent evolution after the acquisition of *Rc* from wild rice or local landraces of red rice.

## Discussion

Two major competing hypotheses (single domestication or multiple domestication) have been proposed to describe the origin of different subspecies of cultivated Asian rice according to previous archaeological and genetic evidence (Molina et al. 2011; Huang et al., 2012; Civáň et al., 2015; Choi et al., 2017; Choi and Purugganan, 2018; Wang et al., 2018b; Carpentier et al., 2019; Zhang et al., 2021). There is no dispute that rice subspecies have different wild progenitors at whole-genome level, especially for XI and GJ, which means subspecies have multiple origins for most genomic regions (Huang et al., 2012). However, the sources of domestication alleles are debated. For well-known domestication genes, phylogeny analysis has supported single domestication (e.g. *Prog1*, *sh4*, *LABA1*, *Bh4*, *OsC1*)(Huang et al., 2012; Choi and Purugganan, 2018), while some wild accessions with domesticated alleles were located within cultivated rice, which is also observed in this study (Fig. 3d). Gene flow from cultivated to wild rice could be used to explained the sporadic distribution of wild rice within cultivated sub-trees (Wang et al., 2017). However, some studies interpreted these as ancestral domesticated alleles in wild rice prior to domestication and highlighted possibility of independent acquisition of domestication alleles from wild populations (Civáň and Brown, 2017; Civáň and Brown, 2018). To this point of view, this inference seems not reasonable. If these wild alleles emerged before domestication, their phylogenetic positions would not been within cultivated accessions.

Wang et al. (2018b) analyzed haplotypes of nine domestication and improvement genes using about 3000 rice genomes, found that many XI domesticated alleles were absent in GJ and hence concluded as a support for the hypothesis of independent domestication in XI, rather than GJ-to-XI introgression. This conclusion is somewhat arbitrary. Firstly, the filtering and selection to variants will somewhat impact the haplotype inference. Second, the method using haplotype-wise genetic distance has simplified the identification criterion of introgression and is not reliable as phylogeny approaches, or ABBA-BABA test. Third, the effects by genetic drift should be taken into consideration. Cultivated rice genomes we have sampled and sequenced now are only a subset of ancestral proto-GJ or proto-XI genetic pool, and can not fully represent the diversity in domestication. GJ has suffered a dramatic genetic bottleneck around three thousand years ago, while XI shows no obvious decline in effective population size in the last few thousand years (Qiu et al., 2020; Gutaker et al., 2020). Here, in our phylogeny analysis on putative introgression blocks, some Or-1 or Or-2 accessions were located between domestication clade and Or-3 clade, as relict alleles of proto-GJ and genomic footprints of ancient introgression, which are absent in current GJ gene pool but present in wild Or-1 or Or-2 (Fig. 3d). Therefore, the genetic drift played unneglectable roles in the presence of non-GJ domesticated alleles in XI, and the hypothesis of GJ-origin of domesticated alleles in XI can not be rejected just based on haplotype presence or absence. Indeed, the non-GJ alleles in XI domestication genes have been observed in this study. *Rc* (HDG=0.803), *An-1* (0.886) and *GW5* (0.875) have high haplotype divergence between XI and GJ, but the genomic regions they are located in showed robust introgression signals, supported by haplotype similarity or sharing, phylogeny and ABBA-BABA tests (Supplementary Tables 2 and 3). Haplotype analysis on a specific gene only sometimes will mislead because of gene diversifying after domestication, and additional approaches should be performed to verify inference in the meantime.

By taking advantage of 74 high-quality rice genomes, we revisited the origin of different rice subspecies and groups from a pan-genome view (Fig. 5). Different from previous studies using selective sweep and nucleotide diversity to infer domestication genomic regions and then reconstruct their evolutionary trajectory (Huang et al., 2012; Civáň et al., 2015), we investigated the core issues of the dispute, whether introgression from GJ to XI has happened and whether the introgressed blocks harbored domesticated alleles. Compared to whole-genome resequencing, genome assembly enables us to analyze high-throughput full-length sequences of genetic elements, which eliminates systematic errors caused by read mapping and sequencing depth. By comparing the absolute differences among predicted protein sequences to assign haplotypes, we compressed the variations within a gene, which provided us with a visual landscape of haplotype similarity among taxa in each syntelog group. Similar approaches using haplotypes of single genes or blocks has been adopted in recent studies (Zhang et al., 2021; Wang et al., 2022). A haplotype map based on SNPs from coding regions was constructed for 3,010 cultivated and 15 wild rice accessions (Zhang et al., 2021). The inferred proto-ancestors of different groups correlated strongly with wild rice from the same geographic regions, which was considered to support a multi-domestication model of rice. However, in spite of extremely insufficient number in wild rice, the whole-genome haplotype similarity among populations can not indicate how domesticated alleles come from, thus the data present in Zhang et al. (2020) could not lead to the conclusion of multi-origin domestication model in rice. Here, by comparing the identity of predicted protein sequences and the frequency of each haplotype in different groups, we assigned ancestral or dominant haplotypes for each SG and determined a total of >20 Mb genomic regions as putative introgression blocks, which avoided potential mis-assignment by hard thresholds used in conventional clustering based on genetic distance or phylogenetic relationship (Supplementary Fig. 13b). Such an ancestral genomic haploblock dissection method has also been employed in tracing the origin of polyploid wheat, and mosaic genomic graphs have suggested dispersed emergence and protracted domestication in wheat (Wang et al., 2022). In the shared haplotype blocks between GJ and XI, the majority of their phylogenetic trees, combined with the relatively low divergence or genetic distances within these regions between XI and GJ, supported the introgression of domestication genes from proto-GJ to Or-1 wild group. Interestingly, the low-recombination centromere region of chromosome 5, which suggested clear genomic affinity between subspecies, remained a relict footprint of ancient introgression (Supplementary Fig. 15). Statistical ABBA-BABA tests using high-depth sequencing wild genomes and larger genome sampling also confirmed the reliability of putative introgression blocks (Supplementary Fig. 18).

Utilizing domestication alleles from other geo-isolated populations or species has facilitated the generation of local domesticates. In wheat, dispersal domestication events generated domesticated alleles or haplotypes for different genes in different locations. Genomic introgression among different populations or species by human activities has gathered domestication alleles of different genes together, leading to the emergence and popularity of hexaploid wheat (Wang et al., 2022). Here, our results strongly confirmed the previous hypothesis that genomic introgression of domestication alleles from proto-GJ to wild Or-1 led to the emergence of XI (Fig. 5). Despite limited sampling of the *aus* type, we inferred that the ancestral wild population of *aus* was different from that of XI, although they were both clustered within the Or-1 wild group. Genomic introgression from local wild rice and domesticated XI rice may have led to the birth of *aus* rice. More genomes and detailed analysis in the future will uncover the complex evolutionary process of the *aus* group.

De-domestication is an atavistic process in domesticates that has been studied in crops (e.g., rice, wheat and sunflower) and livestock (e.g., chicken and dog) (Wu et al., 2021). How de-domestication evolves in rice genomes has been investigated in recent years (Song et al., 2014; Li et al., 2017; Sun et al., 2019; Qiu et al., 2020). A genomic island on chromosome 6 that potentially contributes to rice de-domestication syndromes (mainly red pericarp and seed dormancy or germination) has been defined (Li et al., 2017; Qiu et al., 2020). Here, our analysis combining both structural comparison and phylogenetic analysis highlighted the influences of wild introgression on the emergence of weedy rice, despite independent introgression events for GJ and XI (Fig. 5). For the brown hull of some weedy accessions, the causative structural mutation was indicated to be derived from the corresponding wild groups Or-1 and Or-3 for XI and GJ, respectively. Although pairwise whole-genome comparison identified thousands of structural variants between weedy and cultivated rice, structural convergence in different weedy-cultivated lineages was seldom observed, which implied the recurrent independent emergence of weedy traits. The mechanism underlying high shattering in weedy rice was still not well resolved from the view of structural variation in known domestication genes (except *sh4* and *SHAT1* for shattering). Given that our known shattering-related genes are almost all transcription factors, variations in other regulatory elements or even epigenetic factors may lead to high shattering in weedy rice. Overall, genomic introgression plays an indispensable role throughout the entire evolutionary trajectory of rice, from initial domestication, improvement, modern breeding and feralization.

## Materials and Methods

### Genome sequencing and assembly

To capture the genetic diversity from all ecotypes of rice, we collected 11 accessions of weedy rice from seven countries based on their phylogeny with cultivated rice and geographic positions; we also included one XI cultivar NJ11, or Nanjing 11, which is presumed to be the direct ancestor of weedy rice in the Yangtze River Basin (Qiu et al., 2020). Genomic DNA samples were extracted from young leaves of the 12 rice accessions, and their genomes were sequenced by PacBio HiFi mode according to the instructions from the manufacturer. For each accession, the sequencing depth of HiFi subreads ranged from 21.6× for accession 13-65 to 45.0× for accession YZ-2, with an average of 32.4×. Following the standard protocol, Hi-C libraries of four accessions (NJ11, CX20, YCW03 and 18XHB-83) were constructed using fresh young leaves digested with the 4-cutter restriction enzyme MboI. Hi-C libraries were sequenced on an Illumina HiSeq 4000 platform with 2×150-bp paired reads.

The genomes were first assembled using hifiasm (v0.15.1-r334, default parameters) (Cheng et al., 2020). For each accession, HiFi subreads were mapped against the corresponding assembly using minimap2 (Li, 2018), and Purge_dups was applied to purge duplicates and remove redundant sequences according to the mapping depth (Guan et al., 2020). We further used Racon (v1.4.0) to polish the assemblies with HiFi subreads for three rounds under default parameters (Vaser et al., 2017). Contigs less than 10 kb were removed from the final version. For each accession, contigs were anchored into pseudochromosomes by using a reference-guiding approach RaGOO (Alonge et al., 2019). By aligning contigs against the Nipponbare assembly using minimap2, the contigs were ordered and oriented along 12 chromosomes with no further chimeric splitting.

Previously released rice assemblies based on third-generation sequencing platforms were collected, including PacBio (Du et al., 2017; Sun et al., 2019; Wang et al., 2019; Ma et al., 2020; Xie et al., 2020; Zhou et al., 2020; Qin et al., 2021; Song et al., 2021) and Nanopore sequencing (Choi et al., 2020; Read et al., 2020; Shang et al., 2022; Zhang et al., 2022). Before adopting assemblies in the construction of the rice pangenome, we first systematically evaluated the assembly qualities and ruled out assemblies that did not meet our criteria.

### Quality assessment of rice genome assemblies

We first assessed the quality of our 12 newly assembled rice genomes. Synteny against the reference assembly Nipponbare and gapless assembly MH63 (Song et al., 2021) confirmed their high completeness. The paired Hi-C reads of four accessions (YCW03, 18XHB-83, CX20 and NJ11) were cleaned using NGSQC toolkit (Patel and Jain, 2012) and then mapped to the corresponding assembly using Bowtie2 (v2.3.5.1) (Langmead and Salzberg, 2012). After retaining high-quality and validated paired reads (mapping quality ≥ 30, edit distance ≤ 5, number of mismatches in the alignment ≤ 3, number of gap opens ≤ 2 and number of gap extensions ≤ 2), chromosome interaction maps were plotted by AllHiC_plot (Zhang et al., 2020), and they revealed high accuracy in contig ordering and orientation (Supplementary Fig. 1).

Genome assemblies of *Oryza sativa* and *Oryza rufipogon* based on third-generation sequencing (through May 2022) were collected (Supplementary Table 1). DXCWR (Ma et al., 2020) was excluded considering its low contig N50 of less than 200 kb. IRGC109232 (Zhao et al., 2018) was removed due to the abnormal size of the assembly obtained from the public database. Eleven assemblies in Zhang et al. (2022) were randomly selected from 75 newly generated genomes and used in subsequent quality assessment. Five indices were applied to evaluate the genome quality of all rice assemblies (Supplementary Fig. 2a). Assembly continuity was evaluated by contig N50 and the long-terminal repeat assembly index (LAI), which was revealed by the assembly completeness of long-terminal-repeat (LTR) retrotransposons (Ou et al., 2018). The LTR elements of each assembly were identified by RepeatMasker and RepeatModeler (http://repeatmasker.org/). BUSCO (v4.1.2) metrics were calculated to evaluate the completeness by using dataset poales_odb10 containing 4896 genes (Simao et al., 2015). As expected, assemblies based on long-read sequencing performed well on the above quality indices (Supplementary Fig. 2a). Hence, in addition to the above evaluations at the whole-genome level, base-resolution accuracy and completeness were measured by the consensus quality value (QV) and the number of homozygous variants called by self-short-read mapping. Reference-free QVs were calculated by Merqury (Rhie et al., 2020) and yak (https://github.com/lh3/yak) by comparing *k*-mers derived from unassembled, high-accuracy sequencing reads to a genome assembly. Homozygous variants (SNPs and InDels) called with short reads by self-mapping are regarded as potential assembly errors. Raw short-read data were first cleaned by NGSQC-toolkit and mapped against the corresponding assembly by Bowtie2. Variants were detected using GATK (v3.7, default parameters) (McKenna et al., 2010) and annotated by SnpEff (v3.6) to profile their potential effects on the prediction of amino acid sequences and further gene functions (Cingolani et al., 2012). Variants with high effects (including stop loss and gain, start loss, frame-shift variant) and moderate effects (including in-frame insertion/deletion, missense variant) will directly impact the reliability of gene haplotypes and downstream haplotype analysis. The assembly quality of wild accessions from Shang et al. (2022) and one weedy accession (YCW03) were not assessed at the base level due to the unavailability of NGS data. Assemblies with low base-level quality were removed for cultivated rice, mainly including Nanopore sequencing-based genomes, except three accessions (DomSufid, Basmati334 and JHU) from aromatic and tropical groups of GJ (*O. sativa* ssp. *japonica*), which were kept to balance the sampling of each group. For wild genomes, only two accessions, W2014 from Ma et al. (2020) and IRGC106162 from Xie et al. (2020), sequenced using the PacBio platform were available. Thus, nine assemblies in different wild groups from Shang et al. (2022) sequenced by the Nanopore platform were adopted for analysis.

### Phylogenetic relationship of rice assemblies

Together with the 12 new assemblies in this study, a total of 75 rice assemblies, including 11 wild accessions (*O. rufipogon*), 12 weedy accessions (*O. sativa* ssp. *spontanea*), 51 cultivated accessions (*O. sativa*) and an African cultivated rice outgroup (*Oryza glaberrima*) accession CG14, were used in this study. We first confirmed their phylogenetic relationship and assigned them to taxonomic groups using whole-genome SNPs. The assemblies were aligned against the reference assembly Nipponbare using nucmer implemented in the MUMmer package (v4.0.0) (Marçais et al., 2018), and called SNPs were used to build their phylogeny by FastTreeMP with 1000 bootstrap replicates (Price et al., 2009). According to their phylogeny and prior knowledge of the 74 *Oryza sativa* and *Oryza rufipogon* accessions, each accession was assigned to groups. Briefly, the subspecies GJ (*O. sativa* ssp. *japonica*, *n* = 23 in total) includes four *aromatic* (aro), four tropical (trp), two subtropical (subtrp) and 13 temperate (tmp) accessions; subspecies XI (*O. sativa* ssp. *indica*, *n* = 40) includes four *aus*, three XI2, seven XI3, ten XI1A and 16 XI1B accessions. The wild population (*O. rufipogon*, *n* = 11) includes two accessions from Or-3, four from Or-2, three from Or-1, and two from Or-4. Genotype data for approximately seven thousand rice accessions used in the principal component analysis (PCA) were adopted from a previous study (Wu et al., 2022a). PCA and IBD (Identity-by-descent) calculation were performed using Plink (v1.9) with a pruned subset of SNPs based on linkage disequilibrium (10 SNPs in each 50-kb sliding window with pairwise Pearson’s correlation efficient *r*^2^ less than 0.5) (Chang et al., 2015).

### Genome annotation

In total, 75 rice assemblies (including African rice CG14 as an outgroup) were annotated in a unified pipeline. First, transposon elements (TEs) were identified by using the Extensive *de novo* TE Annotator (EDTA) approach (https://github.com/oushujun/EDTA) (Su et al., 2021). For each accession, gene models were predicted on the repeat-masked genome using an approach integrating *ab initio* predictions and homology-based prediction. For *ab initio* prediction, Augustus (Stanke et al., 2006) and Fgenesh (Salamov and Solovyev, 2000) were performed with default parameters. For homology-based prediction, previously predicted protein sequences of the Nipponbare (IRGSP v1.0), gapless MH63RS3 and ZS97RS3 assemblies (Song et al., 2021) and over 30 other well-annotated assemblies (Qin et al., 2021) were used to search putative protein-coding gene models with GMAP for each accession (Wu and Watanabe, 2005). The predictions were integrated into non-redundant consensus gene models using EVidenceModeler (v1.1.1) (Haas et al., 2008). Short gene models (less than 50 amino acids) and gene models with homology to sequences in Repbase (*e*-value ≤ 1e−5, identity ≥ 30%, coverage ≥ 25%) were further removed from the final annotation. The protein domains of all predicted coding gene models were inferred using InterProScan (v5.24-63.0) (Zdobnov and Apweiler, 2001).

### Pan-genome construction using MCL

A Markov Clustering (MCL) approach OrthoFinder (v2.4.1) was applied to cluster all the predicted gene models of 74 rice genomes (African rice accession CG14 was excluded) with default parameters (diamond all-versus-all *e*-value < 1e-5 and inflation parameter = 1.5) (Emms and Kelly, 2019). Finally, over 3.10 million predicted genes were clustered into 67,080 orthogroups (OGs). Increasing the inflation parameter can be used to achieve higher precision at the cost of lower recall and have a larger size of OGs. Conversely, a smaller value of the inflation parameter could achieve higher recall at the cost of lower precision and produce smaller sized OGs, which easily clusters paralogs together (Emms and Kelly, 2019). Thus, three additional inflation parameters (I = 1.8, 2.0 and 2.5) were set to profile the effects on clustering (Supplementary Fig. 9). Clustered gene families were categorized into core (present in all genomes, *n* = 74), soft-core (present in at least 90% of genomes, *n* = 67 to 73), dispensable (present in more than one but less than 90% of genomes, *n* = 2 to 66) and private (only present in one genome) on the basis of the number of rice accessions in which they were identified.

### Pan-genome construction using synteny

Given that the MCL approach does not efficiently and accurately distinguish paralogs from orthologs with high sequence similarity, we utilized the availability of genomic coordinates of gene models to build the synteny-based pan-genome. Pairwise all-to-all alignments using protein sequences were performed for all 74 genomes. Aligner Diamond runs much faster for large protein sequence data than BLASTP (Buchfink et al., 2021). We compared the recall performance between BLASTP and Diamond in detecting syntelogs between genomes (Supplementary Fig. 6). Diamond was run for each pair under three modes (default, sensitive and ultra-sensitive). The alignments were filtered to keep only the best hits, and DAGchainer (Haas et al., 2004) was used to detect syntenic genomic regions and syntelogs (parameters: -Z 12 -D 200000 -g 1 -A 5). No differences were observed among the three Diamond modes in the number of detected syntelogs (Supplementary Fig. 6). BLASTP was performed under default parameters, and syntelogs were identified using the same pipeline. The BLASTP approach searched more syntelogs, and thus, the syntelogs were used in downstream pan-genome construction (Supplementary Fig. 6). Instead of a reciprocal best hit search, the roles of reference and query in a pairwise BLASTP search sometimes result in differences due to individual-specific tandem duplicates, and such individual-specific tandem duplicates were considered as a single gene in the final pan-genome. Pairwise syntelog information was merged together with Nipponbare as an initial framework one by one using SynPan (https://github.com/dongyawu/PangenomeEvolution). If a gene from an additional genome was syntenic to a previously merged pan-genome, this gene was assigned to an existing SG. If a gene from an additional genome was not syntenic to any gene in the merged iterative pan-genome, a new SG was created. In total, 74 genomes were merged together as a synteny-based pan-genome, including 175,528 SGs. The SGs were further categorized as core, soft-core, dispensable and private following the criteria used in OG categorization.

### Construction of the rice NLRome

To capture the diversity of NLRs in rice, we integrated multiple software predictions and gene synteny in rice genomes to obtain a comprehensive and complete rice NLRome. The NB-ARC domain was first predicted using hmmsearch (HMMER v3.1b2) against the Pfam database (v30.0) with a threshold *e*-value less than 1e-5. The LRR domains were predicted with NLR-parser (v3.0) (Steuernagel et al., 2015) by searching for motifs 9, 11 and 19; the coil domains were predicted by searching for motifs 16 and 17; and the TIR domains were predicted by searching for motifs 13, 15 and 18. All putative NLR types from genome-wide protein sequences were also determined using RGAugury (Li et al., 2016). After the NLR genes for each genome were identified by domain prediction, the NLR genes were mapped back to the SG-based pangenome to involve NLR syntelogs lacking canonical domains and find a more comprehensive and extensive NLR inventory. NLR genes were defined to have at least one NB-ARC, TIR, or CCR (RPW8) canonical domain. LRR or CC motifs alone were not considered sufficient for NLR identification. Finally, NLRs in rice genomes were identified and categorized as CNLs (containing CC, NB-ARC, and LRR domains), CNs (containing CC and NB-ARC domains) and NLs (containing only canonical NB-ARC domains). Noncanonical architectures of some NLRs have additional integrated domains (IDs), while canonical architectures contain only NB-ARC (Pfam accession PF00931), TIR (PF01582), RPW8 (PF05659), or LRR (PF00560, PF07725, PF13306, PF13855) domains or CC motifs. For ID identification, we used the genome protein sequences as the input for InterProScan, and annotations were processed with in-house scripts to obtain ID information. There are currently several known cases in which two NLR genes are required to affect the resistance function, with one protein in the pair acting in effector recognition and the other acting in signaling activation. Since all known functional pairs are present in head-to-head arrangement in the rice genome, we identified head-to-head NLR pairs by searching for NLR genes near each other and no more than 10 kb away. Multiple sequence alignment was performed using coding sequences by MAFFT (v7.490) (Katoh & Standley, 2013), and the nucleotide diversity and Tajima’s *D* value for all members in each SG were calculated using DnaSP (v6) (Rozas et al., 2017).

### Haplotype diversity and ancestral haplotype assignment

We used the average pairwise haplotype difference to measure the haplotype diversity in one population and the divergence between two populations (Supplementary Fig. 13a). The haplotype number (defined as N100) for each SG was determined by counting the unique sequences of all predicted proteins. To exclude rare haplotypes, haplotype N90 was defined as the least haplotype number that needs to be included for covering 90% of sequences in each SG. To understand the haplotype-aware origin of genes from different rice groups, we inferred the ancestral haplotype composition for each SG by determining the dominant haplotype in each rice group and assigning ancestral haplotype IDs for all genes in a group priority-based referring strategy (Supplementary Fig. 13b). Group information is prior based on whole-genome phylogeny or population structure. We labeled five haplotype IDs (hapI to hapV) to represent the dominant haplotypes of groups tmp, XI1A, XI1B, *aus* and XI3 in order, presented by red, blue, orange, yellow, and green in Fig. 4a and Supplementary Fig. 15, respectively. HapI was first defined as the most dominant sequence in group tmp. If the dominant haplotype in the current group was defined by a former group, the haplotype ID was skipped and set as missing. For example, if the dominant haplotype in XI1A was the same as that in tmp (hapI), hapII was then not defined. The frequency of a dominant haplotype within a group should be at least three. All other rare haplotypes were compressed as hapR in gray to simplify the ancestral haplotype graphs. Gene absence is represented by white blocks. Different group priority orders have no influences on the calculation of haplotype diversity and divergence but only change the ancestral haplotype graphs (Supplementary Fig. 13c).

### Inter-subspecies introgression blocks

As observed from the ancestral haplotype landscape, haplotypes in some large genomic regions were shared between XI and GJ. We used inter-population haplotype divergence (HDG) to quantify haplotype sharing by calculating the average differences in haplotypes from the two populations (Supplementary Fig. 13a). We merged adjacent SGs whose divergence values between tmp and XI (excluding *aus*) were less than 0.5 into lowly divergent blocks between subspecies as candidate introgression blocks. At least 10 SGs were required within a single introgression block. To examine the significance of the nonrandom clustered distribution of these lowly divergent SGs in blocks, we randomly sampled the same number of SGs as lowly divergent SGs observed on each chromosome and calculated the density of sampled SGs in sliding windows of every 10 SGs. A total of 100 thousand random samplings were replicated, and the *P* values were determined by counting the sampling times where the density of sampled SGs was higher than that observed for each window. *P* = 0.01 was empirically set as a cutoff value. Assuming that the low divergence in the detected blocks was caused by inter-subspecies introgression after their divergence, the divergence time between GJ and XI should be younger in the identified blocks than in their neighboring regions. We used the synonymous substitution rate (*K*s) to measure the relative divergence time between GJ and XI, free from selection. Significantly lower *K*s values were observed in the majority of 18 large blocks (>300 kb) than in their flanking regions (Supplementary Fig. 16).

### Phylogeny of introgression blocks

To investigate the origin and gene flow of the 73 candidate introgression blocks, we utilized recently released genomic sequences of 184 wild accessions with high sequencing depth (8× on average, much higher than <2× on average in a previously used wild population)(Zheng et al., 2022; Huang et al., 2012). Raw sequencing reads were first cleaned using NGSQC toolkit and mapped against the reference assembly Nipponbare (IRGSP v1.0) using Bowtie2. Assemblies of two wild rice accessions (W1943 and DWCWR) from group Or-3 were added. Combing the 77 assemblies and 184 wild genomes, high-quality SNPs were called following a previous pipeline (Qiu et al., 2020). The population structure was first surveyed by PCA using Plink (v1.9) (Chang et al., 2015) and the phylogeny was produced by FastTreeMP with 1000 bootstrap replicates based on 6.85 million high-quality SNPs (minor allele frequency of 0.02 and maximum missing rate of 0.1). Four wild groups were identified: Or-1, Or-2, Or-3 and Or-4. The SNPs in each introgression block were extracted and used to build the phylogeny using IQ-TREE (v1.6.12) with the best substitution model TIM2e+R2 determined by ModelFinder implemented in IQ-TREE (Nguyen et al., 2015) and FastTreeMP with 1000 bootstrap replications, where the African cultivated rice accession CG14 was set as the outgroup. To avoid over-interpretation, the phylogeny in which the wild accessions were not obviously and empirically clustered into four groups, as defined by the whole-genome SNPs, was discarded.

### ABBA-BABA test

The availability of population-level whole-genome high-depth sequencing data of wild rice from four groups (Or-1, Or-2, Or-3 and Or-4), enables us to perform comprehensive statistical *D* tests, which is widely used and robust in gene flow detection (Green et al., 2010; Wu et al., 2022b). We randomly selected genomes of XI1A rice accessions (*n* = 100), XI1B (*n* = 100), XI2 (*n* = 80), XI3 (*n* = 100) and *aus* (*n* = 60) from 3K RGP (Wang et al., 2018). We employed *f*d statistic to indicate the introgression from GJ to XI in sliding genomic windows (Martin et al., 2015). Under a given quartet topology ((P1, P2), P3, O), positive *f*d statistic values indicate the introgression from P3 to P2, zero represents no introgression, and negative *f*d statistic values have no biological meaning and thus are converted to zero. We estimated the *f*d statistic values under topology ((Or-1, X), tmp, Or-4), where X is Or-2, XI1A, XI1B, XI2, XI3 and *aus* in topology T1 to T6, respectively (Supplementary Fig. 18). T1 is set as a background control in detecting introgression. To eliminate the influence of modern breeding where inter-subspecies hybridization is frequently performed on ancient introgression inference, we used genomes of only landraces in each group to repeat the *f*d calculation, where tmp, XI1A, XI2, XI3, and *aus* included 47, 24, 21, 52 and 32 landrace accessions. Generally, no obvious differences are observed between introgression block determination using all and landrace accessions only (Supplementary Fig. 19). Python scripts are available at https://github.com/simonhmartin/genomics_general. Parameters are set as “window size: 20 kb, step size: 2 kb, minimum good sites per window: 50, and minimum proportion of samples genotyped per site: 0.4”. The final putative introgression regions were determined by integrating evidences form haplotype divergence, phylogeny and ABBA-BABA tests. The functional enrichment analysis was performed using ShinyGO (v0.77) (Ge et al., 2020).

### Structural variations in de-domestication

To gain a more detailed understanding of structural variations in de-domestication, we compared the genome assemblies of weedy and cultivated rice. It has been found that the sensitivity of detecting deletions is higher than that of insertions, and we adopted a pairwise genome alignment strategy as mentioned previously in Jayakodi et al. (2020). Each pair contains a weedy accession and a cultivar accession, which compose a monophyly in the phylogenetic tree, and the genome assembly of cultivated rice was considered a query or reference genome. Nucmer in the MUMmer package was used to obtain the results of these two alignments (Marçais et al., 2018), and then PAVs (presence-and-absence variants, including insertions and deletions) were called using Assemblytics (v.1.2.1) (Nattestad & Schatz, 2016). The structural PAVs were classified as InDels (<50 bp) and SVs (≥ 50 bp). Only deletions were kept in both alignments and converted into PAVs according to the reference genome in each pair. Genes located in or intersected with each PAV were obtained, as well as corresponding gene annotations. The variations in domestication genes and their flanking regions (15 kb for *qSH1* and 2 kb for other genes) were manually investigated, and the causative mutations during the domestication process were checked. To validate the reliability of structural variations in domestication or improvement genes (e.g., *OsC1* and *Bh4*), HiFi subreads of weedy rice were mapped against themselves and corresponding cultivated assemblies to check the local alignments using IGV (Thorvaldsdóttir et al., 2013). To infer the source of the causative mutation in *Bh4* in weedy accessions, the *Bh4* phylogeny based on SNPs was analyzed.

The 73 rice genome assemblies were aligned against the Nipponbare reference genome using the nucmer program implemented in the MUMmer package with the default parameter, and only the best position of each query on the reference was preserved. The alignments from 6.0 to 6.5 Mb on chromosome 7 in the Nipponbare assembly were extracted for visualization by synteny plots and comparison among wild, cultivated and weedy assemblies. To validate the candidate introgression event inferred from the synteny plots, SNPs around the *Rc* gene (including its flanking 2-kb regions) were extracted and used in phylogeny construction by FastTreeMP under the GTR+CAT model with 1000-times bootstrapping.

### Data and code availability

All the PacBio HiFi subreads for 12 rice accessions and Hi-C data for four rice accessions generated in this study have been deposited in NGDC (https://ngdc.cncb.ac.cn/) under the accession code PRJCA012143. The newly generated assemblies for 12 accessions and the annotations (including GFF, CDS sequences and predicted protein sequences) for all 74 rice accessions can be found under project accession PRJCA012309 in NGDC. The raw resequencing data of previously published wild accessions can be downloaded from NCBI under accession number PRJNA657701. The VCF file of SNPs from all 75 assemblies and an additional 184 wild rice accessions with high sequencing depth is available at Zenodo (10.5281/zenodo.7196576). The gene re-annotations, NLR annotations and Pfam annotations for all 75 rice accessions are deposited at Zenodo (10.5281/zenodo.7248110). The in-house scripts used in this study have been deposited in GitHub (https://github.com/dongyawu/PangenomeEvolution).

## Acknowledgements

We thank Peng Qin (Sichuan Agricultural University) and Jian Sun (Shenyang Agricultural University) for providing rice seeds. We thank Lianguang Shang (AGIS, Chinese Academy of Agricultural Sciences) for providing the genome assemblies of nine wild rice accessions. We thank Jie Qiu (Shanghai Normal University) for his constructive suggestions. This work is supported by National Natural Science Foundation of China (31971865) and Department of Science and Technology of Zhejiang Province (2022C02032 and 2020C02002) to L.F. and China National Postdoctoral Program for Innovative Talents (BX20220269) to D.W.

## Author contributions

L.F. and Q.Q. conceived and supervised the study. D.W., L.J. and C.D. assembled and annotated the genomes. D.W. and L.J. collected previously released assemblies of rice genomes. Y.H. and L.X. evaluated the quality of rice assemblies. D.W., L.X. and Y.H. constructed the rice syntelog-based pan-genome and analyzed the ancestral haplotypes. L.X. carried out the analysis of NLR genes. Y.S. and L.X. performed the analysis of structural variations. L.F., C.Y. and Q.Q. discussed the results. D.W., Y.S. and L.X. wrote the manuscript and L.F. and C.Y. revised it. All authors discussed the results and commented on the manuscript.

## Competing interests

The authors declare no competing interests.

## Additional information

The supplementary material is available online.

## Supplementary Information

**Supplementary Table 1.** Meta information of 75 rice genome assemblies used in this study.

**Supplementary Table 2.** Haplotype divergence between XI and GJ for domestication and improvement genes

**Supplementary Table 3.** Putative introgression blocks identified by haplotype divergence between GJ and XI and validation by phylogeny and ABBA-BABA tests

**Supplementary Table 4.** Gene functional enrichment in final putative introgression blocks

**Supplementary Table 5.** Cloned genes in the putative introgression blocks

**Supplementary Table 6.** Structural variations between each pair of weedy and cultivated accessions in agronomy-related genes

**Supplementary Fig. 1** Hi-C interaction heatmaps for each chromosome from four rice accessions (a, YCW03; b, 18XHB-83; c, CX20; d, NJ11).

**Supplementary Fig. 2** Statistical information of the rice genomes used in this study. Assembly (size), annotation (gene number, TE size and proportion) and quality assessment (contig N50, BUSCO and LAI) of rice genomes. (b) Differences in genomic features between subspecies GJ and XI. In the boxplots, the horizontal line shows the median value, and the whiskers show the 25% and 75% quartile values of each genomic feature. *P* values were calculated by Student’s *t* test.

**Supplementary Fig. 3** Dot plots of newly generated rice assemblies in this study against the reference assembly Nipponbare (IRGSP) and gapless assembly MH63RS3.

**Supplementary Fig. 4** Assembly quality assessment in base accuracy. (a) QVs for rice genomes in four rice pan-genome projects using yak (https://github.com/lh3/yak). The number and annotation to SNPs and InDels for each genome by mapping NGS short reads against their own assembly.

**Supplementary Fig. 5** PCA reveals the representativeness and diversity of genome assemblies used in this study. The first two principle components are shown. Filled circles indicate the assemblies used in this study. Green, blue and red represent wild, cultivated and weedy assemblies, respectively.

**Supplementary Fig. 6** Performance of BLASTP and Diamond (under different modes) in syntelog identification. (a) The BLASTP approach identifies more syntelogs than Diamond. (b) Syntelog number between accession 02428 and other accessions identified using BLASTP and Diamond.

**Supplementary Fig. 7** Pairwise synteny reveals evolutionary signatures in groups and individuals. (a) Syntelog numbers between rice groups. (b) Syntelog numbers between each accession and other accessions from the GJ and XI (including *aus*) subspecies. In the boxplots, the horizontal line shows the median value, and the whiskers show the 25% and 75% quartile values of syntelog numbers.

**Supplementary Fig. 8** Comparison of MCL and synteny-based clustering. (a) A brief scheme illustrating MCL ortholog clustering and syntelog clustering. (b) Group size comparison using synteny-based clustering (SynPan) and MCL clustering (OrthoFinder).

**Supplementary Fig. 9** Benchmarking analysis on the influences of inflation parameters in the MCL clustering in rice genomes. (a) The OG numbers shared by different sizes of rice genomes under different inflation parameters. (b) Gene counts in OGs shared by different sizes of genomes. (c) Average gene number per OG under different inflation parameters.

**Supplementary Fig. 10** Composition and features of the rice syntelog-based pangenome. (a) Pan-gene composition (core, soft-core, dispensable and private) of the rice pan-genome. (b) Proportions of domain-annotated genes in four categories of the rice pan-genome. (c) Percentage of SGs (blue) and genes (red) with domain gain-and-loss.

**Supplementary Fig. 11** Comparison of NLR genes in rice genomes from different ecotypes and subspecies. (a) Distribution of different NLR genes in wild, cultivated and weedy accessions. (b) Distribution of different NLR genes in wild rice, XI and GJ. Clustered NLRs in wild rice, XI and GJ. *P* values are calculated using Wilcoxon test. (d) Relationship between genome assembly completeness (as indicated by LAI) and NLR size. In the boxplots, the horizontal line shows the median value, and the whiskers show the 25% and 75% quartile values of NLR sizes.

**Supplementary Fig. 12** Genomic features of NLRs in rice genomes. (a) Dynamics of domain architectures in the rice NLRome. CNL, NL and null (no canonical architectures) are defined as three NLR architecture types in rice. Multiple types are observed for most SGs. (b) Nucleotide diversity and selection of NLRs from core, soft-core and dispensable SGs. In the boxplots, the horizontal line shows the median value, and the whiskers show the 25% and 75% quartile values of Pi and Tajima’s *D*. Significance *P* values are performed using Wilcoxon test. *, *P* < 0.05.

**Supplementary Fig. 13** Haplotype analysis on rice syntelogs. (a) Definition of haplotype diversity and divergence. Haplotype diversity and divergence represent average haplotype differences among sequences in a syntelog group within one group and among two groups, respectively, where Xi and Xj are the presence count of haplotype i and haplotype j in group X, and ∑X is the total sequence count within a syntelog group. (b) Brief scheme illustrating the assignment and visualization of ancestral haplotypes for each genome. A group priority-based referring strategy is used to assign ancestral haplotypes. Group information is prior based on whole-genome phylogeny or population structure. The most dominant sequence in syntelogs from Group 1 is set as hapI. If the most dominant sequence from Group 2 is not HapI, then define HapII, otherwise HapII is skipped. If the most dominant sequence from Group 3 is neither HapI nor HapII, then define HapIII, otherwise HapIII is skipped. By analogy, dominant haplotypes are determined and colored for each syntelog group. Rare haplotypes are named as HapR colored by dark gray. In this study, the group priority order is set as tmp > XI1A > XI1B > *aus* > XI3. Different orders have no influences on the calculation of haplotype diversity and divergence. (c) mosaic graphs of ancestral haplotypes in chromosome 1 across 74 rice genomes with different priority orders, tmp > XI1A > XI1B > *aus* > XI3, XI1B > XI1A > tmp > *aus* > XI3 and XI1A > XI1B > tmp > *aus* > XI3. For each window, the same color indicates the same haplotype and dark and light gray indicates rare haplotypes and syntelog absence.

**Supplementary Fig. 14** Genetic diversity measured by full-length gene, coding region, and predicted protein sequences in rice genomes. (a) Comparison of diversity measured by haplotype N100, N90 and haplotype diversity using full-length nucleotide sequences, coding sequences and predicted protein sequences. In the boxplots, the horizontal line shows the median value, and the whiskers show the 25% and 75% quartile values of each diversity indice. (b) Haplotype diversity in GJ and XI, compared to all rice genomes.

**Supplementary Fig. 15** Ancestral haplotype landscape on chromosome 1 to chromosome 12. Putative introgression blocks are shown in gray rectangles and numbered. Haplotype divergence and *P* values (scaled by −log10) of non-random distribution significance tests are shown along each chromosome. Red dashed lines represent thresholds to determine introgression blocks.

**Supplementary Fig. 16** Synonymous substitution rates (*K*s) of genes in putative introgression blocks and their neighboring left and right regions. Three replicates (a, b, and c) between the GJ and XI genomes were performed. In the boxplots, the horizontal line shows the median value, and the whiskers show the 25% and 75% quartile values of *K*s. *P* values are calculated using Wilcoxon test.

**Supplementary Fig. 17** Population structure of wild rice accessions used in this study. Phylogenetic tree of *Oryza rufipogon* and *Oryza sativa*. Four wild groups (Or-1, Or-2, Or-3 and Or-4) are indicated in different colors. (b) PCA plots of the first three principle components, where “W”, “J” and “I” represent wild, GJ and XI accessions. (c) Geographical sources of wild accessions in different groups used in this study.

**Supplementary Fig. 18** Introgression *f*d distributions of ABBA-BABA test on chromosome 1 to chromosome 12 in topology T1 to T6, where P1 is Or-1 (*n* = 37), P3 is tmp (GJ, *n* = 100), O/outgroup is Or-4 (*n* = 25), and P2 was set as Or-2 (*n* = 42), XI1A (*n* = 100), XI1B (*n* = 100), XI2 (*n* = 80), XI3 (*n* = 100) and *aus* (*n*=60), respectively. T1 was set as a background control in introgression detection. Genomic positions of putative introgression regions are indicated by gray rectangles and detailed coordinates are provided in Supplementary Table 3. Blocks larger than 300kb are highlighted in red and blocks not supported by *f*d are underlined.

**Supplementary Fig. 19** Comparison of *f*d using all genomes and landraces only under different topologies on chromosome 1. (a) *f*d distribution along chromosome 1. Group tmp, XI1A, XI2, XI3, and *aus* include 47, 24, 21, 52 and 32 landrace accessions, respectively. (b) Comparison of *f*d on chromosome 1 in T2 *vs* T12, T4 *vs* T14, and T5 *vs* T15.

**Supplementary Fig. 20** Genome synteny between each weedy rice assembly and the corresponding phylogenetically closest cultivated rice assembly.

**Supplementary Fig. 21** Summary of structural variations between weedy and cultivated rice genomes.

**Supplementary Fig. 22** Structural variations (>50 bp) between assemblies of cultivar accession NJ11 and weedy rice accession CX20 on 12 chromosomes. The largest six SVs (numbered from 1 to 6) are zoomed in and annotated, including four insertions (1, 2, 5 and 6) and two deletions (3 and 4) in CX20.

**Supplementary Fig. 23** The Integrative Genomics Viewer (IGV) snapshots show the structural variations in *OsC1* and *Bh4* between weedy and cultivated rice.

**Supplementary Fig. 24** Phylogeny of *Bh4* in rice and morphology of rice seed hulls. Bootstrap values less than 0.90 are indicated on branches.

## Notes

### Competing Interest Statement

The authors have declared no competing interest.

