## Supplementary Figures for "A syntelog-based pan-genome provides insights into rice domestication and de-domestication"

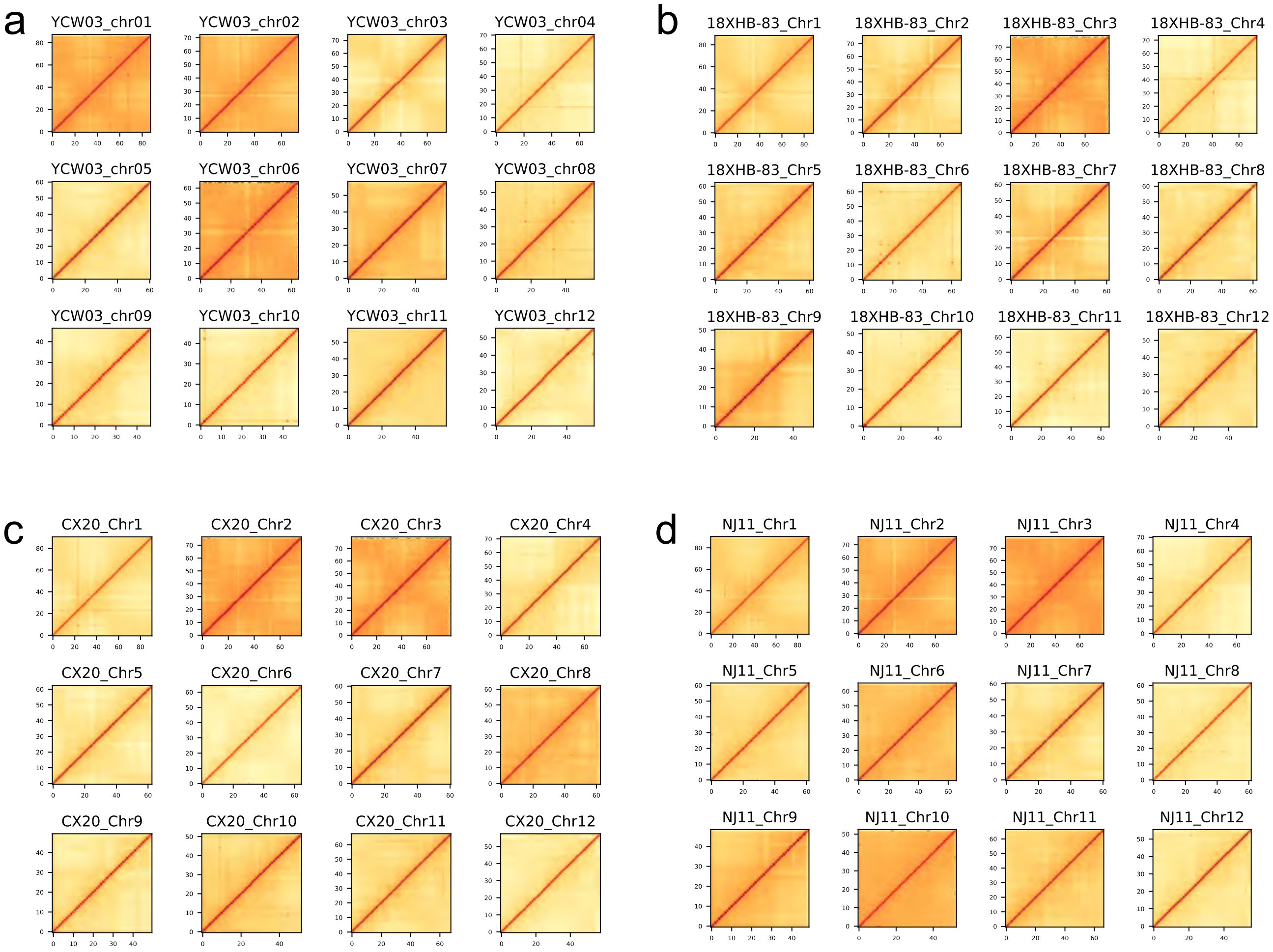


**Supplemenatry Fig. 1** Hi-C interaction heat-maps for each chromosome of four rice accessions (**a**, YCW03; **b**, 18XHB-83; **c**, CX20; **d**, NJ11).

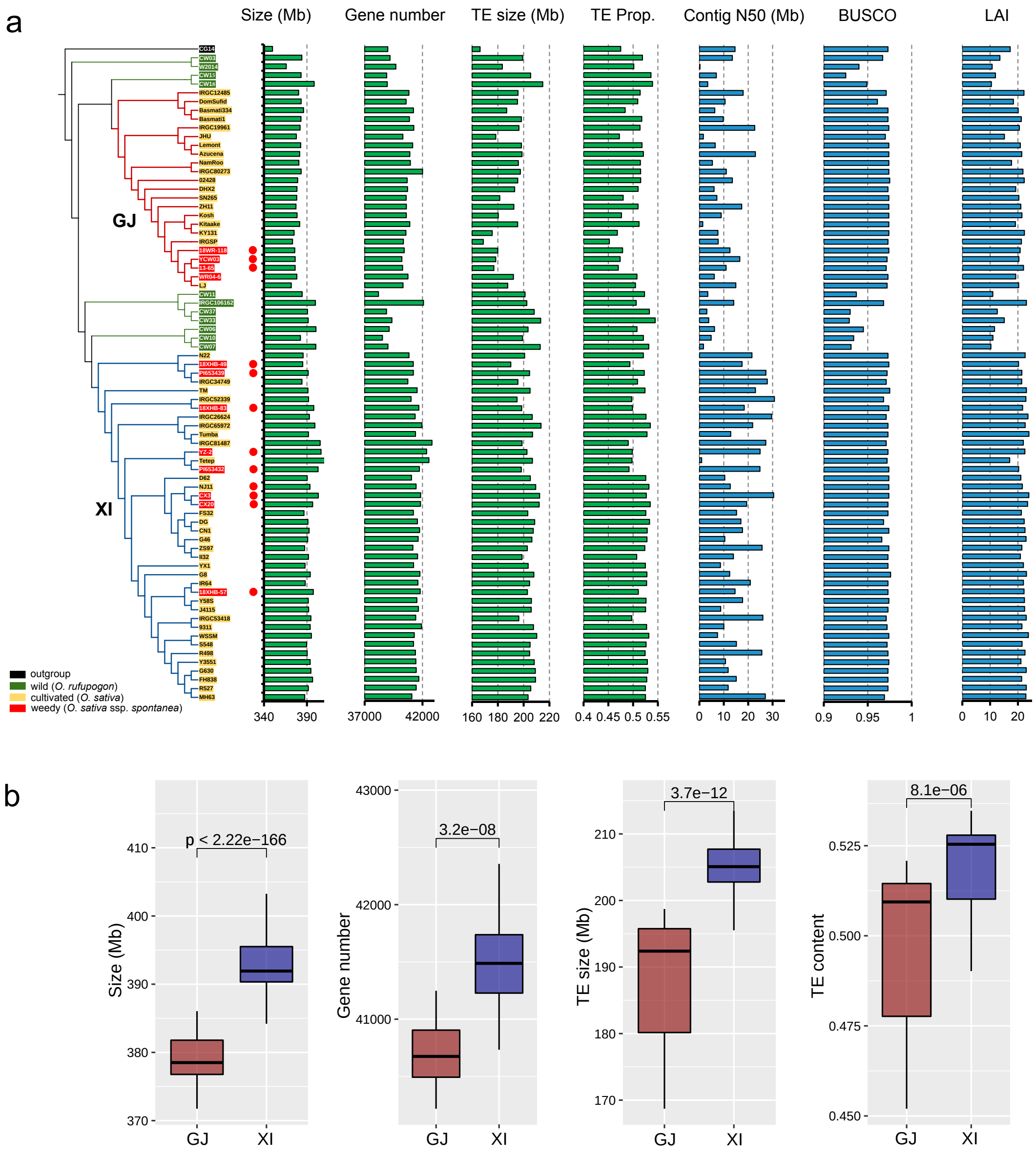


**Supplementary Fig. 2** Statistical information of rice genomes used in this study. (**a**) assembly (size), annotation (gene number, TE size and proportion) and quality assessment (Contig N50, BUSCO and LAI) of rice genomes. (**b**) Differences in genomic features between subspecies GJ and XI. In the boxplots, the horizontal line shows the median value, and the whiskers show the 25% and 75% quartile values of each genomic feature. *P* values are calculated by Student's *t* test.

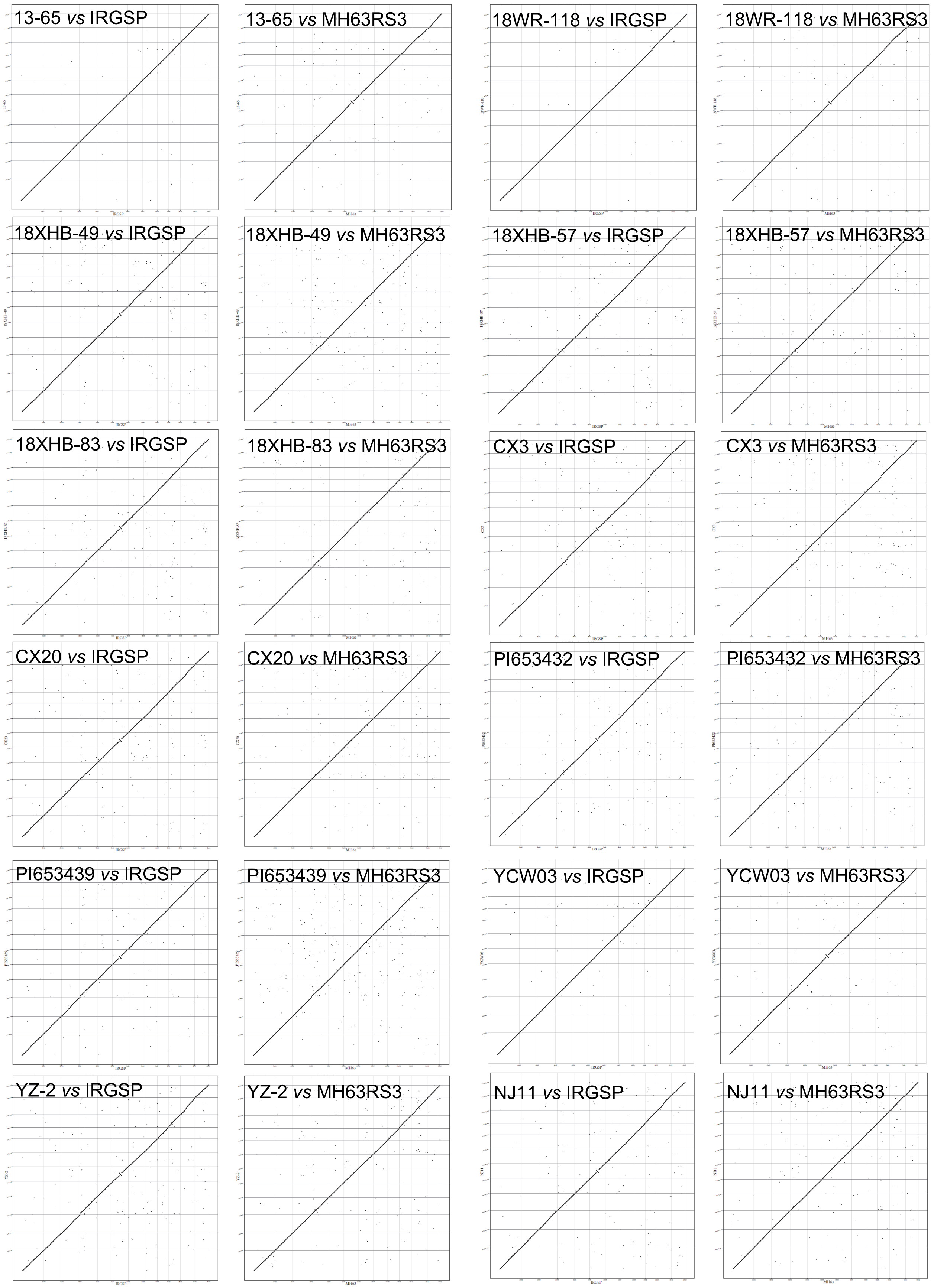


**Supplementary Fig. 3** Dot plots of newly generated rice assemblies in this study against reference assembly Nipponbare (IRGSP) and gapless assembly MH63RS3.

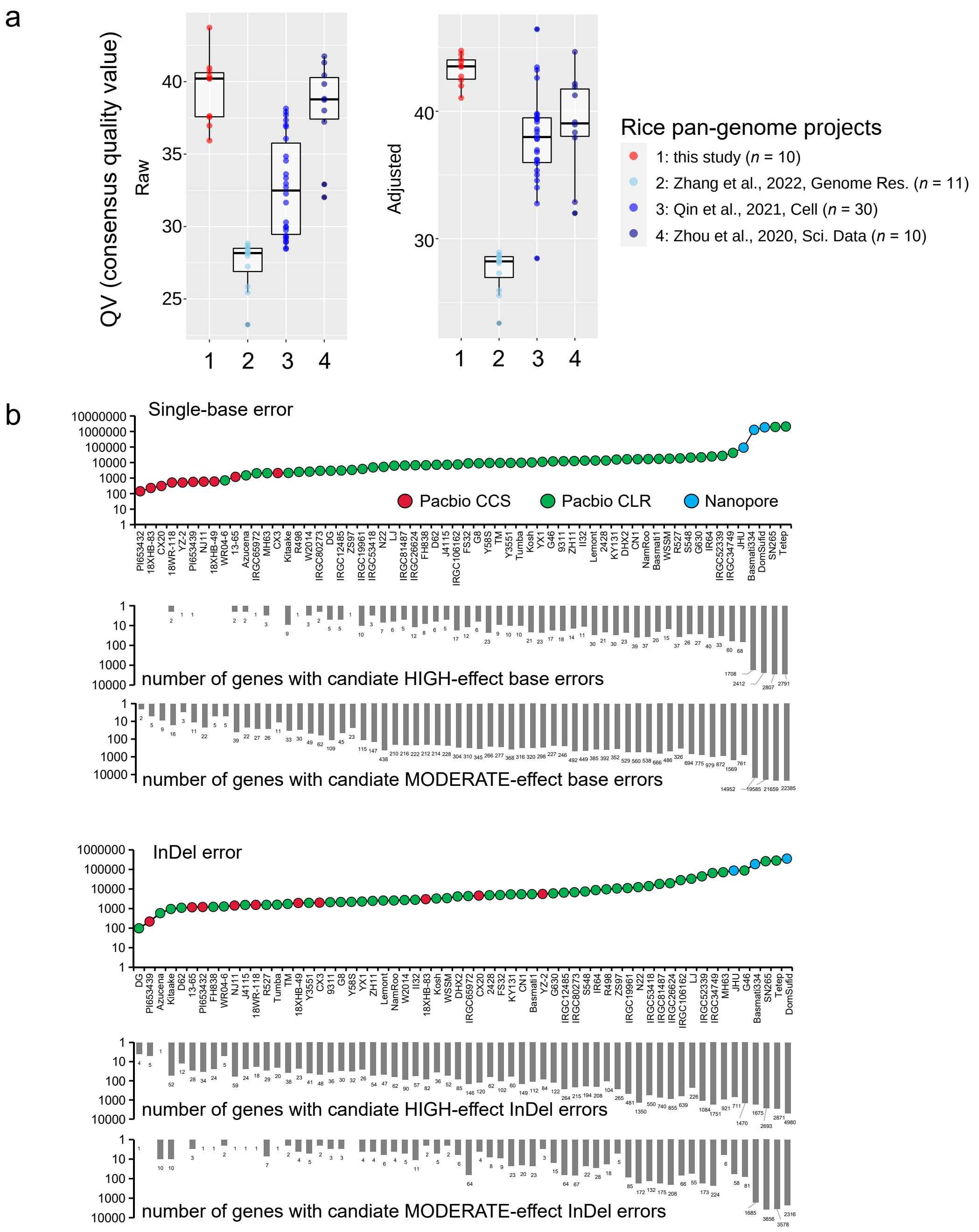


**Supplementary Fig. 4** Assembly quality assessment in base accuracy. (**a**) QVs for rice genomes in four rice pan-genome projects, using yak (https://github.com/lh3/yak). In the boxplots, the horizontal line shows the median value, and the whiskers show the 25% and 75% quartile values of QVs. (**b**) The number and annotation to SNVs and InDels for each genome by mapping NGS short reads against itself assembly.

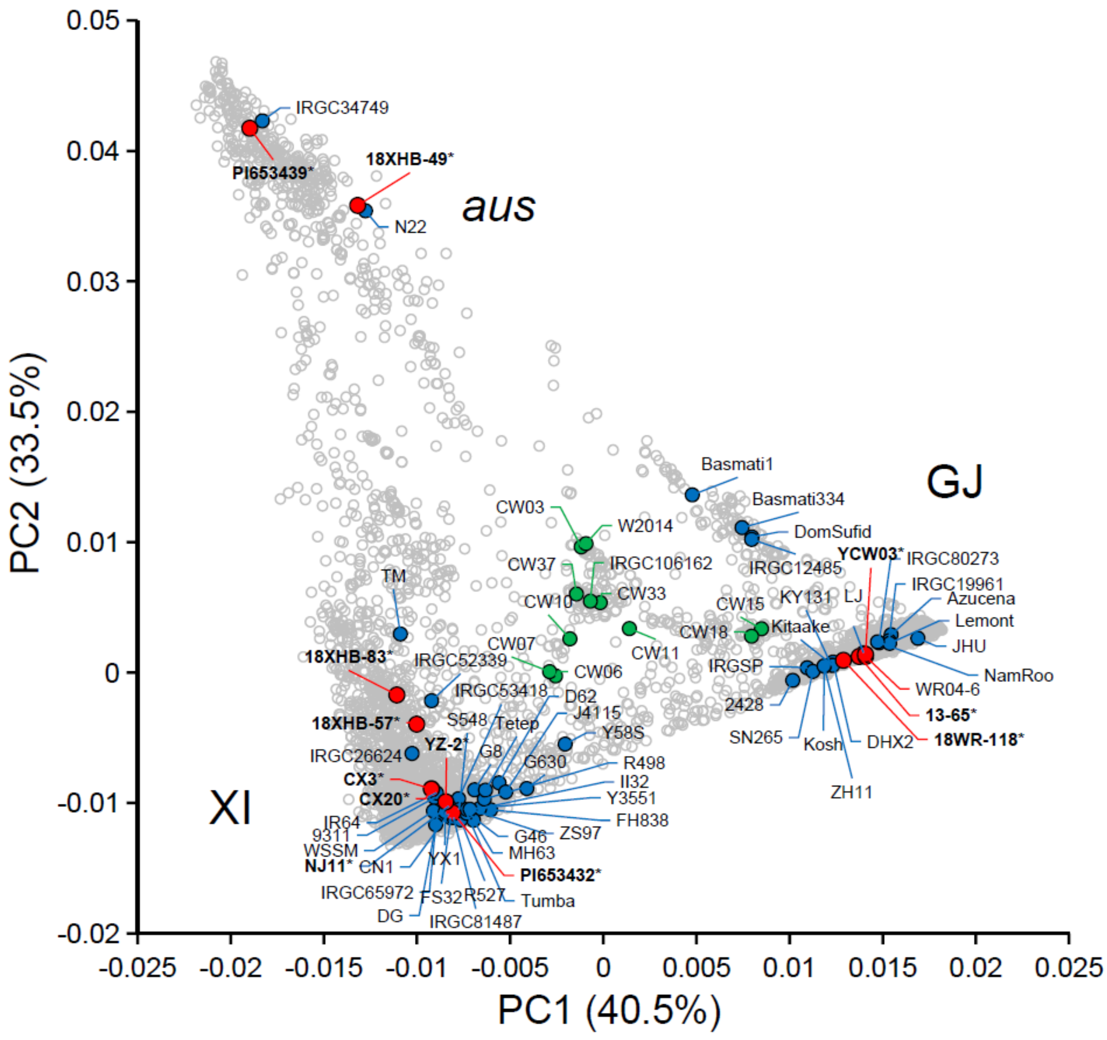


**Supplementary Fig. 5** PCA analysis reveals the representativeness and diversity of genome assemblies used in this study. The first two principle components are shown. Filled circles indicate the assemblies used in this study. Green, blue and red represent wild, cultivated and weedy assemblies, respectively.

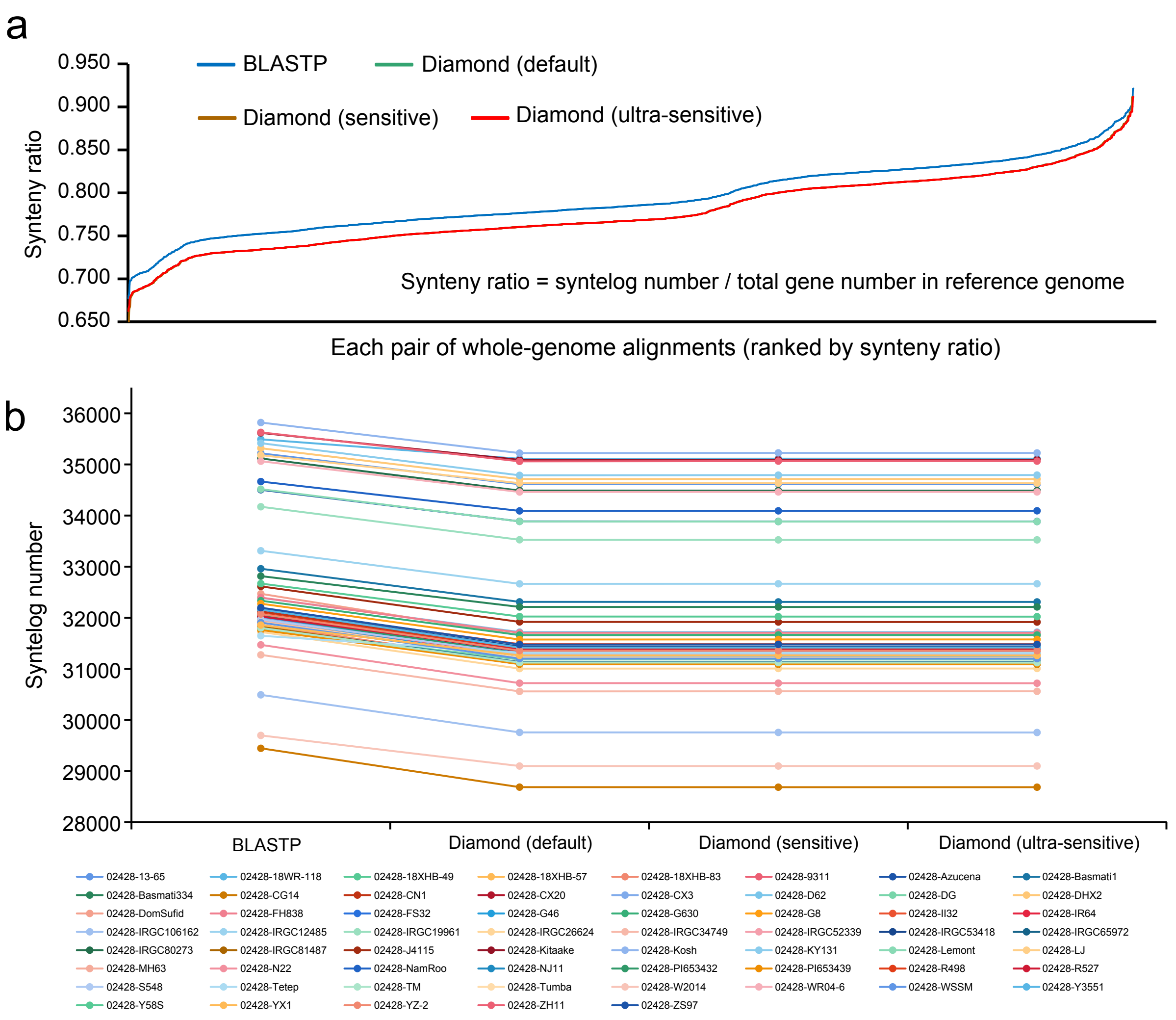


**Supplementary Fig. 6** Performance of BLASTP and Diamond (under different modes) in syntelog identification. (**a**) BLASTP approach identifies more syntelogs than Diamond. (**b**) Syntelog number between accession 02428 with other accessions identified using BLASTP and Diamond under different modes.

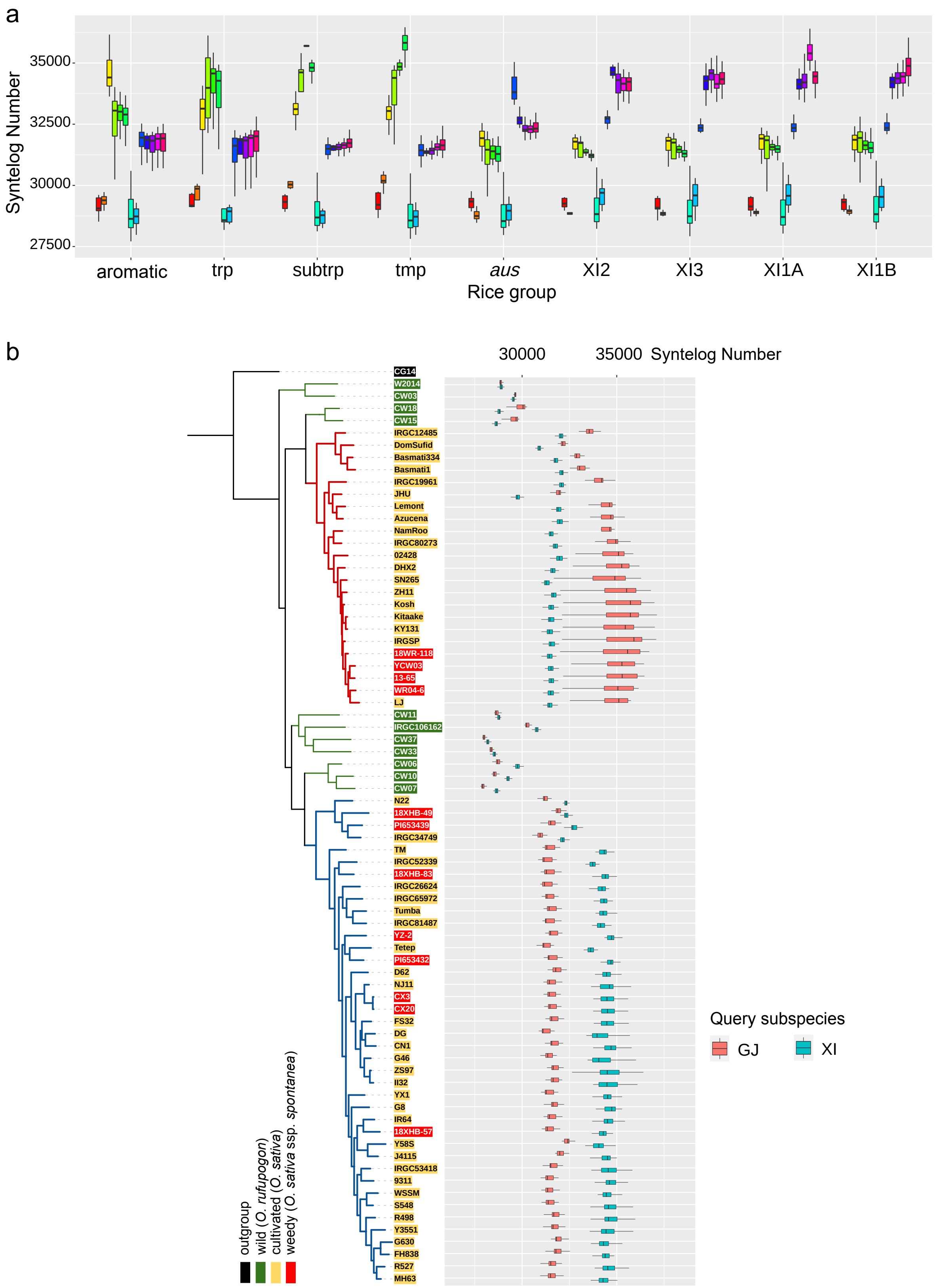


**Supplementary Fig. 7** Pairwise synteny reveals evolutionary signatures in groups and individuals. (**a**) Syntelog numbers between rice groups. (**b**) Syntelog numbers between each accession to other accessions from GJ and XI (including *aus*) subspecies. In the boxplots, the horizontal line shows the median value, and the whiskers show the 25% and 75% quartile values of syntelog numbers.

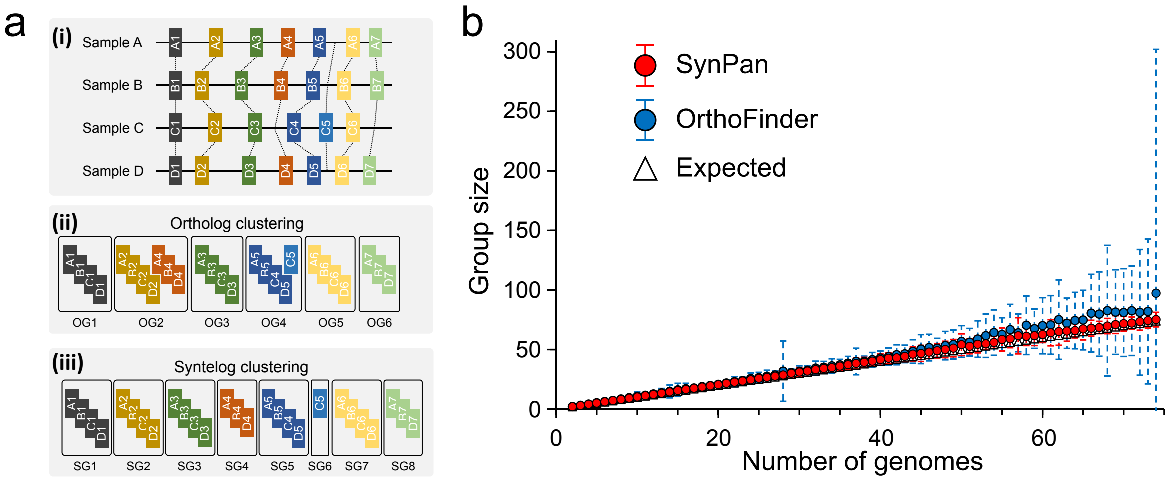


**Supplementary Fig. 8** Comparison between MCL and synteny-based clustering. (**a**) A brief scheme illustrating MCL ortholog clustering and syntelog clustering. (**b**) Group size comparison using synteny-based clustering (SynPan) and MCL clustering (OrthoFinder).

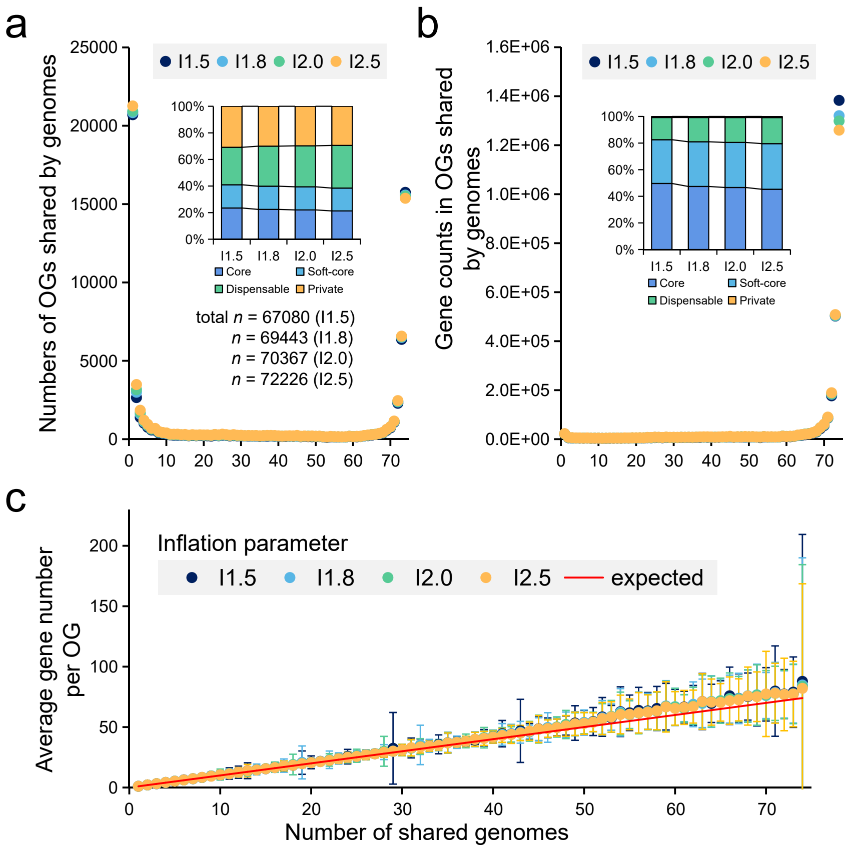


**Supplementary Fig. 9** Benchmarking analysis on the influences of inflation parameters in the MCL clustering in rice genomes. (**a**) The OG numbers shared by different sizes of rice genomes under different inflation parameters. (**b**) Gene counts in OGs shared by different sizes of genomes. (**c**) Average gene number per OG under different inflation parameters.

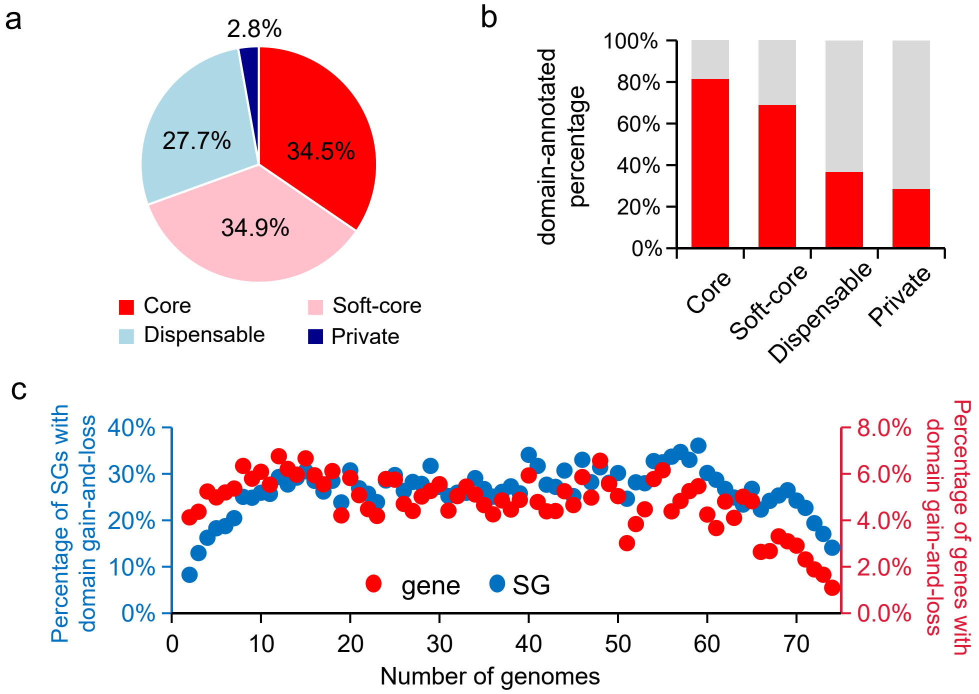


**Supplementary Fig. 10** Composition and features of rice syntelog-based pan-genome. (**a**) Pan-gene composition (core, soft-core, dispensable and private) of rice pan-genome. (**b**) Proportions of domain-annotated genes in four categories of rice pan-genome. (**c**) Percentage of SGs (blue) and genes (red) with domain gain-and-loss.

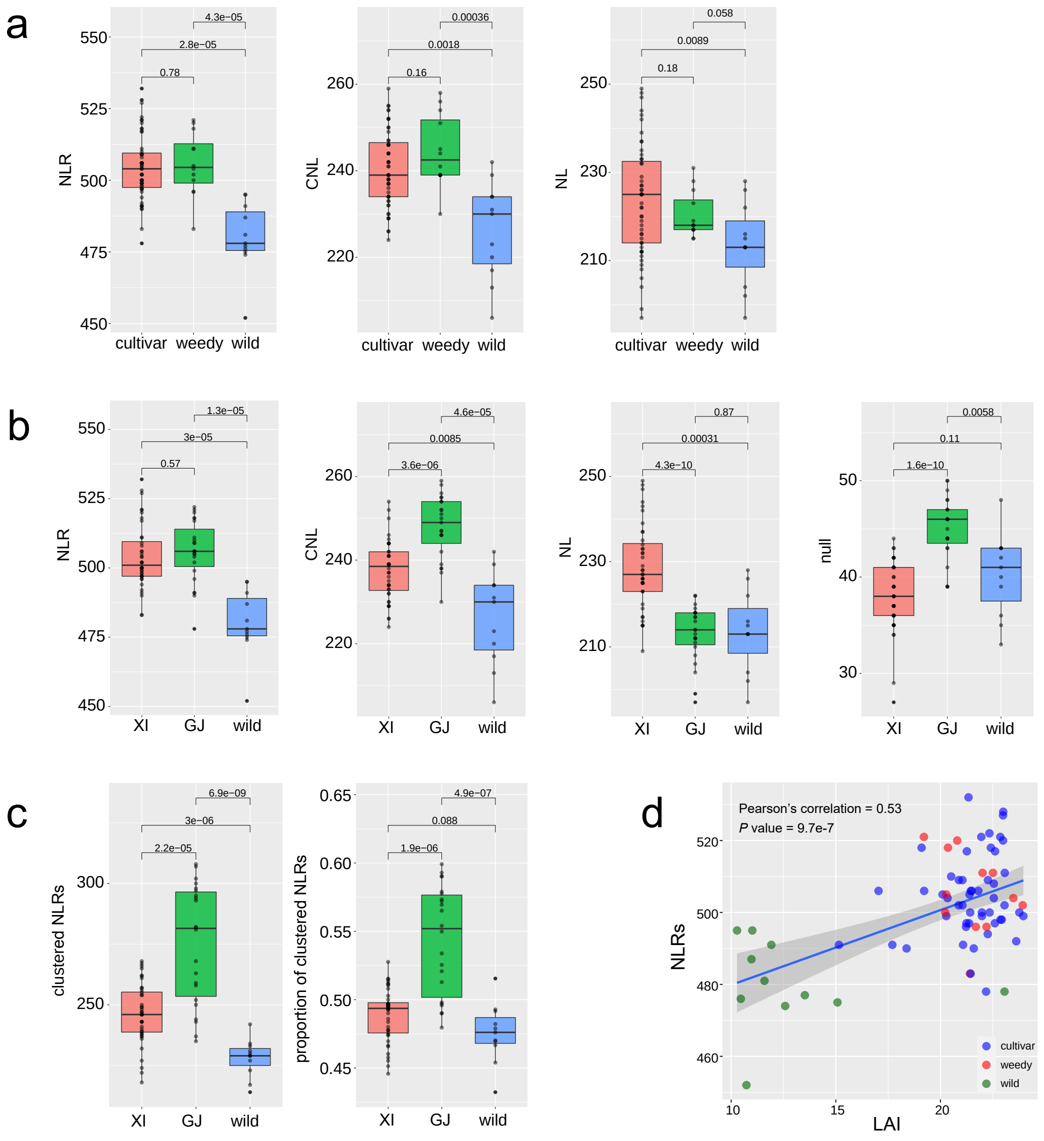


**Supplementary Fig. 11** Comparison of NLR genes in rice genomes from different ecotypes and subspecies. (**a**) Distribution of different NLR genes in wild, cultivated and weedy accessions. (**b**) Distribution of different NLR genes in wild rice, XI and GJ. (**c**) Clustered NLRs in wild rice, XI and GJ. *P* values are calculated using Wilcoxon test. (**d**) Relationship between genome assembly completeness (as indicated by LAI) and NLR size. In the boxplots, the horizontal line shows the median value, and the whiskers show the 25% and 75% quartile values of NLR sizes.

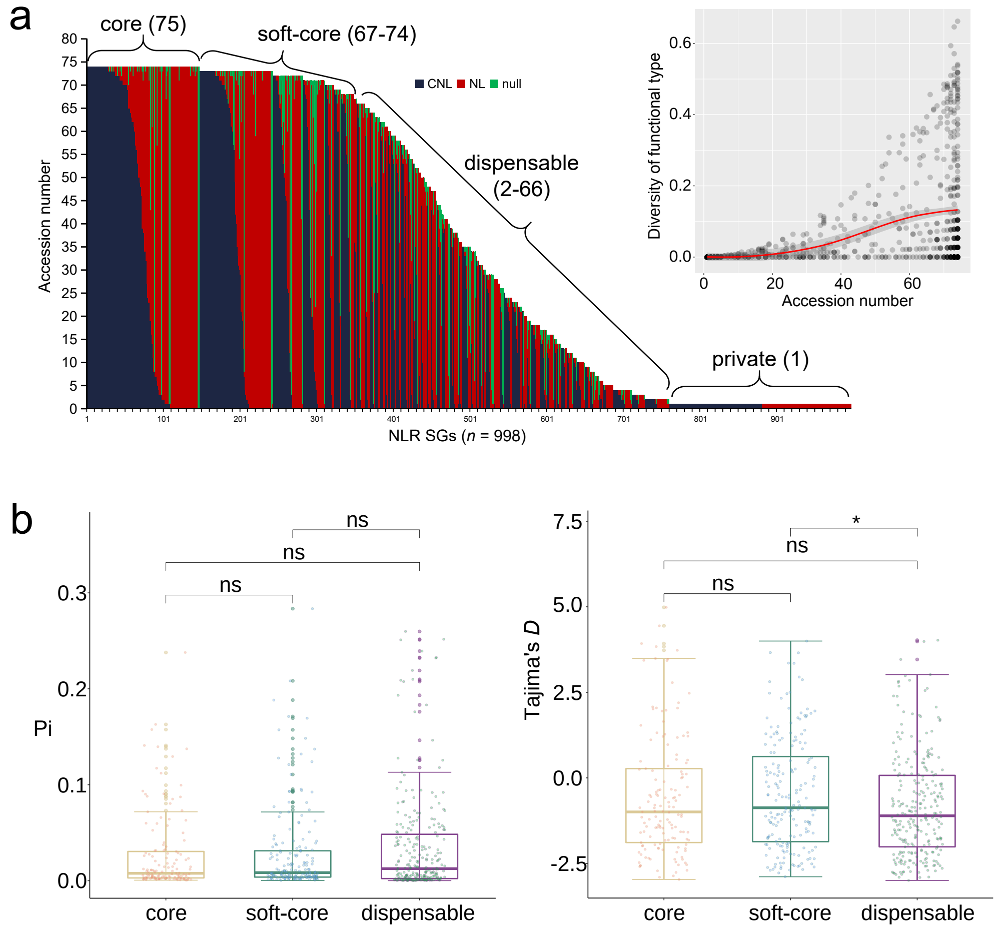


**Supplementary Fig. 12** Genomic features of NLRs in rice genomes. (**a**) Dynamics of domain architectures in the rice NLRome. CNL, NL and null (no canonical architectures) are defined as three NLR architecture types in rice NLRome. Multiple types are observed for most SGs. (**b**) Nucleotide diversity and selection of NLRs from core, soft-core and dispensable SGs. In the boxplots, the horizontal line shows the median value, and the whiskers show the 25% and 75% quartile values of Pi and Tajima's *D*. Significance *P* values are performed using Wilcoxon test. *, $P < 0.05$.

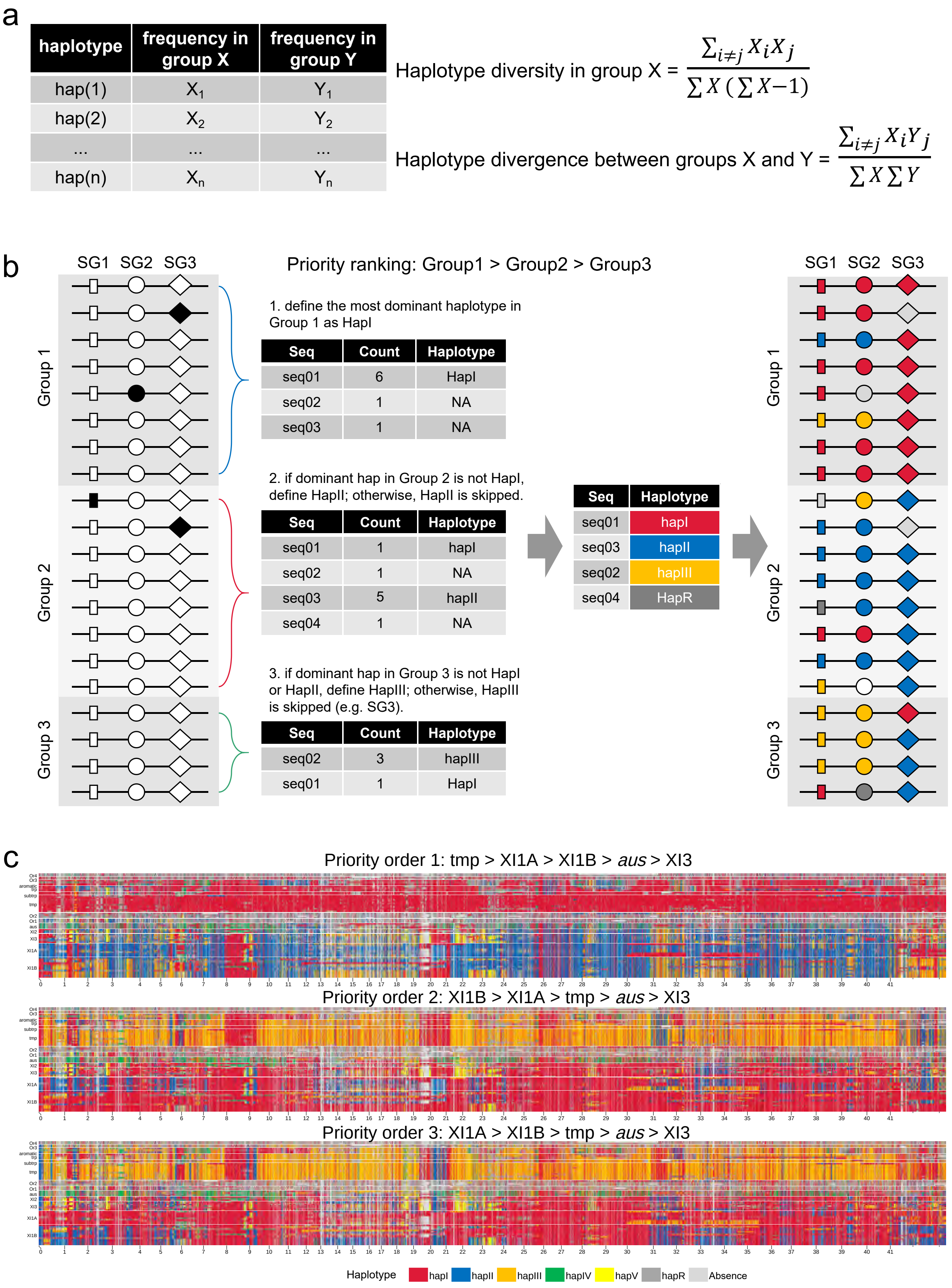
a
haplotype
frequency in group X
frequency in group Y
hap(1)
$X_1$
$Y_1$
hap(2)
$X_2$
$Y_2$
...
hap(n)
$X_n$
$Y_n$
Haplotype diversity in group X = $\frac{\sum_{i \neq j} X_i X_j}{\sum X (\sum X - 1)}$
Haplotype divergence between groups X and Y = $\frac{\sum_{i \neq j} X_i Y_j}{\sum X \sum Y}$
b
SG1 SG2 SG3
Group 1
Group 2
Group 3
Priority ranking: Group1 > Group2 > Group3
1. define the most dominant haplotype in Group 1 as HapI
Seq | Count | Haplotype
seq01 | 6 | HapI
seq02 | 1 | NA
seq03 | 1 | NA
2. if dominant hap in Group 2 is not HapI, define HapII; otherwise, HapII is skipped.
Seq | Count | Haplotype
seq01 | 1 | hapI
seq02 | 1 | NA
seq03 | 5 | hapII
seq04 | 1 | NA
3. if dominant hap in Group 3 is not HapI or HapII, define HapIII; otherwise, HapIII is skipped (e.g. SG3).
Seq | Count | Haplotype
seq02 | 3 | hapIII
seq01 | 1 | HapI
Seq | Haplotype
seq01 | hapI
seq03 | hapII
seq02 | hapIII
seq04 | HapR
c
Priority order 1: tmp > XI1A > XI1B > aus > XI3
Priority order 2: XI1B > XI1A > tmp > aus > XI3
Priority order 3: XI1A > XI1B > tmp > aus > XI3
Haplotype
hapI
hapII
hapIII
hapIV
hapV
hapR
Absence

**Supplementary Fig. 13** Haplotype analysis on rice syntelogs. (**a**) Definition of haplotype diversity and divergence. Haplotype diversity and divergence represent average haplotype differences among sequences in a syntelog group within one group and among two groups, respectively, where Xi and Xj are the presence count of haplotype i and haplotype j in group X, and ∑X is the total sequence count within a syntelog group. (**b**) Brief scheme illustrating the assignment and visualization of ancestral haplotypes for each genome. A group priority-based referring strategy is used to assign ancestral haplotypes. Group information is prior based on whole-genome phylogeny or population structure. The most dominant sequence in syntelogs from Group 1 is set as hapI. If the most dominant sequence from Group 2 is not HapI, then define HapII, otherwise HapII is skipped. If the most dominant sequence from Group 3 is neither HapI nor HapII, then define HapIII, otherwise HapIII is skipped. By analogy, dominant haplotypes are determined and colored for each syntelog group. Rare haplotypes are named as HapR colored by dark gray. In this study, the group priority order is set as tmp > XI1A > XI1B > *aus* > XI3. Different orders have no influences on the calculation of haplotype diversity and divergence. (**c**) Mosaic graphs of ancestral haplotypes on chromosome 1 across 74 rice genomes with different priority orders: tmp > XI1A > XI1B > *aus* > XI3, XI1B > XI1A > tmp > *aus* > XI3, and XI1A > XI1B > tmp > *aus* > XI3. For each window, the same color indicates the same haplotype, and dark and light gray indicate rare haplotypes and syntelog absence, respectively.

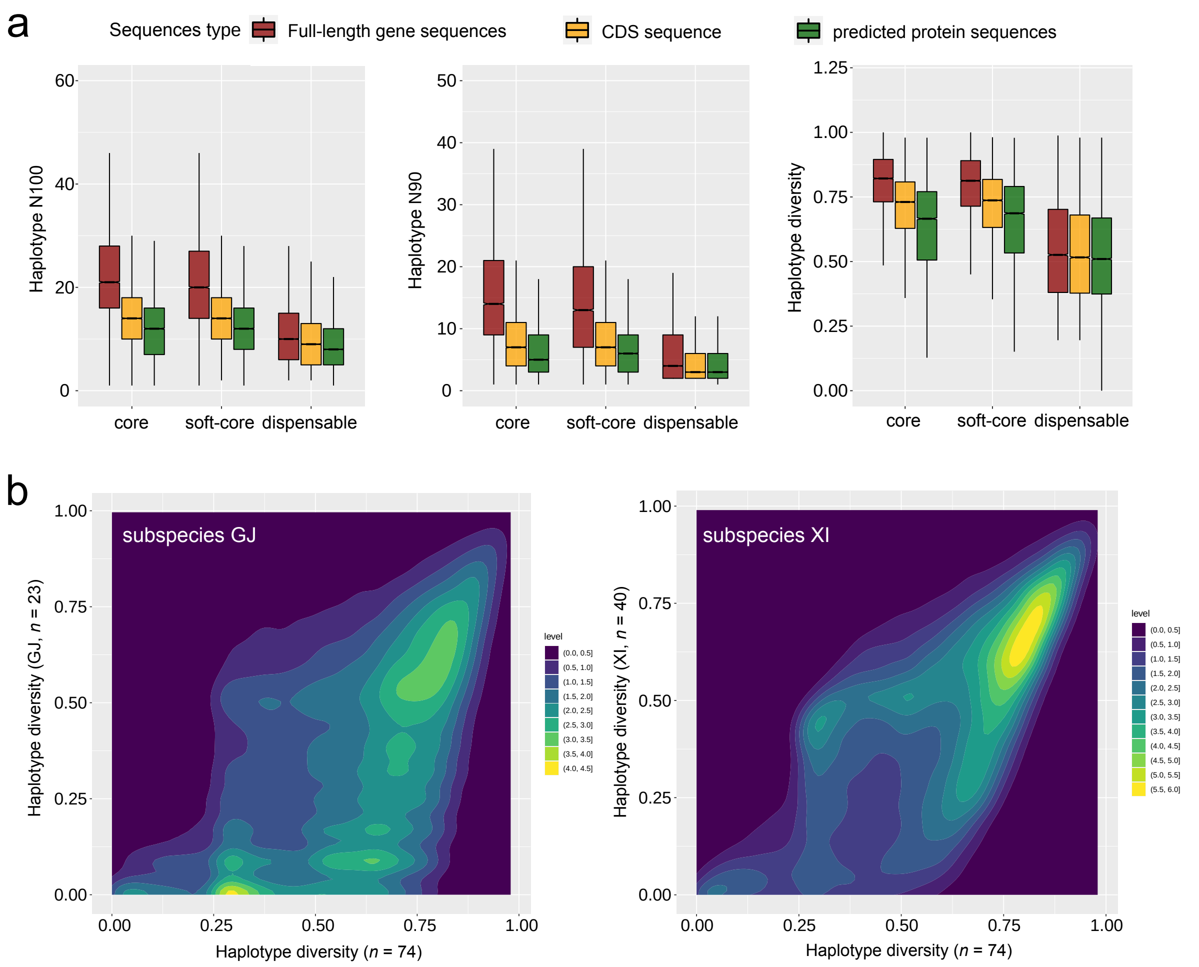


**Supplementary Fig. 14** Genetic diversity measured by full-length gene, coding region, and predicted protein sequences in rice genomes. (a) Comparison of diversity measured by haplotype N100, N90 and haplotype diversity using full-length nucleotide sequences, coding sequences and predicted protein sequences. In the boxplots, the horizontal line shows the median value, and the whiskers show the 25% and 75% quartile values of each diversity indice. (b) Haplotype diversity in GJ and XI, compared to all rice genomes.

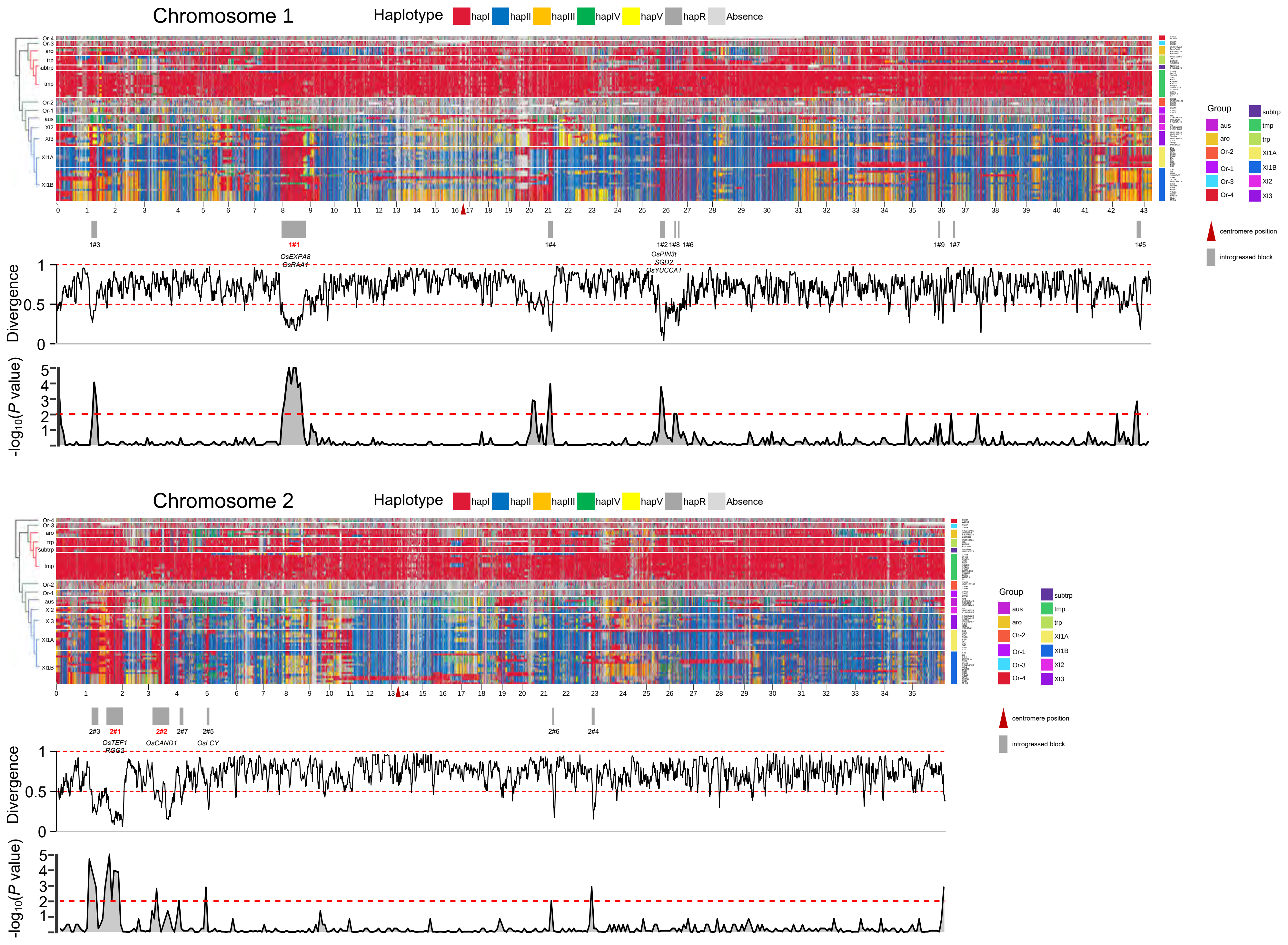

Chromosome 1
Haplotype
hapI
hapII
hapIII
hapIV
hapV
hapR
Absence
Or-4
Or-3
aro
trp
subtrp
tmp
Or-2
Or-1
aus
XI2
XI3
XI1A
XI1B
Group
centromere position
introgressed block
1#3
1#1
OsEXPA8
OsRAA1
1#4
1#2 1#8 1#6
OsPIN3t
SGD2
OsYUCCA1
1#9 1#7
1#5
Divergence
-log10(P value)
Chromosome 2
2#3
2#1
OsTEF1
RGG2
2#2
OsCAND1
2#7
2#5
OsLCY
2#6
2#4

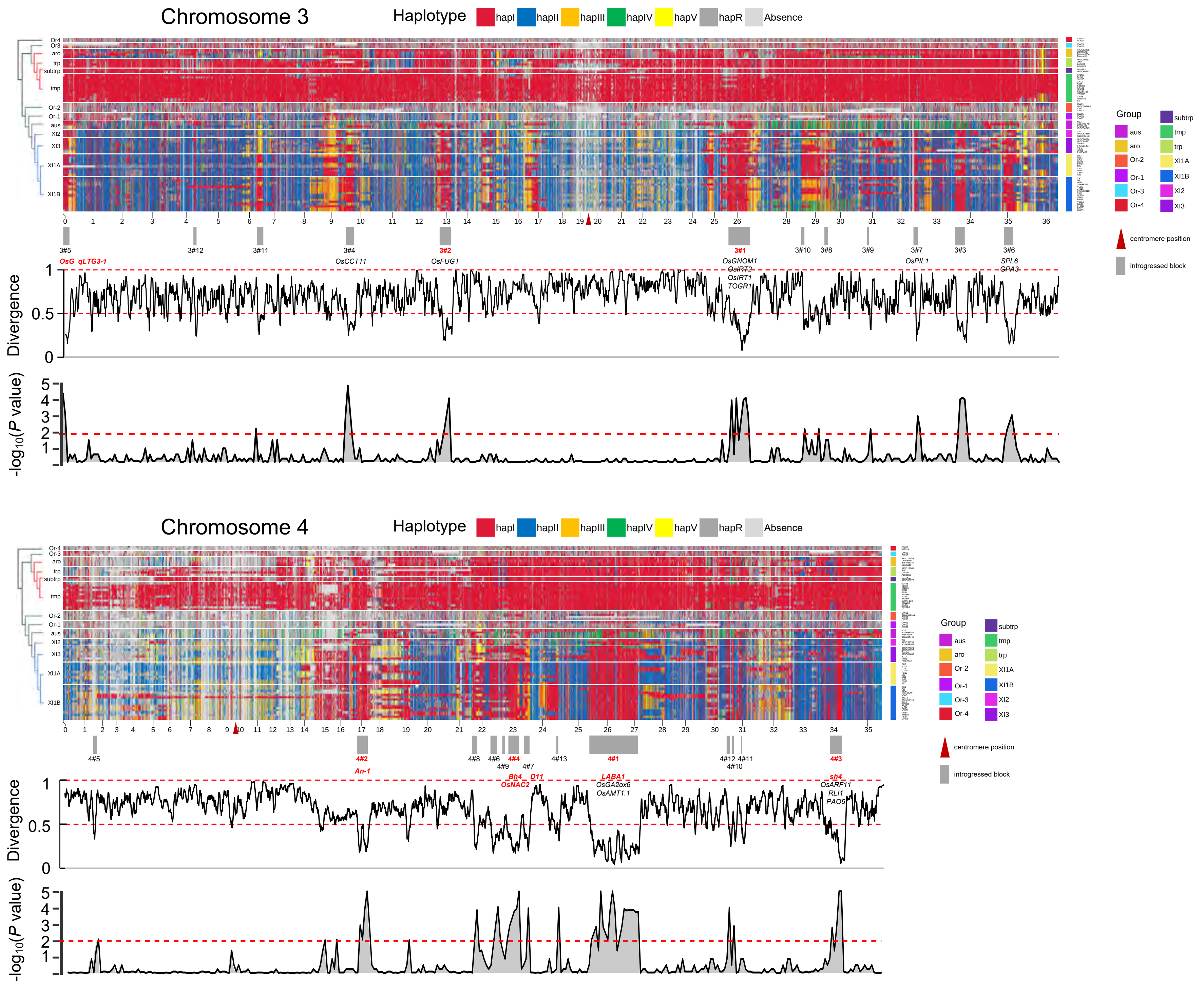

Chromosome 3
Haplotype
hapI
hapII
hapIII
hapIV
hapV
hapR
Absence
Or4
Or3
aro
trp
subtrp
tmp
Or-2
Or-1
aus
XI2
XI3
XI1A
XI1B
Group
centromere position
introgressed block
3#5
3#12
3#11
3#4
3#2
3#1
3#10
3#8
3#9
3#7
3#3
3#6
OsG qLTG3-1
OsCCT11
OsFUG1
OsGNOM1
OsIRT2
OsIRT1
TOGR1
OsPIL1
SPL6
GPA3
Divergence
-log10(P value)
Chromosome 4
Or-4
Or-3
4#5
4#2
4#8
4#6
4#9
4#4
4#7
4#13
4#1
4#12
4#10
4#11
4#3
An-1
Bh4
OsNAC2
D11
LABA1
OsGA2ox6
OsAMT1.1
sh4
OsARF11
RLI1
PAO5

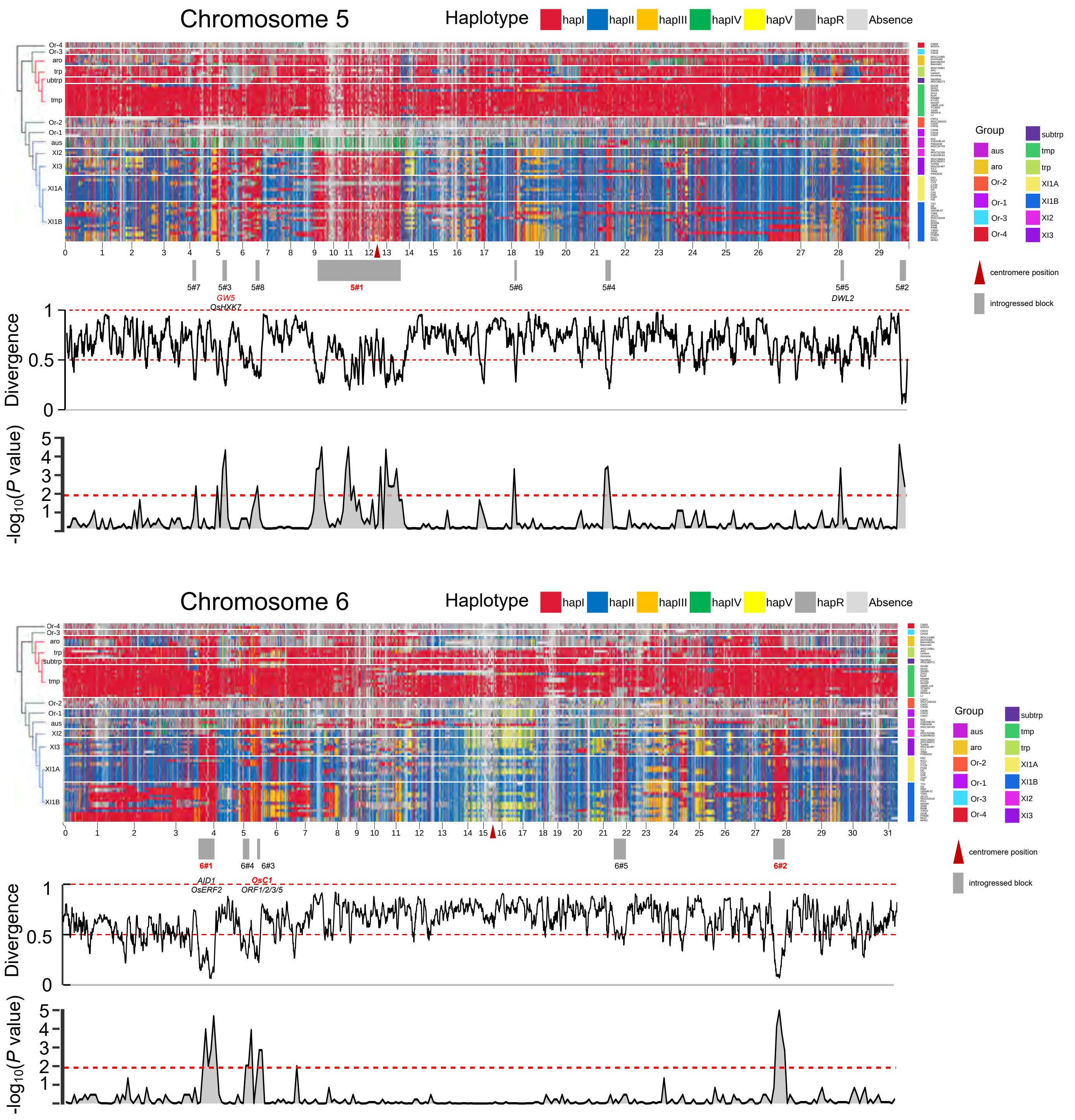

Chromosome 5
Haplotype
hapI
hapII
hapIII
hapIV
hapV
hapR
Absence
Or-4
Or-3
aro
trp
subtrp
tmp
Or-2
Or-1
aus
XI2
XI3
XI1A
XI1B
Group
aus
aro
Or-2
Or-1
Or-3
Or-4
subtrp
tmp
trp
XI1A
XI1B
XI2
XI3
centromere position
introgressed block
5#7
5#3
5#8
5#1
5#6
5#4
5#5
5#2
GW5
OsHXK7
DWL2
Divergence
-log10(P value)
Chromosome 6
Haplotype
hapI
hapII
hapIII
hapIV
hapV
hapR
Absence
6#1
6#4
6#3
6#5
6#2
AID1
OsERF2
OsC1
ORF1/2/3/5
Divergence
-log10(P value)

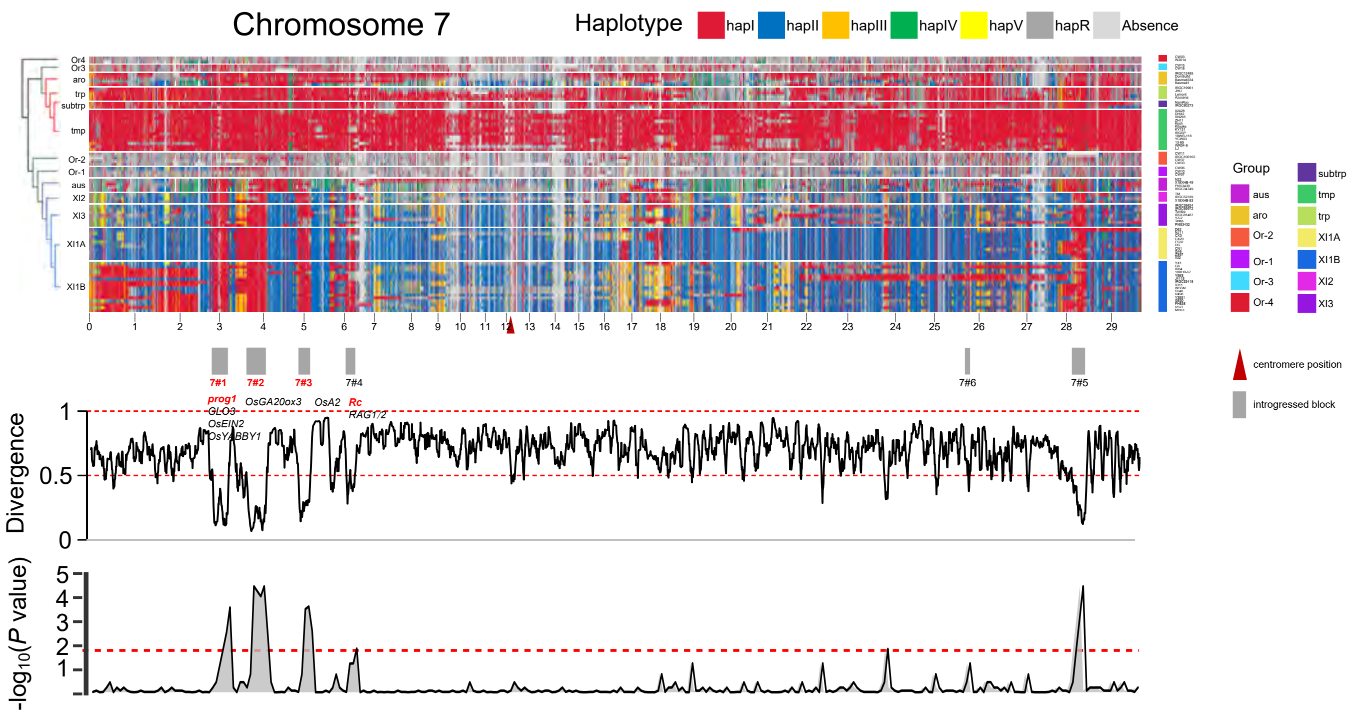

Chromosome 7
Haplotype
hapI
hapII
hapIII
hapIV
hapV
hapR
Absence
Or4
Or3
aro
trp
subtrp
tmp
Or-2
Or-1
aus
XI2
XI3
XI1A
XI1B
7#1
7#2
7#3
7#4
7#6
7#5
prog1
GLO3
OsEIN2
OsYABBY1
OsGA20ox3
OsA2
Rc
RAG1/2
Divergence
-log10(P value)
Group
aus
aro
Or-2
Or-1
Or-3
Or-4
subtrp
tmp
trp
XI1A
XI1B
XI2
XI3
centromere position
introgressed block


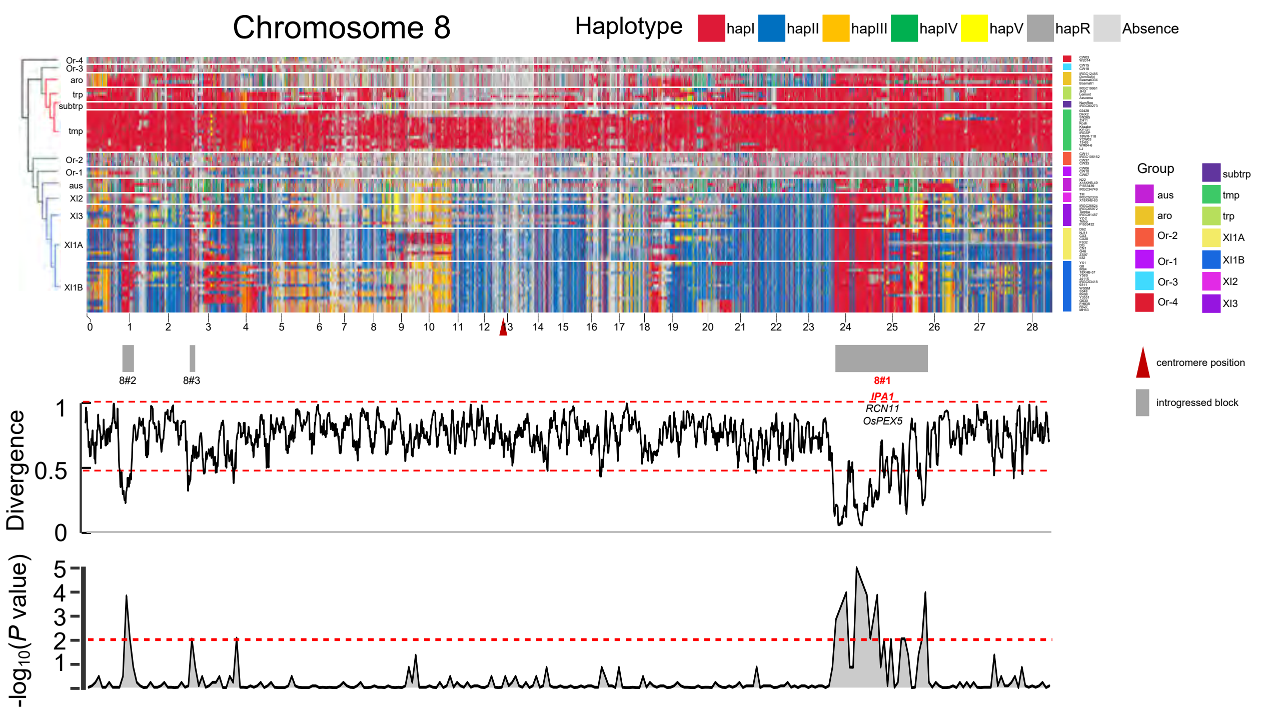

Chromosome 8
Haplotype
hapI
hapII
hapIII
hapIV
hapV
hapR
Absence
Or-4
Or-3
aro
trp
subtrp
tmp
Or-2
Or-1
aus
XI2
XI3
XI1A
XI1B
8#2
8#3
8#1
IPA1
RCN11
OsPEX5
Divergence
-log10(P value)
Group
aus
aro
Or-2
Or-1
Or-3
Or-4
subtrp
tmp
trp
XI1A
XI1B
XI2
XI3
centromere position
introgressed block

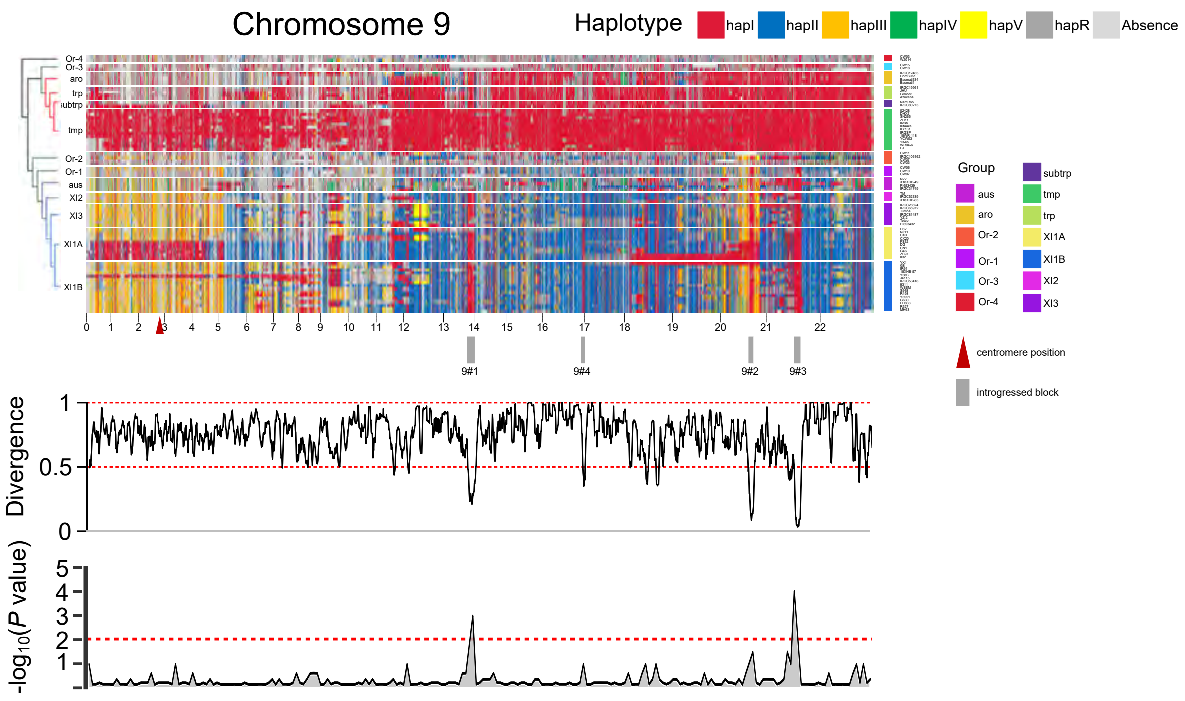
Chromosome 9
Haplotype
hapI
hapII
hapIII
hapIV
hapV
hapR
Absence
Or-4
Or-3
aro
trp
subtrp
tmp
Or-2
Or-1
aus
XI2
XI3
XI1A
XI1B
0 1 2 3 4 5 6 7 8 9 10 11 12 13 14 15 16 17 18 19 20 21 22
9#1
9#4
9#2
9#3
Group
aus
aro
Or-2
Or-1
Or-3
Or-4
subtrp
tmp
trp
XI1A
XI1B
XI2
XI3
centromere position
introgressed block
Divergence
1
0.5
0
-log10(P value)
5
4
3
2
1

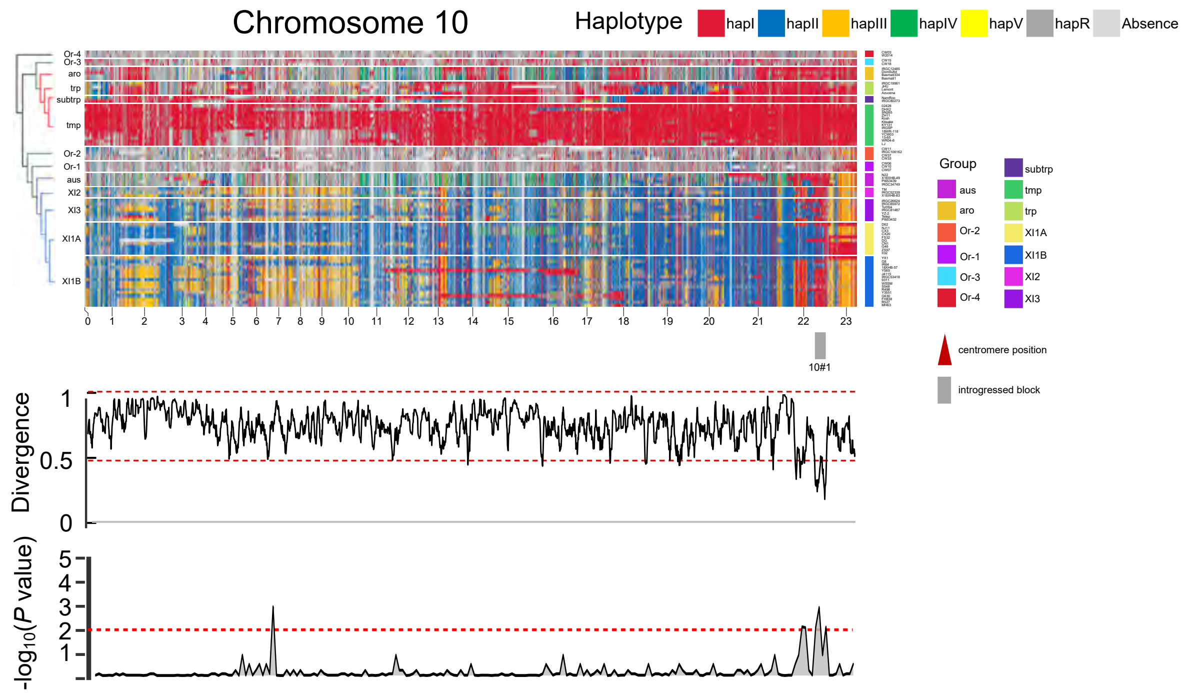
Chromosome 10
Haplotype
hapI
hapII
hapIII
hapIV
hapV
hapR
Absence
Or-4
Or-3
aro
trp
subtrp
tmp
Or-2
Or-1
aus
XI2
XI3
XI1A
XI1B
0 1 2 3 4 5 6 7 8 9 10 11 12 13 14 15 16 17 18 19 20 21 22 23
10#1
Group
aus
aro
Or-2
Or-1
Or-3
Or-4
subtrp
tmp
trp
XI1A
XI1B
XI2
XI3
centromere position
introgressed block
Divergence
1
0.5
0
-log10(P value)
5
4
3
2
1

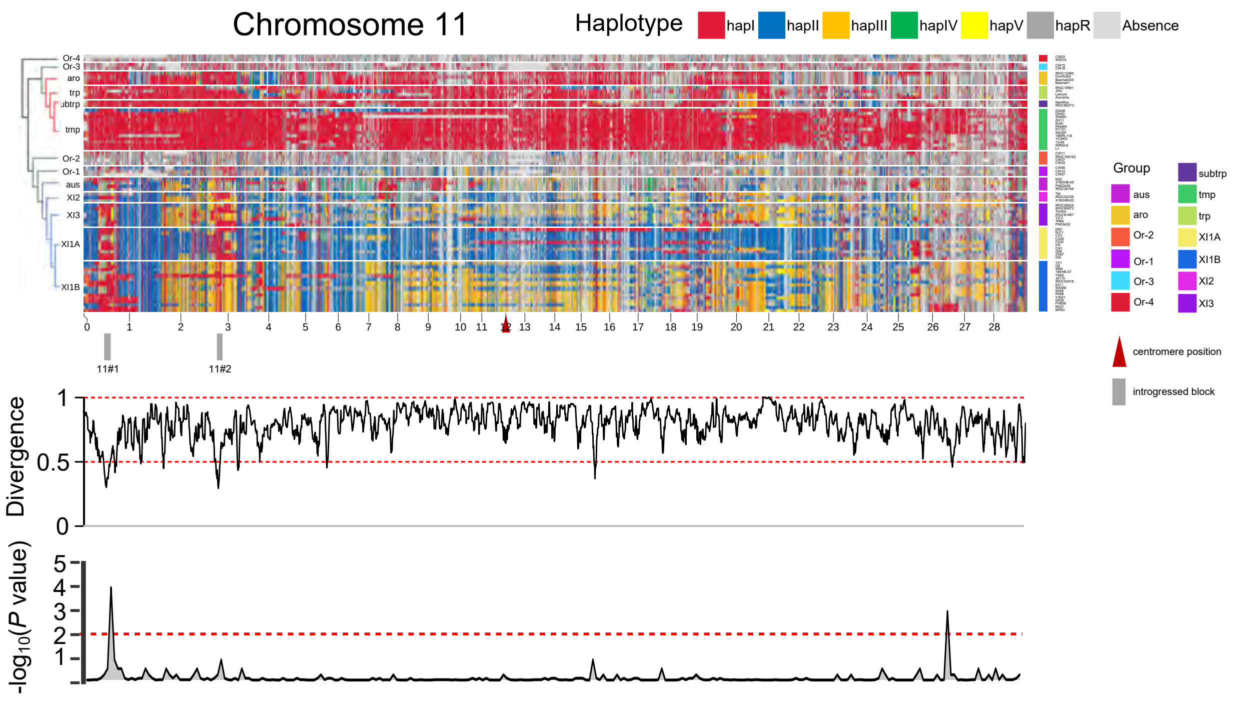
Chromosome 11
Haplotype
hapI
hapII
hapIII
hapIV
hapV
hapR
Absence
Or-4
Or-3
aro
trp
subtrp
tmp
Or-2
Or-1
aus
XI2
XI3
XI1A
XI1B
Group
aus
aro
Or-2
Or-1
Or-3
Or-4
subtrp
tmp
trp
XI1A
XI1B
XI2
XI3
centromere position
introgressed block
11#1
11#2
Divergence
-log$_{10}$($P$ value)

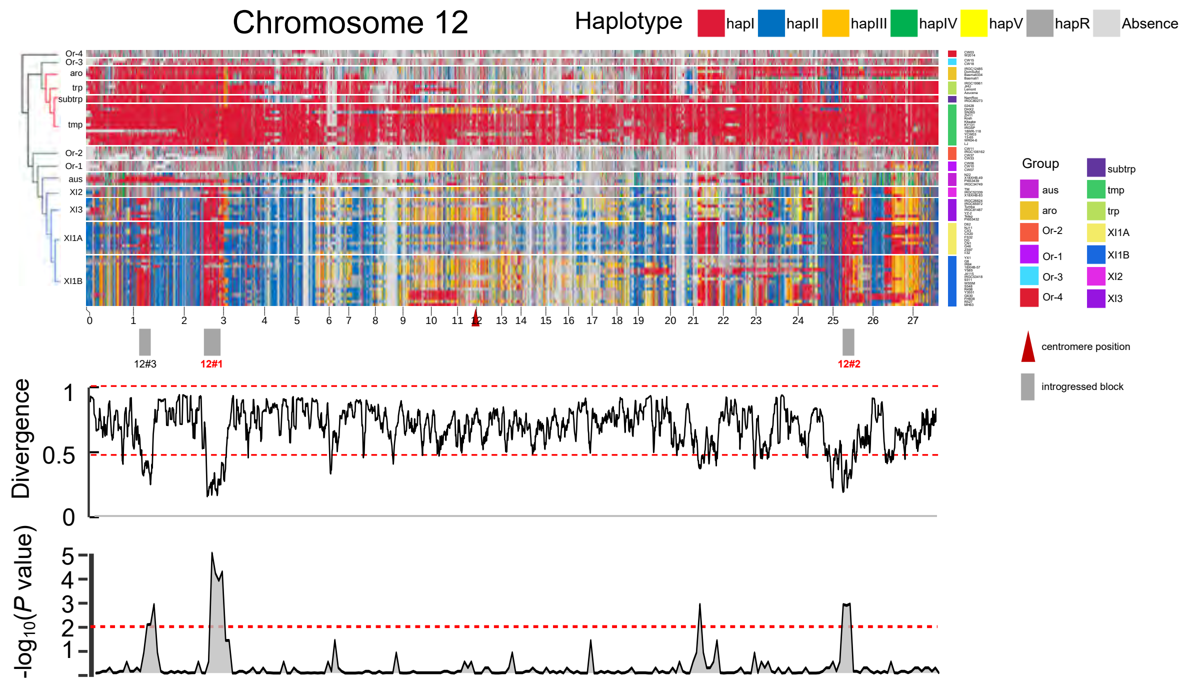
Chromosome 12
Haplotype
hapI
hapII
hapIII
hapIV
hapV
hapR
Absence
Or-4
Or-3
aro
trp
subtrp
tmp
Or-2
Or-1
aus
XI2
XI3
XI1A
XI1B
Group
aus
aro
Or-2
Or-1
Or-3
Or-4
subtrp
tmp
trp
XI1A
XI1B
XI2
XI3
centromere position
introgressed block
12#3
12#1
12#2
Divergence
-log$_{10}$($P$ value)

**Supplementary Fig. 15** Ancestral haplotype landscape on chromosome 1 to chromosome 12. Putative introgression blocks are shown in gray rectangles and numbered. Haplotype divergence and *P* values (scaled by $-\log_{10}$) of non-random distribution significance tests are shown along each chromosome. Red dashed lines represent thresholds to determine introgression blocks.

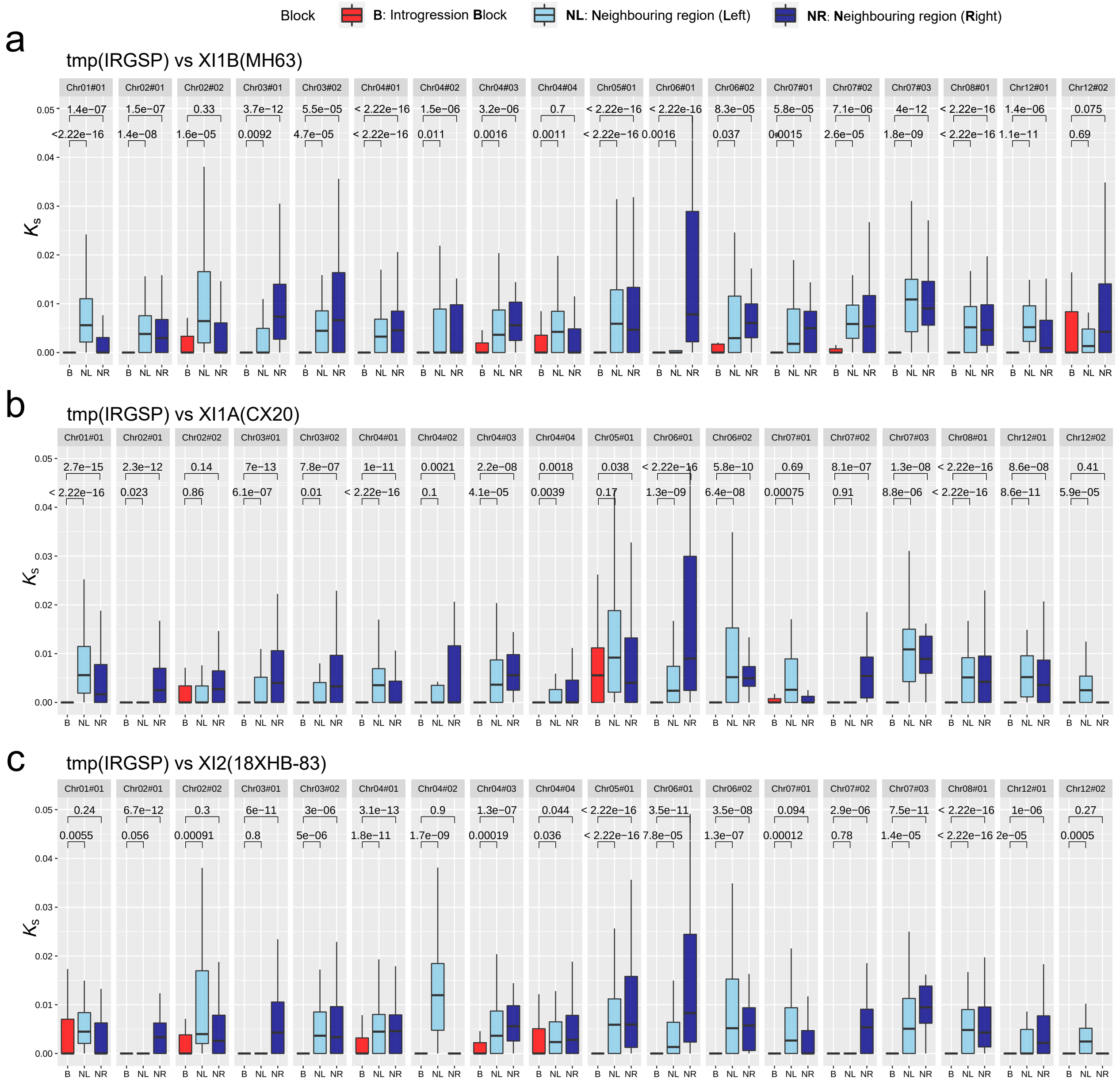


**Supplementary Fig. 16** Synonymous substitution rates ($K_s$) of genes in putative introgression blocks and their neighboring left and right regions. Three replicates (a, b, and c) between the GJ and XI genomes were performed. In the boxplots, the horizontal line shows the median value, and the whiskers show the 25% and 75% quartile values of $K_s$. *P* values are calculated using Wilcoxon test.

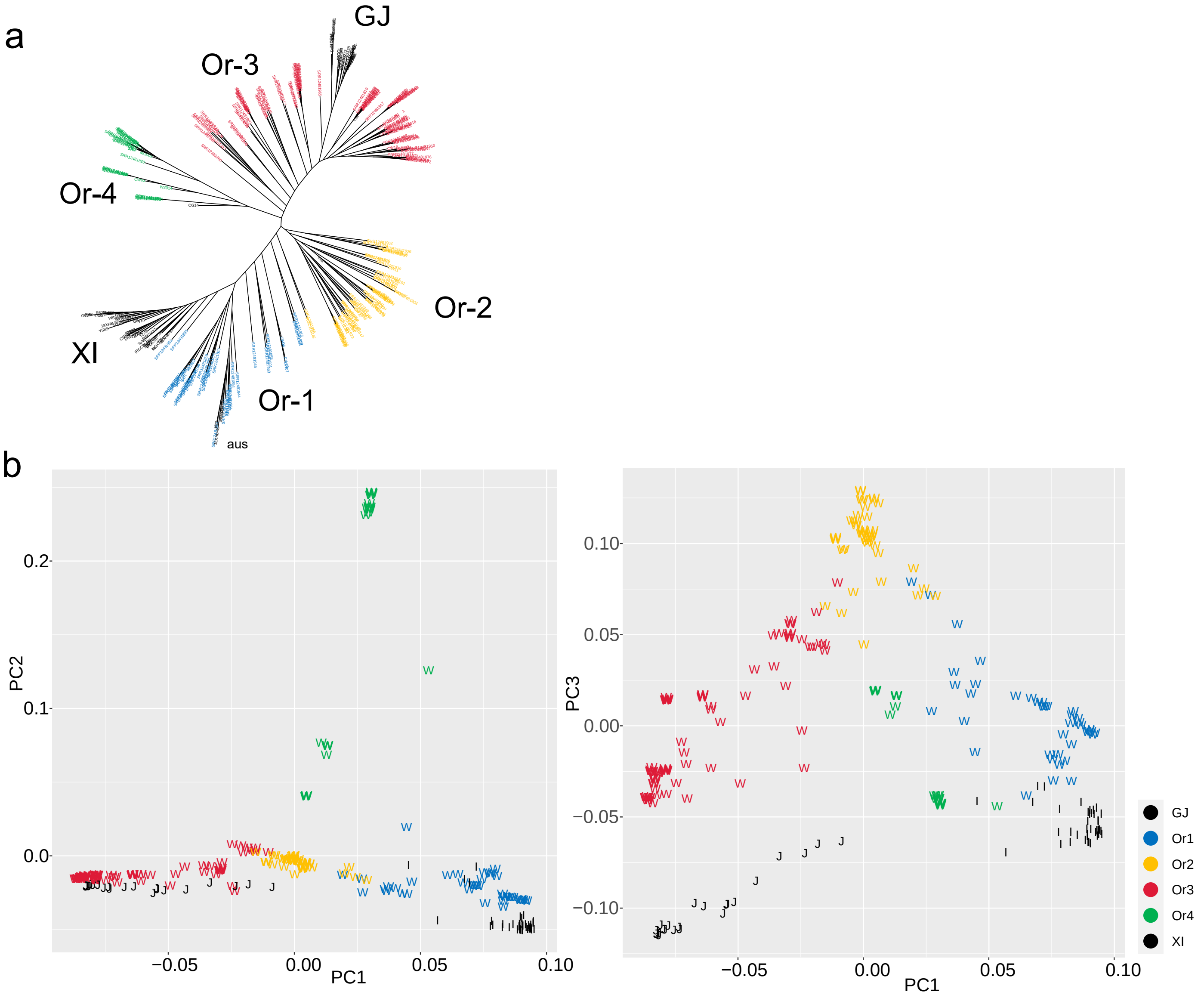


c

| Group | India | Nepal | Sri Lanka | Bangladesh | Myanmar | Laos | Thailand | Cambodia | Vietnam | China | Indonesia | Papua New Guinea | Sum |
|---|---|---|---|---|---|---|---|---|---|---|---|---|---|
| Or-1 | 7 | | 1 | 2 | 1 | 6 | 5 | 6 | 6 | 1 | 2 | | 37 |
| Or-2 | 1 | 1 | 3 | | 4 | 4 | 2 | 5 | 3 | 10 | 4 | 5 | 42 |
| Or-3 | 2 | 6 | 4 | | 1 | | 1 | | 2 | 64 | | | 80 |
| Or-4 | 7 | 8 | 5 | | 5 | | | | | | | | 25 |

**Supplementary Fig. 17** Population structure of wild rice accessions used in this study. (**a**) Phylogenetic tree of *Oryza rufipogon* and *Oryza sativa*. Four wild groups (Or-1, Or-2, Or-3 and Or-4) are indicated in different colors. (**b**) PCA plots of the first three principle components, where “W”, “J” and “I” represent wild, GJ and XI accessions. (**c**) Geographical sources of wild accessions in different groups used in this study

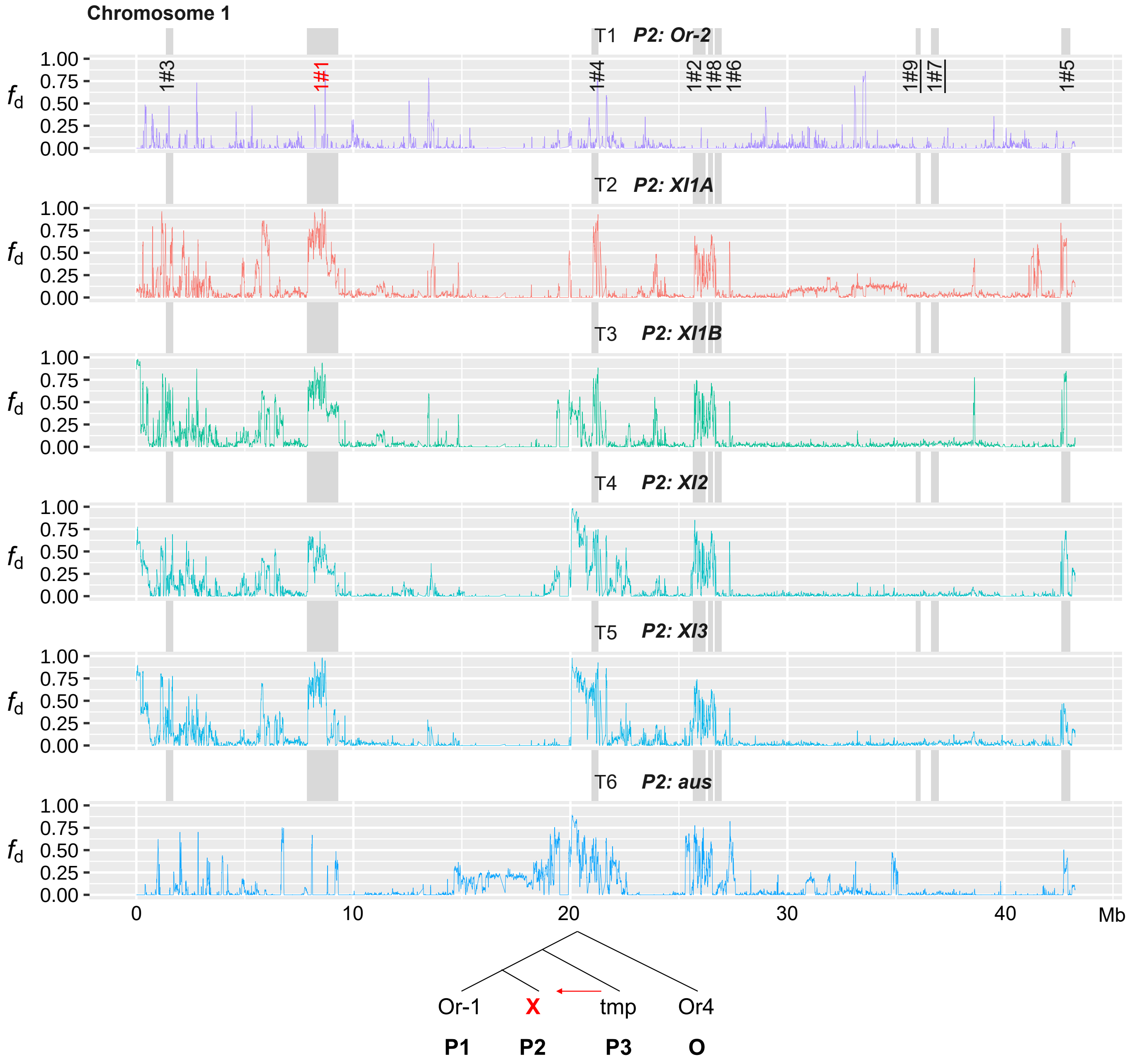


**Supplementary Fig. 18** Introgression fd distributions of ABBA-BABA test on chromosome 1 to chromosome 12 in topology T1 to T6. P1 is Or-1 (*n* = 37), P3 is tmp (GJ, *n* = 100), O/outgroup is Or-4 (*n* = 25), and P2 was set as Or-2 (*n* = 42), XI1A (*n* = 100), XI1B (*n* = 100), XI2 (*n* = 80), XI3 (*n* = 100) and aus (n=60), respectively. T1 is set as a background control in introgression detection. Genomic positions of putative introgression regions are indicated by gray rectangles and detailed coordinates are provided in Supplementary Table 2. Blocks larger than 300kb are highlighted in red and blocks not supported by $f_d$ are underlined.

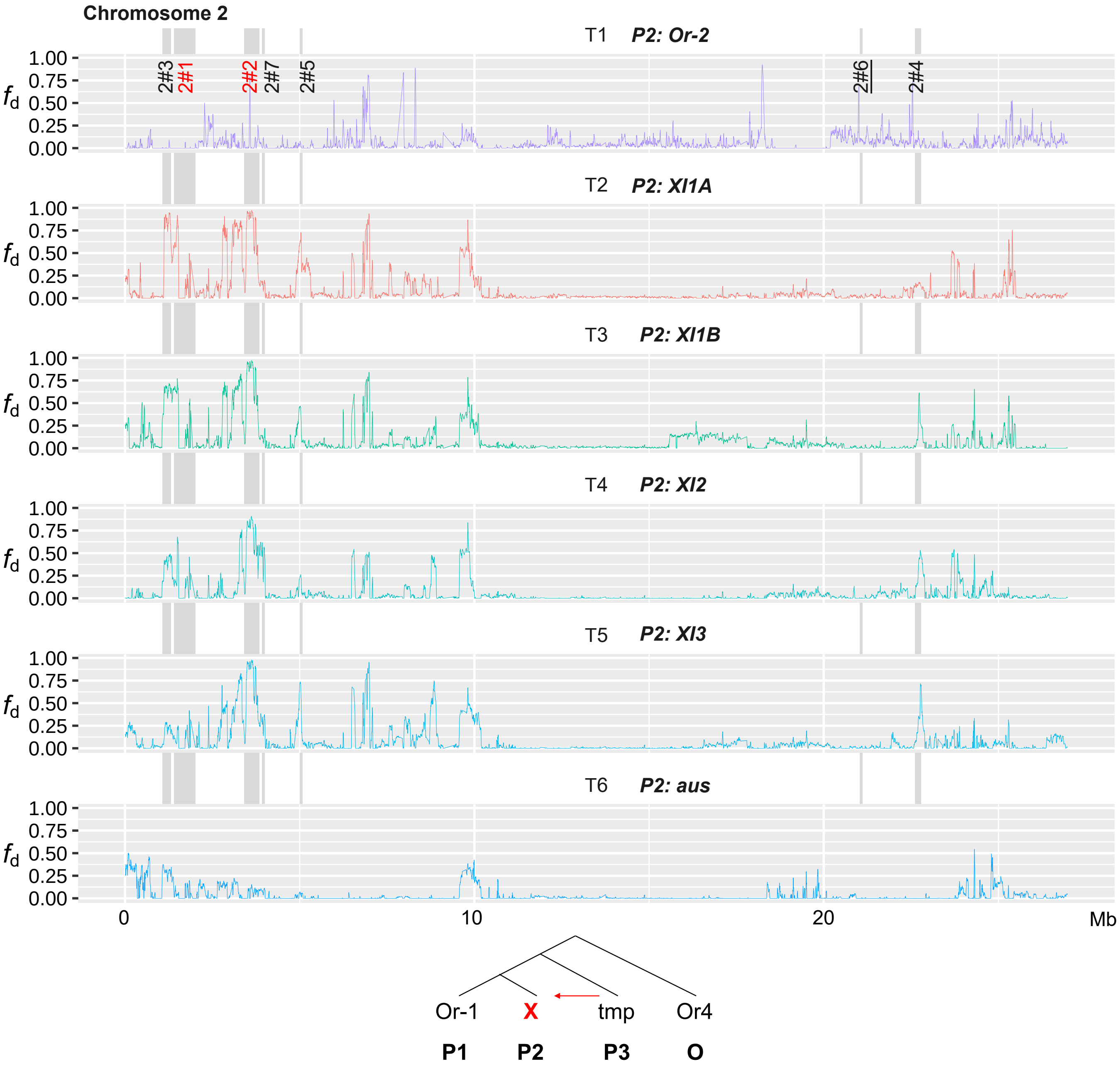


**Supplementary Fig. 18**

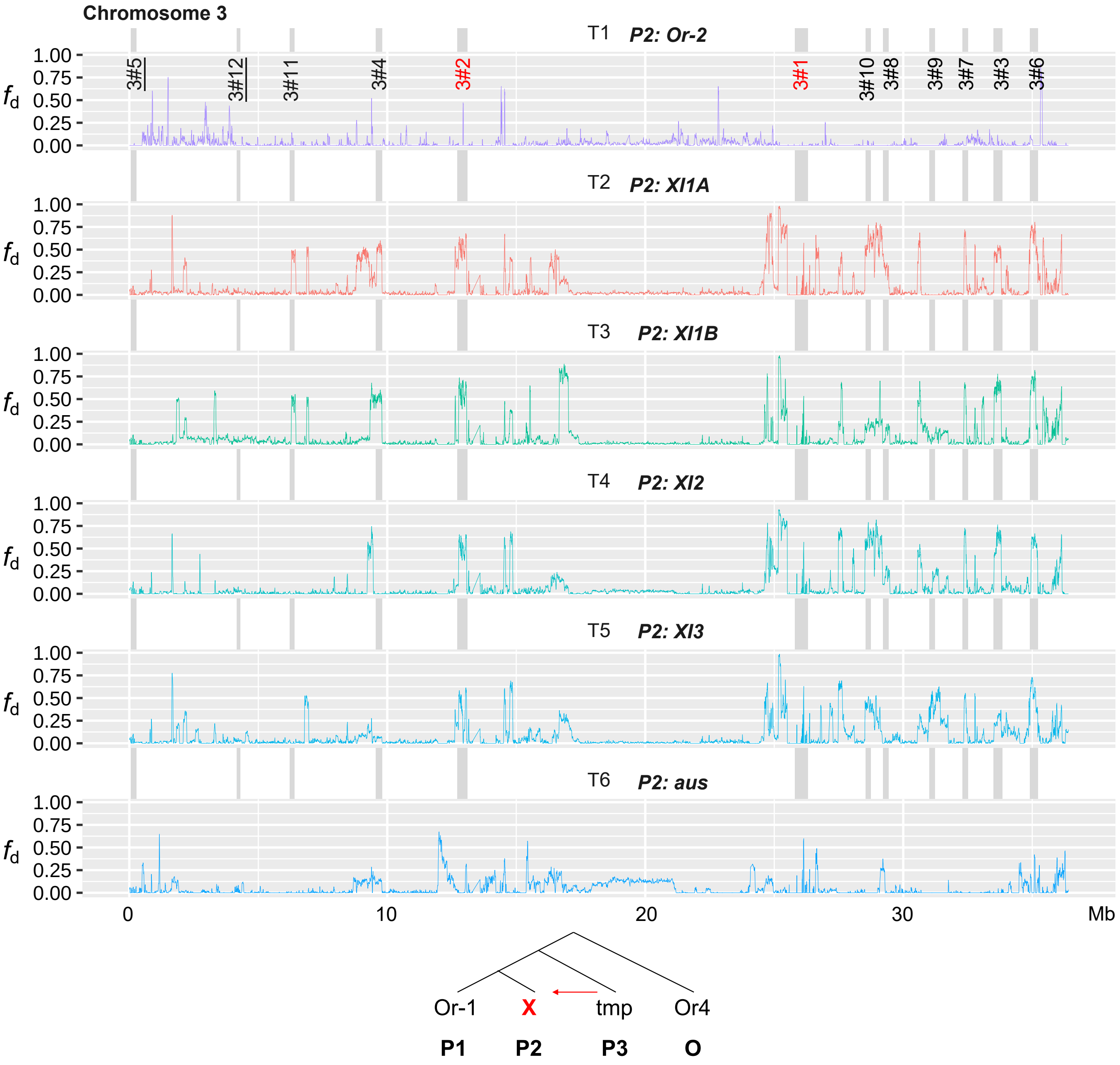


**Supplementary Fig. 18**

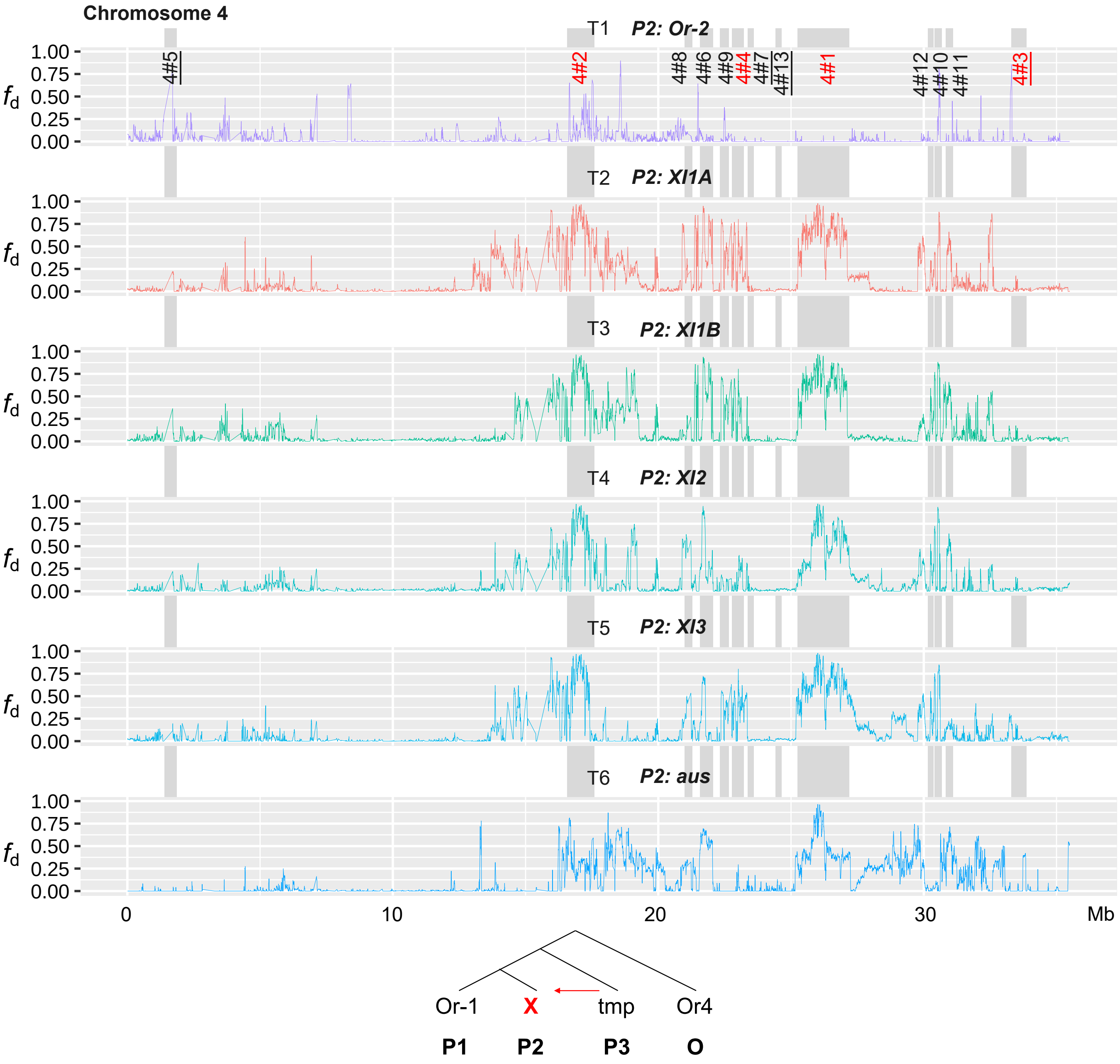


**Supplementary Fig. 18**

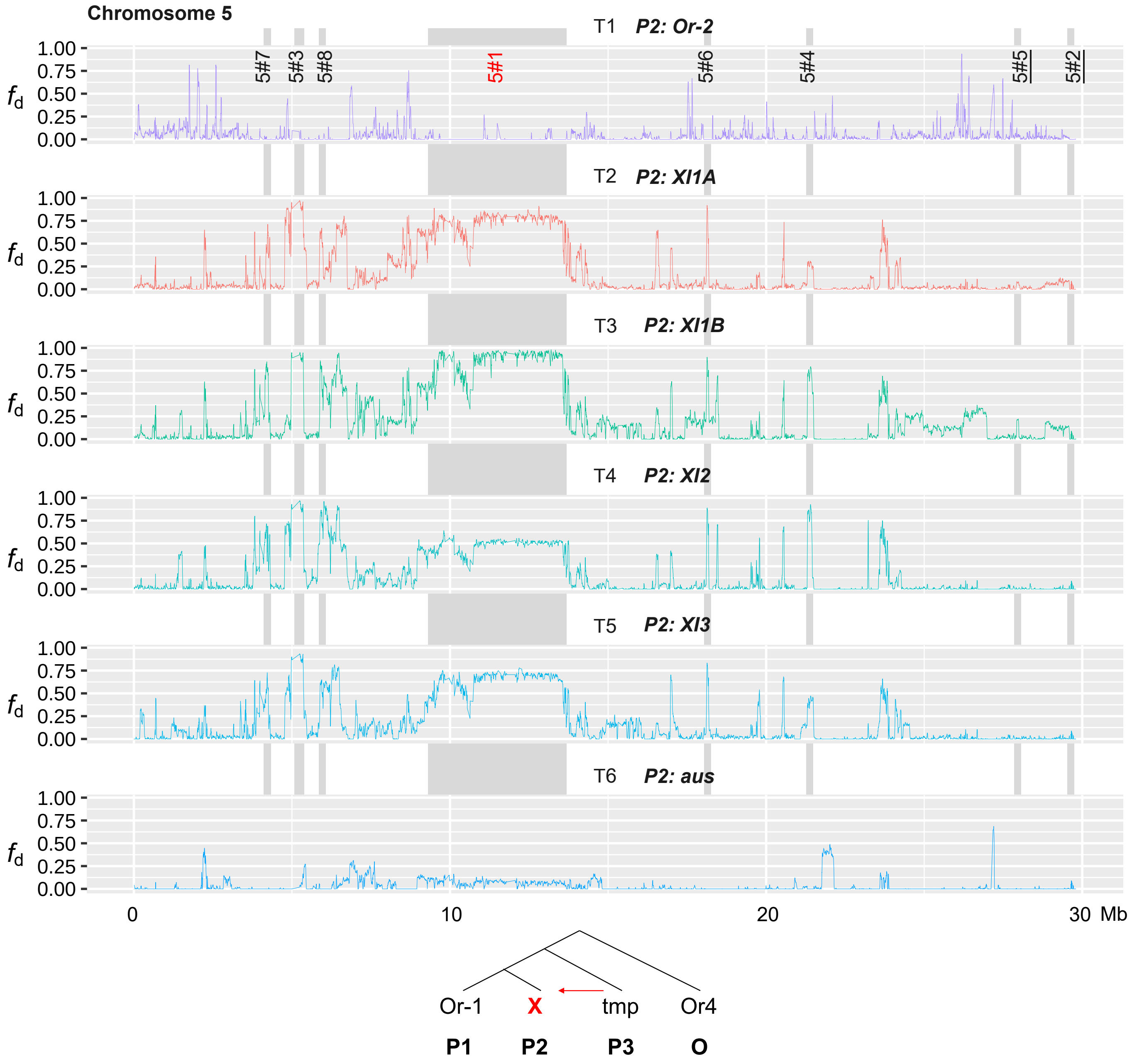


**Supplementary Fig. 18**

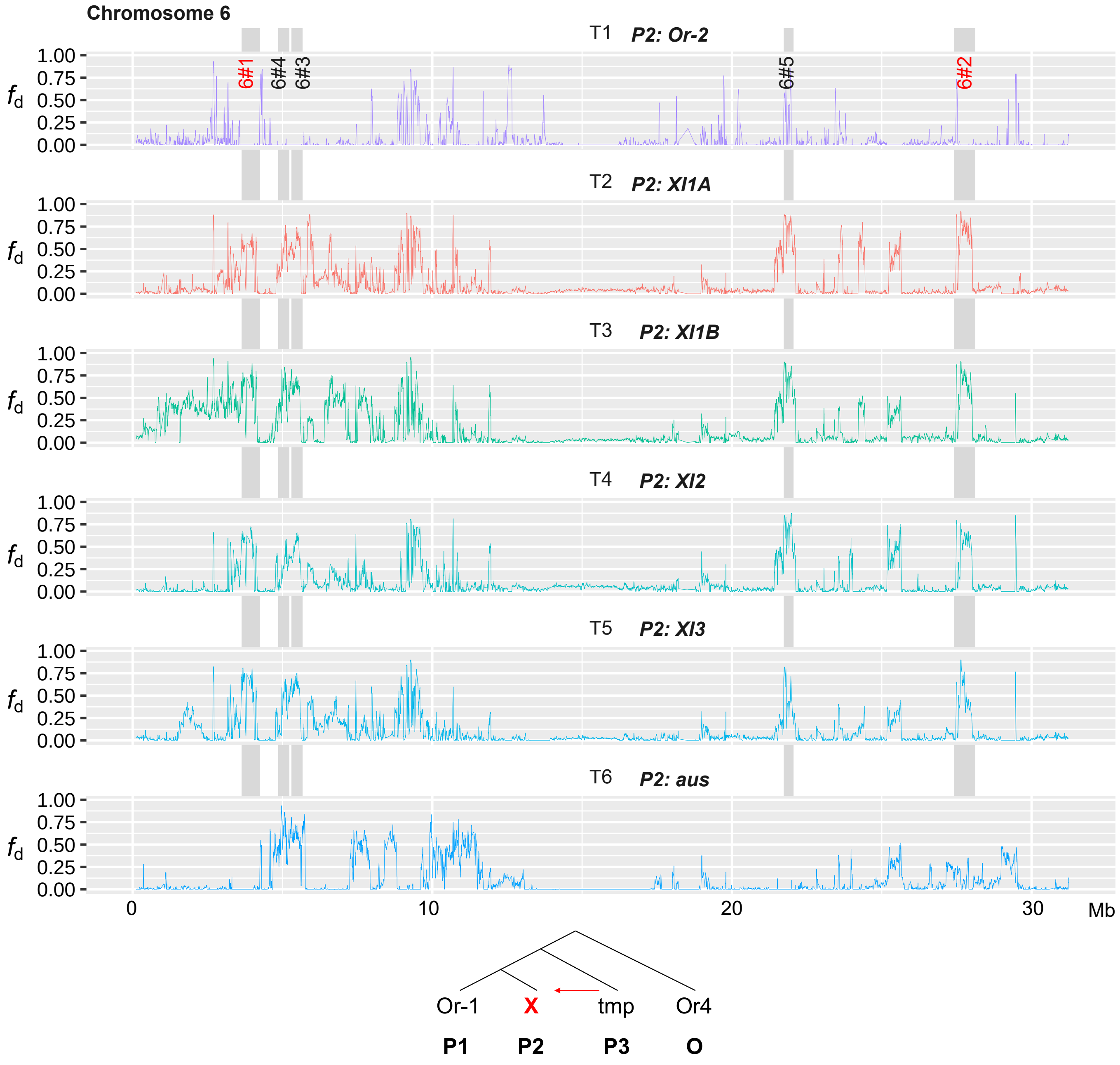


**Supplementary Fig. 18**

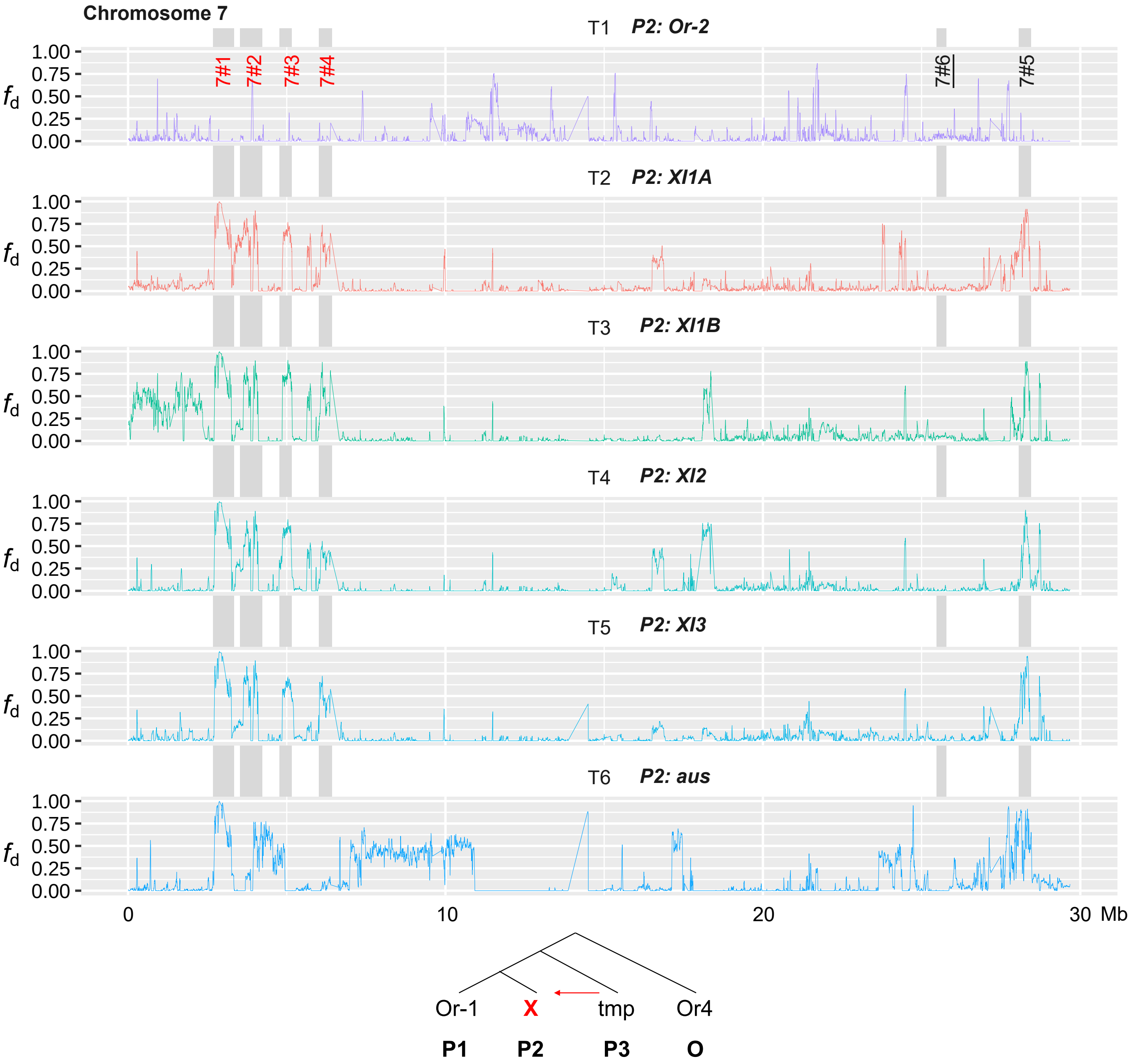


**Supplementary Fig. 18**

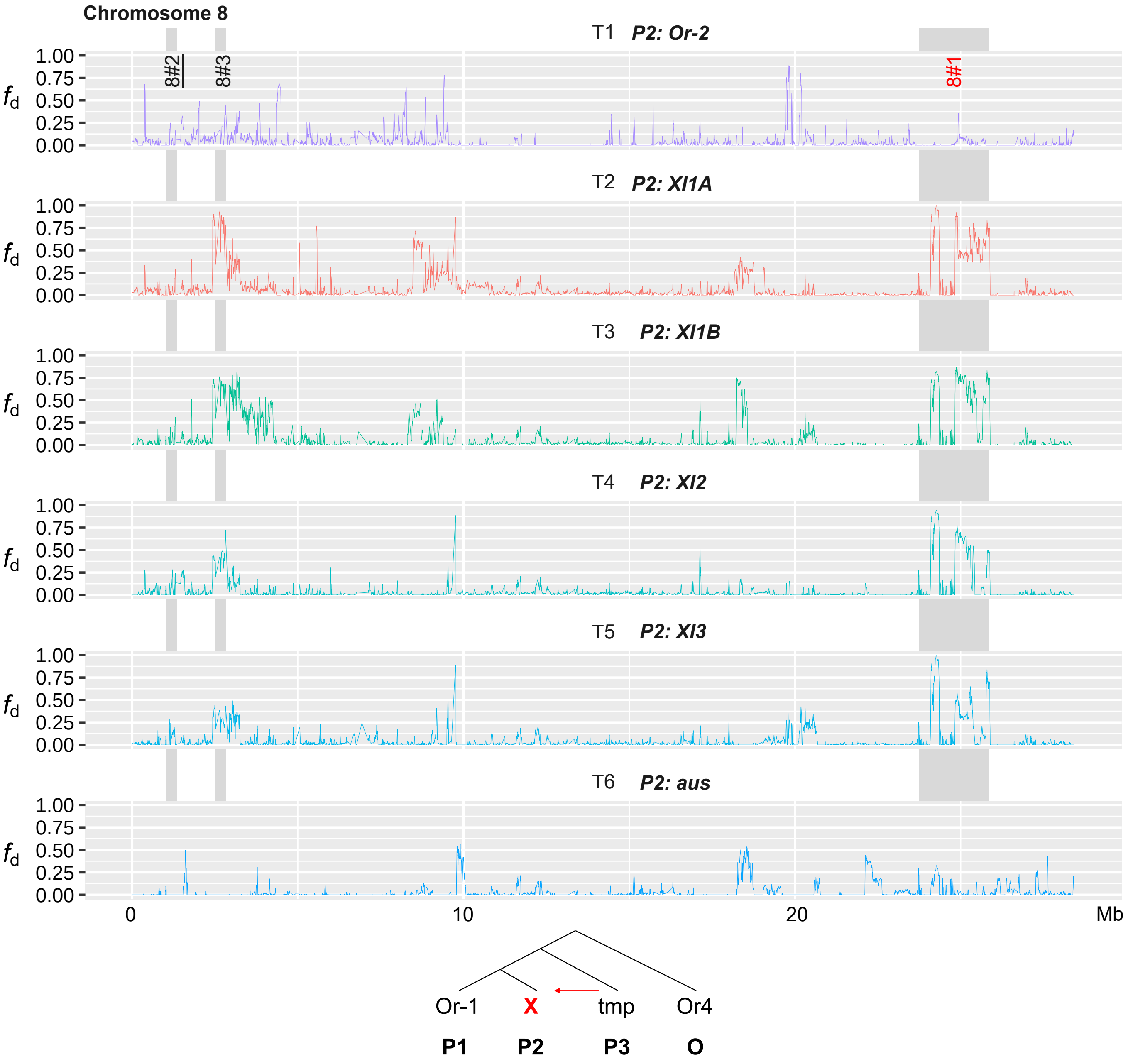


**Supplementary Fig. 18**

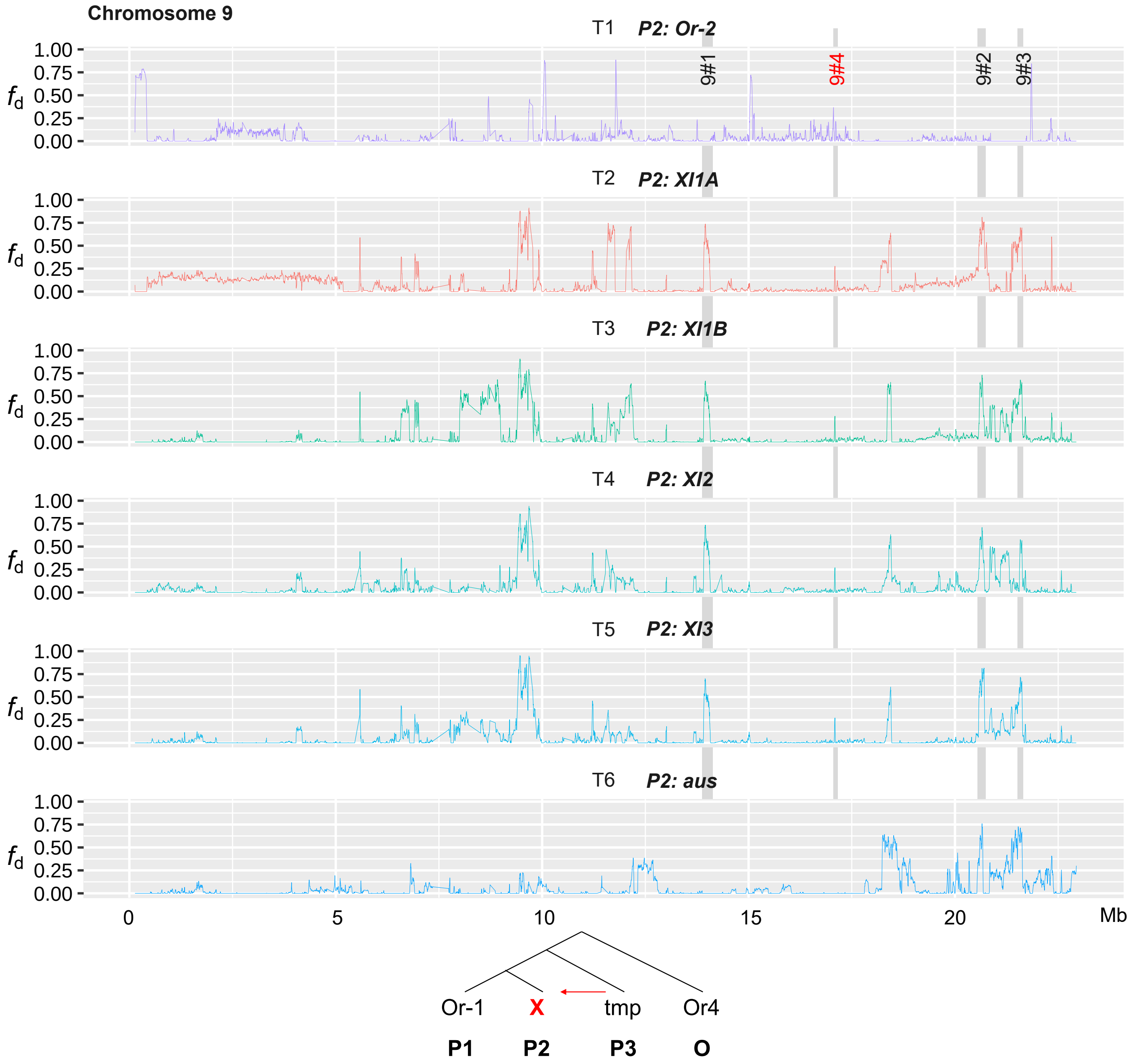


**Supplementary Fig. 18**

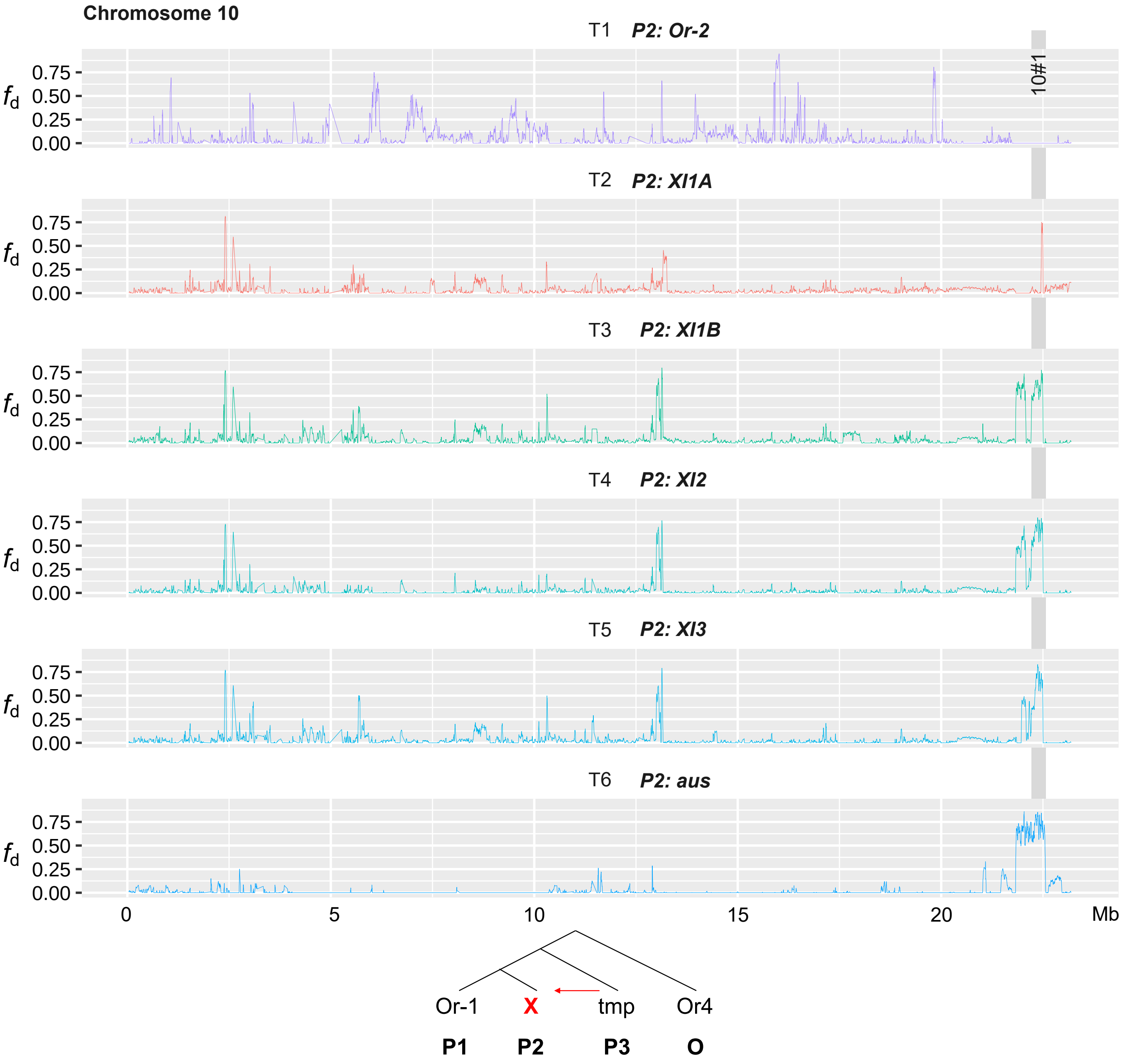


**Supplementary Fig. 18**

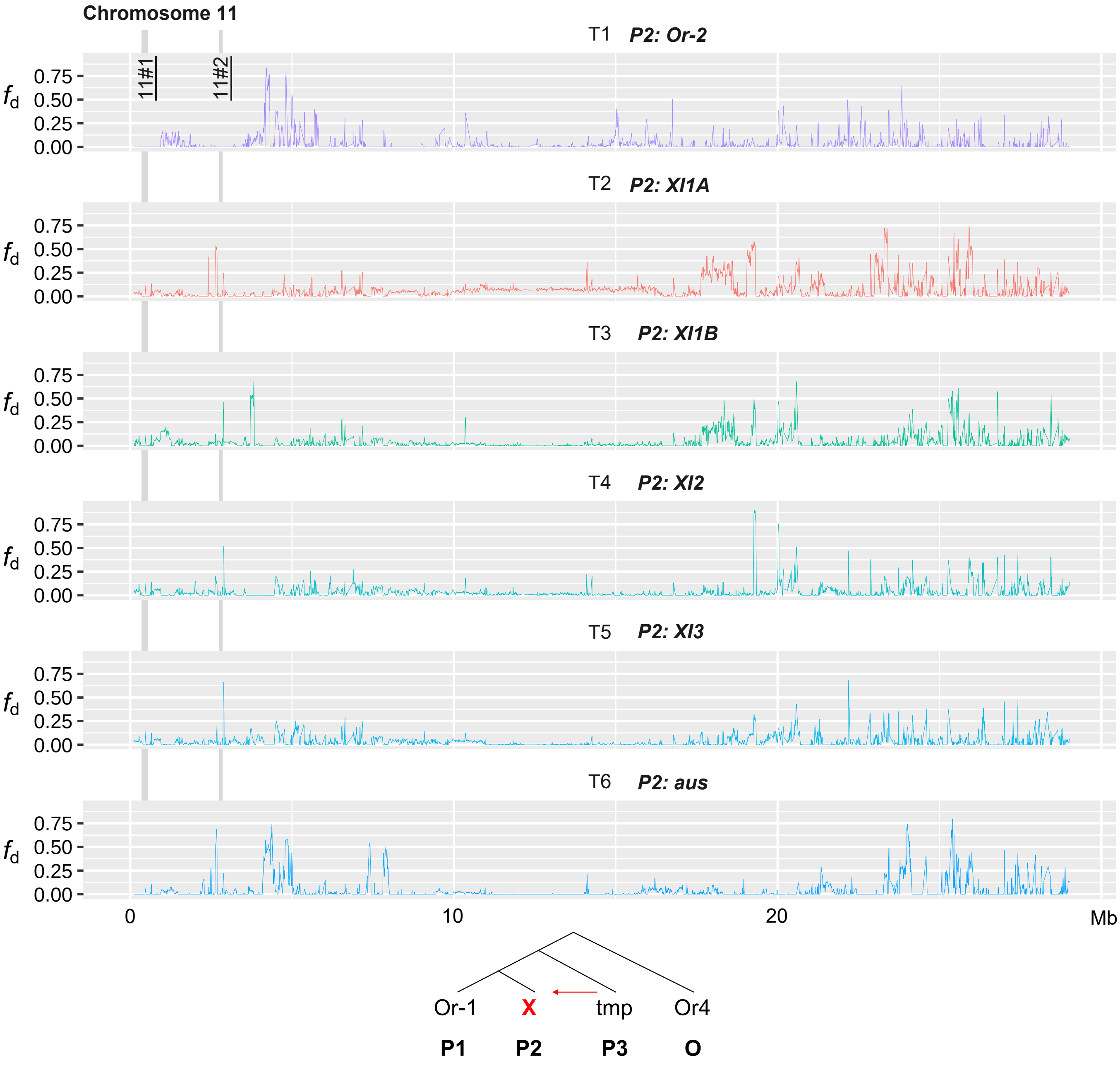


**Supplementary Fig. 18**

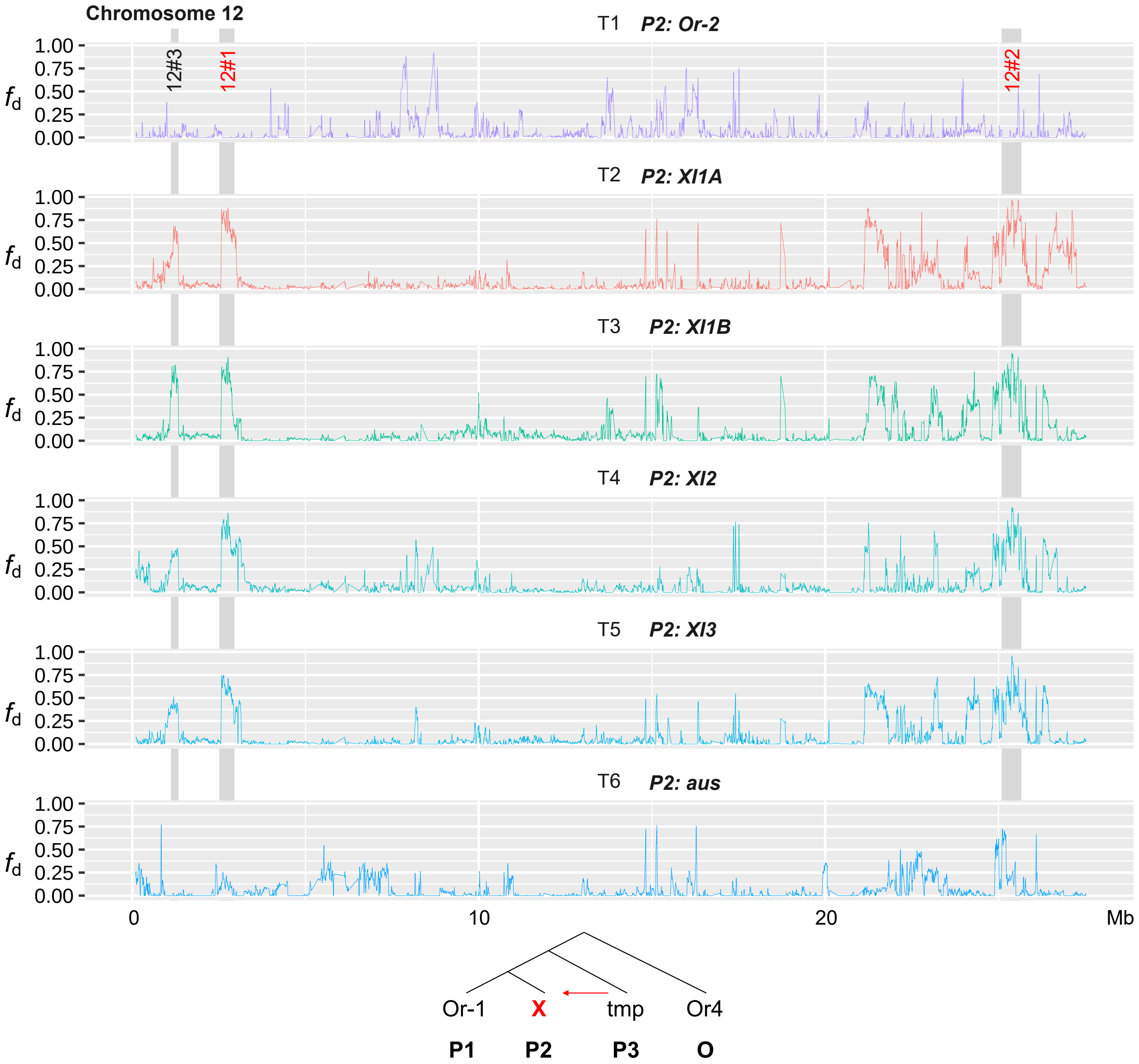


**Supplementary Fig. 18**

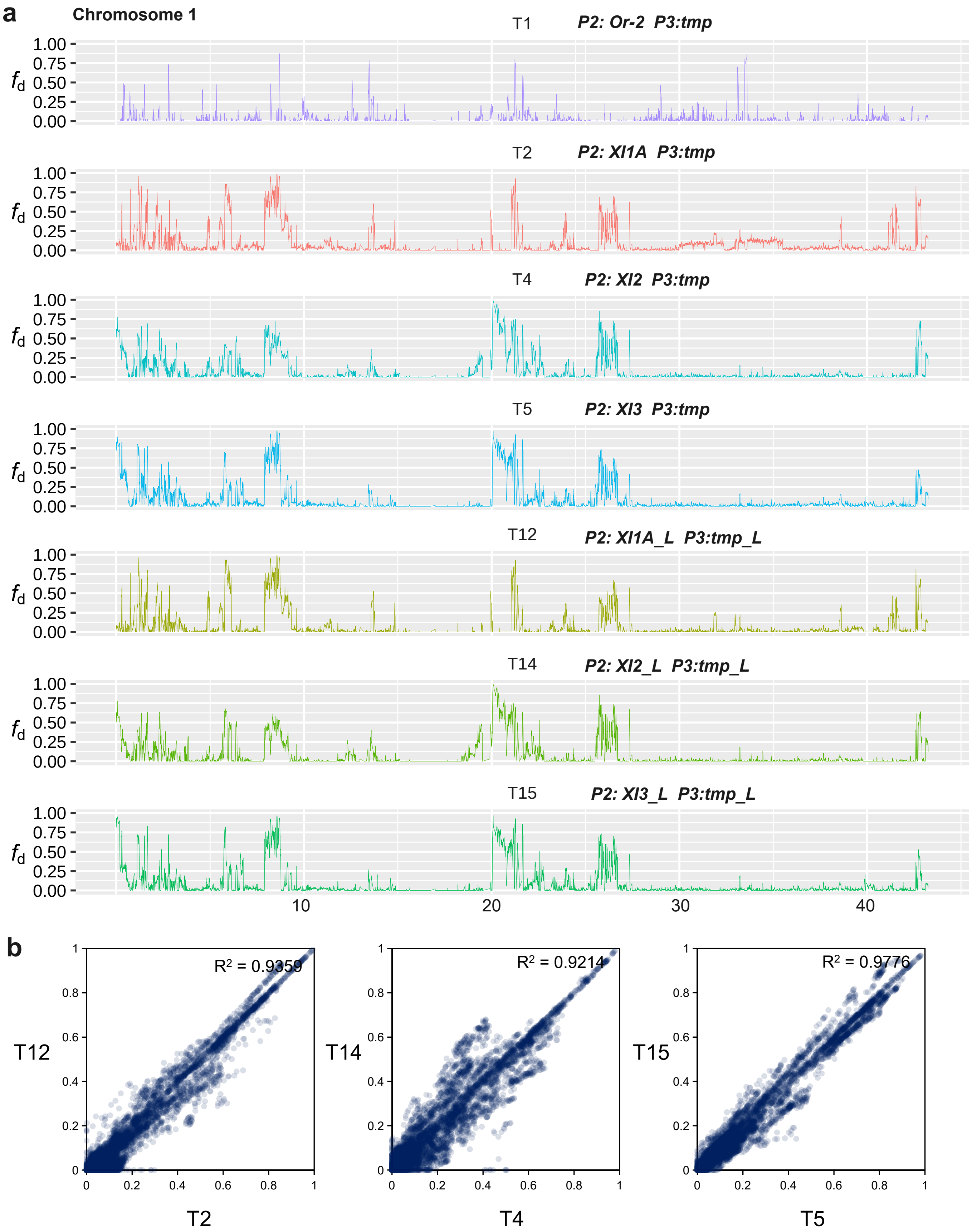

a
Chromosome 1
T1 P2: Or-2 P3:tmp
T2 P2: Xl1A P3:tmp
T4 P2: Xl2 P3:tmp
T5 P2: Xl3 P3:tmp
T12 P2: Xl1A_L P3:tmp_L
T14 P2: Xl2_L P3:tmp_L
T15 P2: Xl3_L P3:tmp_L
$f_d$
1.00
0.75
0.50
0.25
0.00
10
20
30
40
b
$R^2$ = 0.9359
$R^2$ = 0.9214
$R^2$ = 0.9776
T12
T14
T15
T2
T4
T5
0
0.2
0.4
0.6
0.8
1

**Supplementary Fig. 19** Comparison of $f_d$ using all genomes and landraces only under different topologies on chromosome 1. (**a**) $f_d$ distribution along chromosome 1. Group tmp, XI1A, XI2, XI3, and *aus* include 47, 24, 21, 52 and 32 landrace accessions, respectively. (**b**) Comparison of $f_d$ on chromosome 1 in T2 *vs* T12, T4 *vs* T14, and T5 *vs* T15.

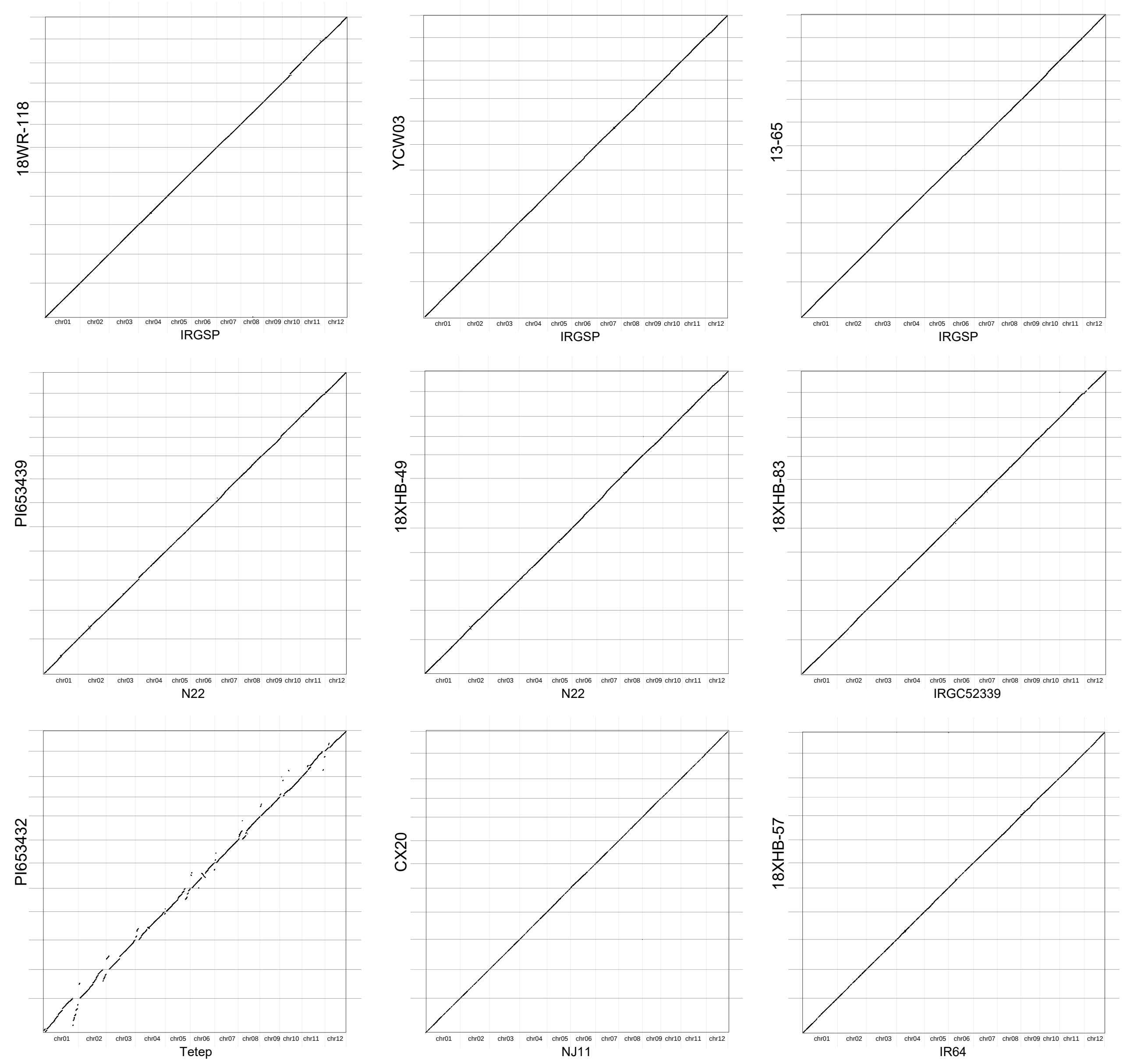


**Supplementary Fig. 20** Genome synteny between each weedy genome and corresponding phylogenetically closest cultivated rice assembly.

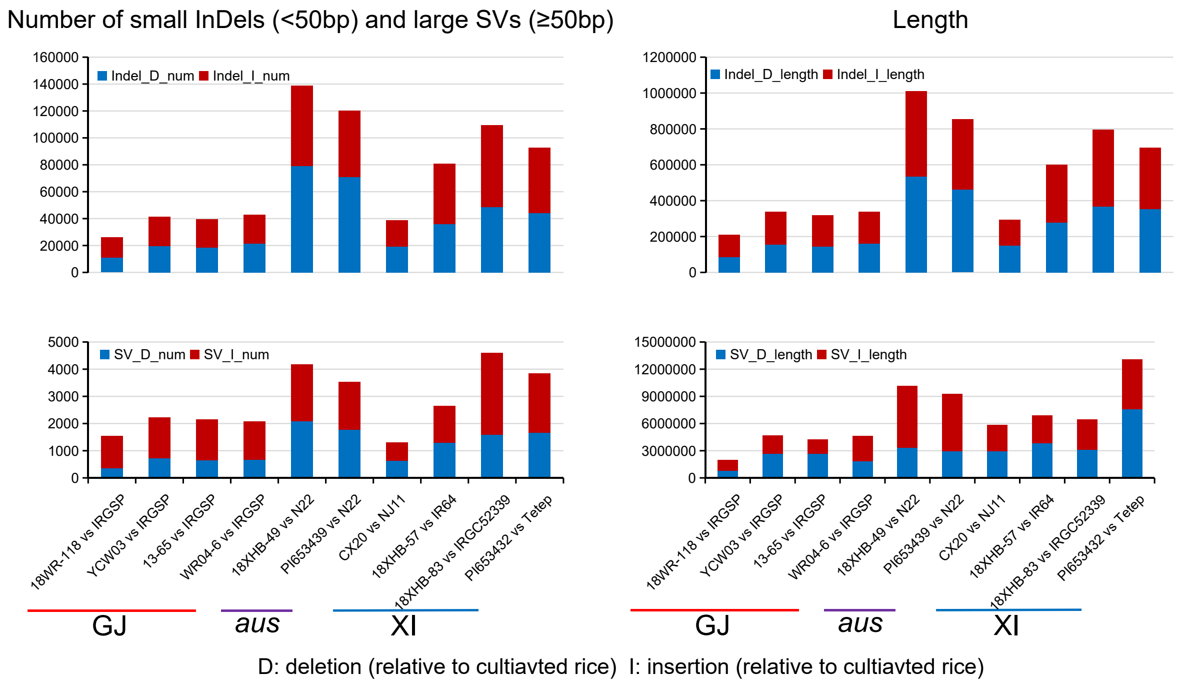


**Supplementary Fig. 21** Summary of structural variations between weedy and cultivated rice genomes.

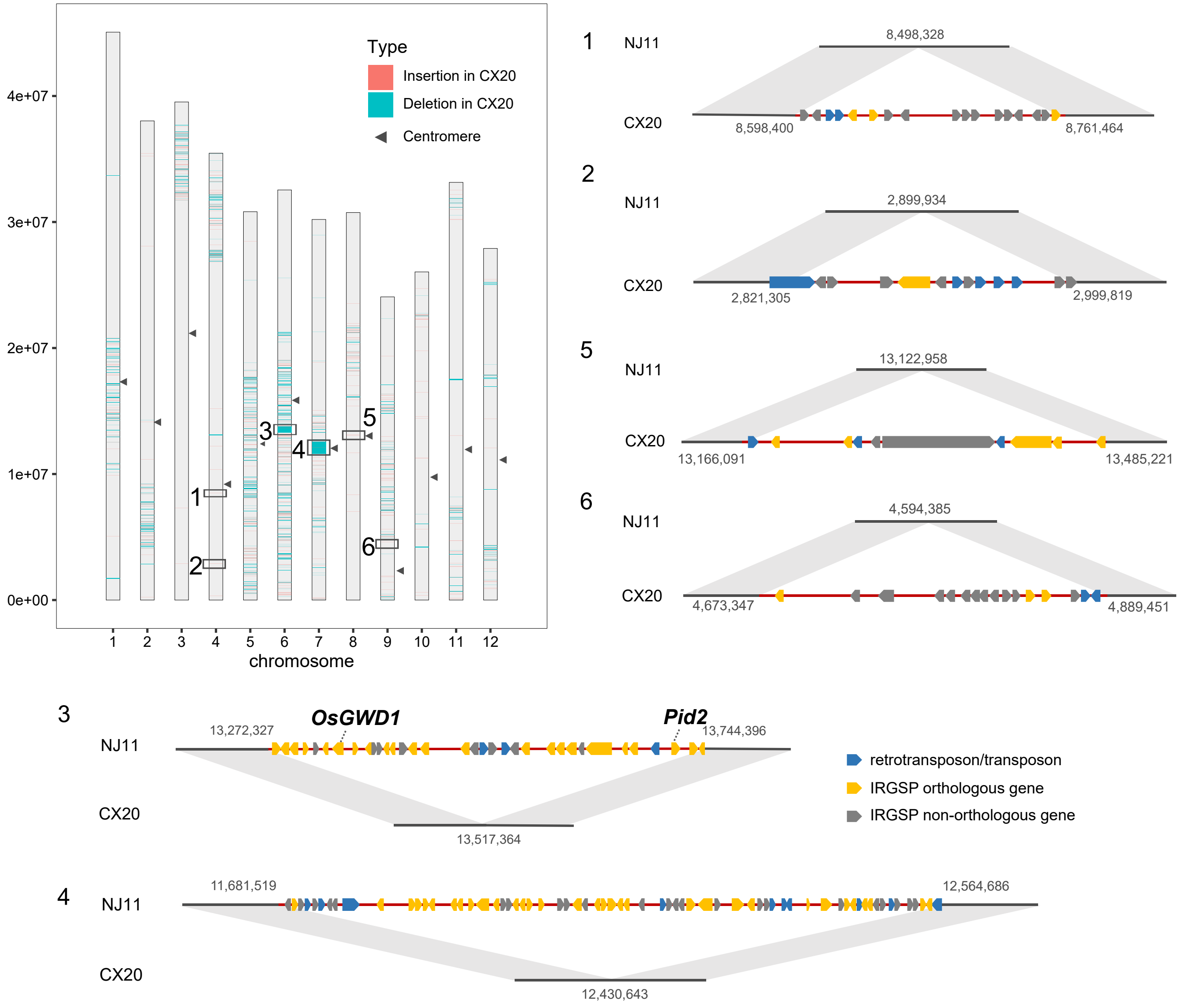


**Supplementary Fig. 22** Structural variations (>50bp) between assemblies of cultivar accession NJ11 and weedy rice accession CX20 on 12 chromosomes. The largest six SVs (numbered from 1 to 6) are zoomed in and annotated, including four insertions (1, 2, 5 and 6) and two deletions (3 and 4) in CX20.

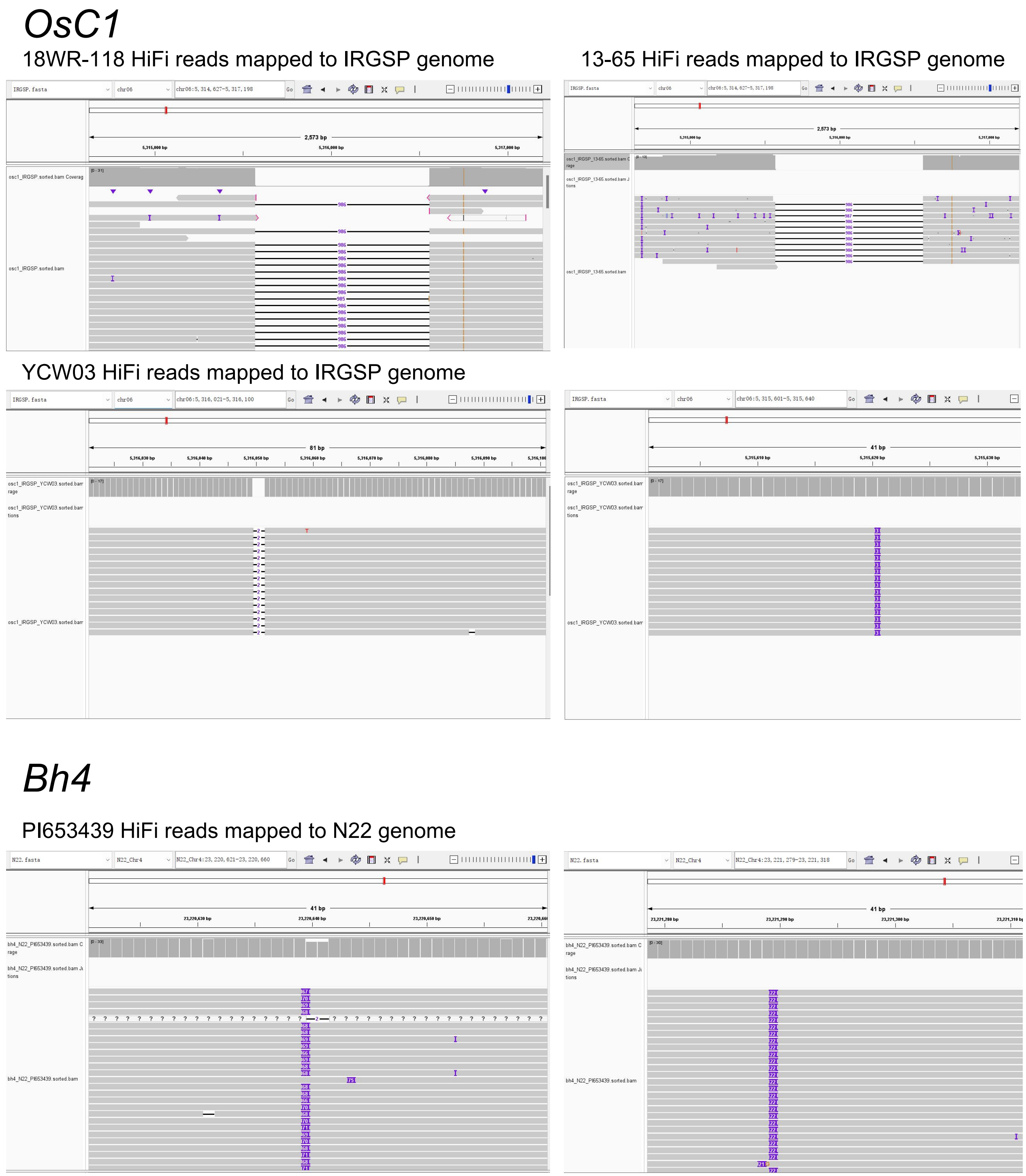


**Supplementary Fig. 23** The Integrative Genomics Viewer (IGV) snapshots show the structural variations in *OsC1* and *Bh4* between weedy and cultivated rice.

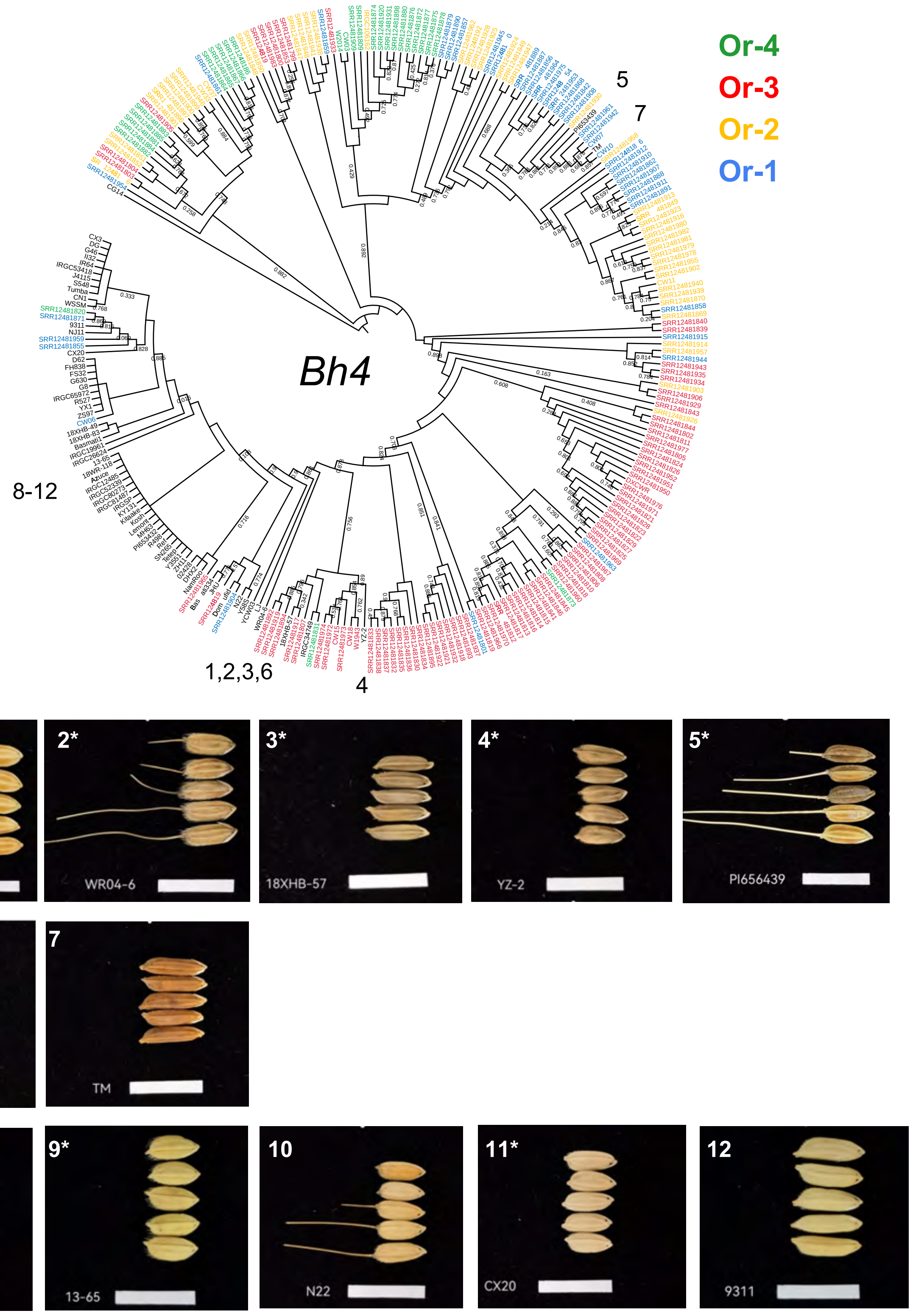


**Supplementary Fig. 24** ML phylogeny tree of *Bh4* in rice and morphology of rice seed hulls. Bootstrap values less than 0.90 are indicated on branches.
